# Metabolic arms race between a plant and a fungal pathogen

**DOI:** 10.1101/2023.07.29.551000

**Authors:** Y. Liu, L.K. Mahdi, A. Porzel, P. Stark, D. Esposto, A. Scherr-Henning, U. Bathe, I.F. Acosta, A. Zuccaro, G.U. Balcke, A. Tissier

**Affiliations:** Department of Cell and Metabolic Biology, Leibniz Institute of Plant Biochemistry, Halle, Germany; Institute for Plant Sciences, Cluster of Excellence on Plant Sciences (CEPLAS), Cologne Biocenter, University of Cologne, Cologne, Germany; Department of Bioorganic Chemistry, Leibniz Institute of Plant Biochemistry, Halle, Germany; Max-Planck Institute for Plant Breeding Research, Cologne, Germany

**Author notes:** Horticultural Sciences, University of Florida, Gainesville, USA.

## Abstract

In this work, we uncover a metabolite interaction between barley and the fungal pathogen *Bipolaris sorokiniana* (*Bs*), involving hordedanes, a previously undescribed set of labdane-related diterpenoids with antimicrobial properties. *Bs* infection of barley roots elicits hordedane synthesis from a 600-kb gene cluster. Heterologous reconstruction of the synthesis pathway in yeast produced several hordedanes, including one of the most advanced products 19-b-hydroxy-hordetrienoic acid (19-OH-HTA). Barley mutants in the diterpene synthase genes of the cluster are unable to produce hordedanes but, unexpectedly, show reduced *Bs* colonization. Accordingly, 19-OH-HTA enhances both germination and growth of *Bs*, while it inhibits other fungi, and *Bs* chemically modifies 19-OH-HTA. Thus, plant and pathogen molecular interactions extend beyond protein-protein recognition and the simple detoxification of plant antimicrobial metabolites.

**One-Sentence Summary:** **A fungal pathogen uses barley diterpenoid phytoalexins to facilitate root colonization**.

## Introduction

Plant pathogenic fungi impose a major burden on crop yield, and this impact is expected to increase with climate change (*1*). As the use of agrochemicals is increasingly under scrutiny by environmental agencies and single gene-for-gene resistance can be rapidly overcome in a changing climate, there is a strong need for more durable resistance traits. The fungus *Bipolaris sorokiniana* (*Bs*, teleomorph *Cochliobolus sativus*) is the pathogenic agent of spot blotch and root rot in wheat and barley, and is particularly prevalent in regions with warmer climates (*2*). Therefore, it represents a typical future important threat in the context of global warming. *Bs* can infect both aerial and underground parts of the plant but knowledge on how it interacts with roots is still limited (*3*).

There is ample evidence that infection of plants by microbial pathogens triggers the production of specialized metabolites that exhibit antimicrobial or antioxidant activities (*4*). These compounds, called phytoalexins, are not restricted to a particular chemical class but belong, for example, to phenylpropanoids, alkaloids or terpenoids (*4*). In barley, there is a number of reports of various phytoalexins produced in response to diverse pathogens. These include phenylamides, such as the dimeric hordatine A and B, the indole-derived gramine, benzoxazinones such as 2,4-dihydroxy-1,4-benzoxazin-3-one (DIBOA), methoxychalcones as well as tyramine and related amines (*5-7*). There is also extensive data on the nature and biosynthesis of a range of terpenoid phytoalexins in other important grass crops. Rice produces several classes of labdane-related diterpenoids, including momilactones (A and B) (*8, 9*), phytocassanes, oryzalexins (*10-14*) and oryzalides (*15, 16*), and the macrocyclic *ent-*oxodepressin (*17*). Maize produces dolabralexins and kauralexins, both labdane-related diterpenoid phytoalexins (*18*), as well as the sesquiterpenoid zealexins (*19*). In contrast, no sesqui-or diterpenoid phytoalexins have been identified in barley yet.

Here, we characterize a barley gene cluster responsible for the production of a set of diterpenoid phytoalexins and investigate their role during the interaction with *Bs*. Unexpectedly, we find that colonization by *Bs* of gene-edited barley mutants that do not produce these diterpenoids is less effective than that of wild-type (WT) plants. In agreement of this colonization phenotype, *Bs* grows faster in the presence of a major barley diterpenoid and displays the ability to modify it, providing an explanation for the enhanced colonization when these phytoalexins are produced.

## Results

### A barley gene cluster for diterpenoid phytoalexins

In previous work, we identified a set of genes induced upon infection with *Bs* potentially involved in diterpenoid biosynthesis, including homologs of a copalyl diphosphate synthase (*HvCPS2*), a kaurene synthase-like (*HvKSL4*) and several cytochrome P450 oxygenases (CYPs) (*3*). These genes are clustered on chromosome 2 in a region spanning around 600 kb (**Fig. 1, Table S1**). A phylogenetic analysis of the encoded proteins suggests that HvCPS2 is a (*+*)-copalyl diphosphate synthase and shows that the CYPs belong to two families, respectively CYP99A and CYP89E (**Fig. 1, Fig. S1-S2, Table S2-S4**). Notably, CYP99As are also found in the momilactone gene cluster in rice (*20*). Most genes in the barley cluster show strong transcriptional induction upon infection by *Bs* (**Fig. 1**). Analysis of extracts from roots and root exudates by GC-MS and LC-MS/MS indicates the presence of several compounds with masses and fragmentation patterns indicative of diterpenoids (**Fig. 1, Fig. S3**), suggesting that they may be products of the chromosome 2 cluster.

**Fig. 1:**
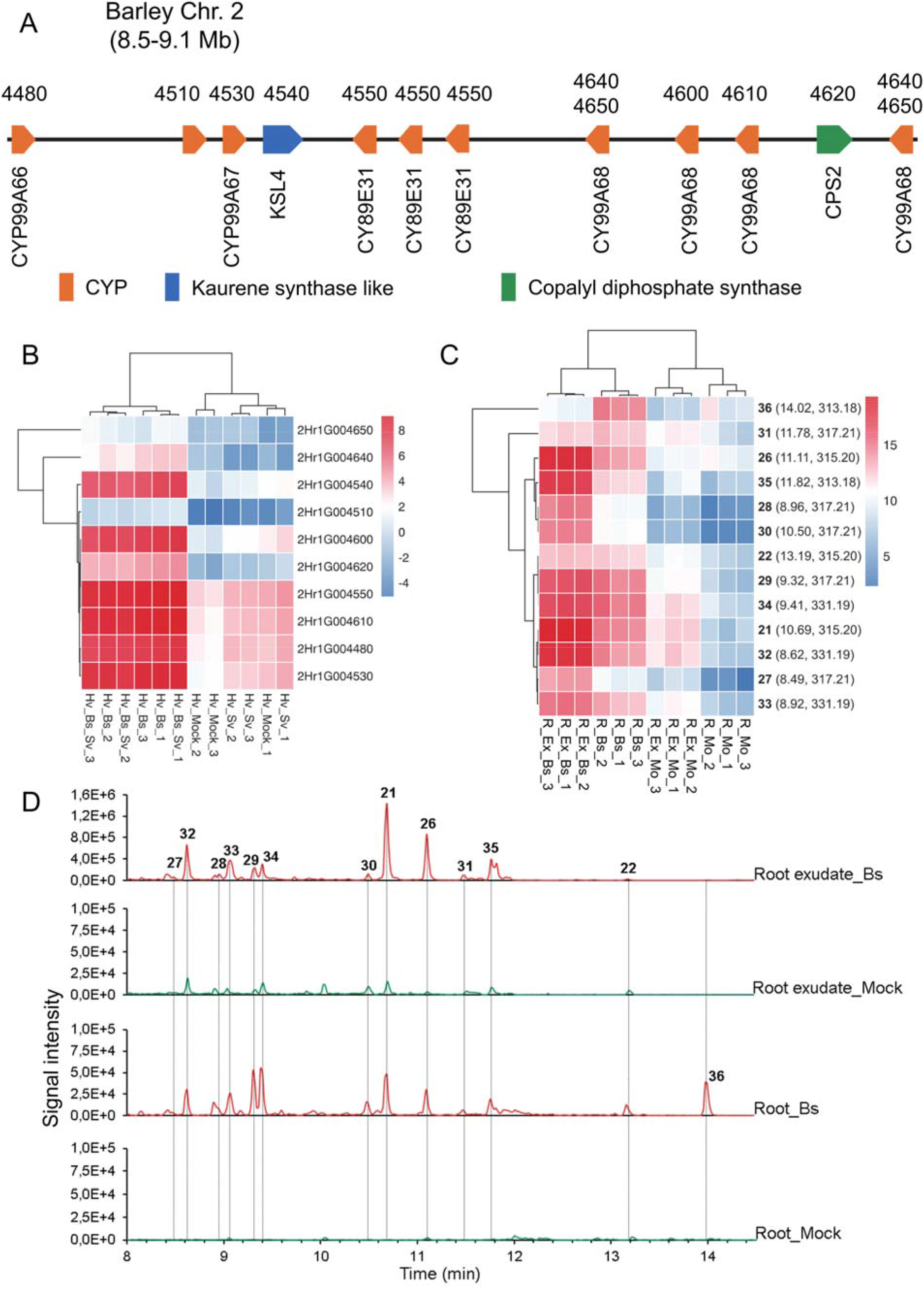
Pathogen-inducible expression in a barley gene cluster for diterpenoid biosynthesis and production of diterpenoids by barley roots upon pathogen infection. **A)** Overview of the chromosome 2 biosynthetic gene cluster. Three versions of the barley reference genome, including Morex V1, V2 and V3 (*35-37*) were used to generate the gene annotation. The number on the top of genes indicates the last 4 digits of the gene ID according to Morex V1, and the given name of the genes is provided below. CYP: cytochrome P450 oxygenase; Mb: million base pairs. **B)** Differential expression of genes from the cluster upon infection of barley roots by pathogenic and beneficial endophytic fungi. The colors represent gene expression values as log2-transformed FPKM values. The raw data are from (*3*). Hv_Mock, barley roots mock inoculated; Hv_Sv; barley roots inoculated with *Serendipita vermifera* (*Sv*); Hv_Bs, barley roots inoculated with *Bipolaris sorokiniana* (*Bs*); Hv_Bs_Sv, barley root co-inoculated with *Bs* and *Sv*. **C)** Diterpenoids identified from barley roots or root exudates after *Bs* infection and compared with mock-treated samples. R_Bs, roots inoculated with *Bs*; R_Mo, roots mock inoculated; R_Ex_Bs, exudates of roots inoculated with Bs; R_Ex_Mo, Exudates of roots mock inoculated. The digits at the end of the sample names indicate independent biological replicates. The color scale bar represents log2-transformed peak intensity values measured by UPLC-QToF. The identity of each diterpenoid is indicated by a number in bold, followed by its retention time and mass to charge ratio. **D)** LC-HRMS (negative mode) chromatograms of diterpenoids from barley roots or root exudates after *Bs* infection or mock treatment. Selected ions, m/z 313.2, 315.2, 317.2 and 331.2.

### Heterologous expression identifies a set of barley diterpenoids of the cleistanthane group

Using previously established expression systems in yeast (*Saccharomyces cerevisiae*) and *Nicotiana benthamiana* (*21, 22*) as well as *in vitro* assays, we reconstituted major parts of a network of diterpenoids and characterized several of the intermediates and products by NMR. As reported previously by our group and others (*23, 24*), the diterpene backbone (compound **1**) produced by the combined action of HvCPS2 and HvKSL4 stems from (+)-copalyl diphosphate and belongs to the cleistanthane group, with two double bonds in the C-ring at positions C8-C9 and C12-C13, a methyl group attached to C13 and an ethyl group attached to C14 in the α-configuration (**Fig. S4, Fig. S9, Fig. S13, Tab. S1**). We named this diterpene olefin hordediene and the group of barley diterpenoids derived from it hordedanes. We then expressed the four most abundant CYPs from the cluster (CYP89E31, CYP99A66, CYP99A67, CYP99A68) with HvCPS2 and HvKSL4 either individually or in combinations of 2 to 4 CYPs. CYP89E31 carries out aromatization of the C-ring (compound **5**) and C11-hydroxylation (compound **8**), as described in our earlier preprint (*23*) (**Fig. 2, Fig. S5-S6, Fig. S13, Tab. S6-S7**). Both **5** and **8** are present in barley root extracts elicited by *Bs* infection (**Fig. S3**). CYP99A66 and CYP99A67 have partially overlapping activities with both of them able to oxidize the C17-methyl group multiple times and only CYP99A67 able to oxidize the C19-methyl group (**Fig. 2, Fig. S5, Fig. S7, Fig. S13**). CYP99A68 oxidizes compound **8** multiple times on C16, leading to compound **22**, which is present in *Bs* infected roots and root exudates (**Fig. 2C, Fig. S8, Fig. S13**). In total, we detected at least 17 hordedanes in barley roots of which 8 were produced in yeast (**Fig. 2**).

**Fig. 2:**
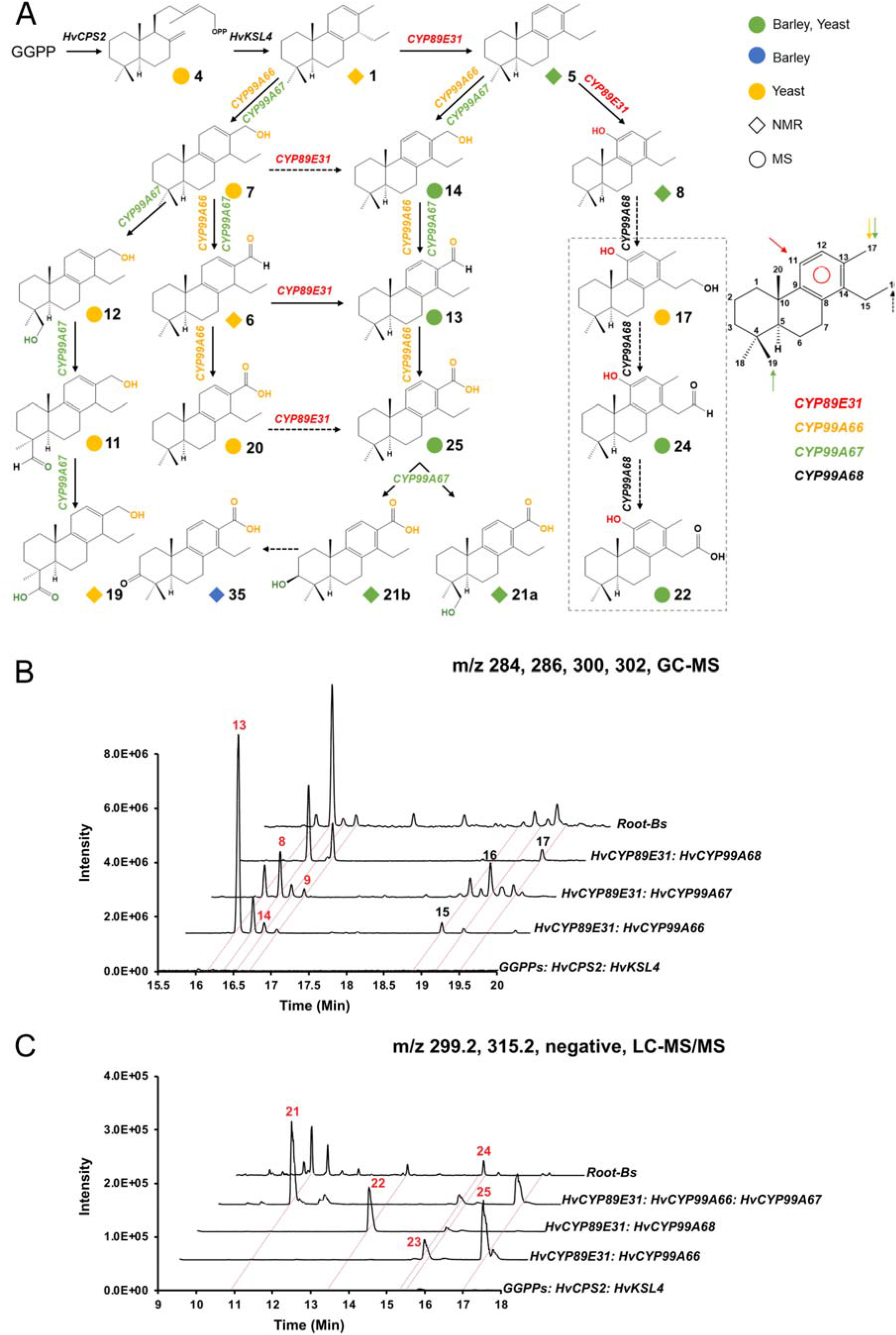
Elucidation of the pathway for biosynthesis of barley diterpenoids. **A)** Biosynthesis pathway of barley diterpenoids by heterologous expression in yeast and *N. benthamiana*. The diamond and the circle symbols indicate diterpenoids characterized by NMR and MS, respectively. Green symbols indicate diterpenoids identified in barley, yeast and *N. benthamiana*. Blue symbols indicate diterpenoids identified in barley only; Orange symbols indicate diterpenoids identified only in heterologous expression. GC-MS (**B**) or LC-MS/MS (**C**) analysis of diterpenoid products from heterologous expression in yeast of two or three *CYPs. CYPs* were always co-expressed together with *HvCPS2, HvKSL4* and *GGPPs*. Peaks that are present in barley are indicated by red numbers.

Importantly, the combination of CYP89E31, CYP99A66 and CYP99A67 with HvCPS2 and HvKSL4 results in the production of compounds **21a** and **21b**, respectively 19-β-hydroxy-hordetrien-17-oic acid, (19-OH-HTA) and 3-β-hydroxy-hordetrien-17-oic acid (3-OH-HTA), which combined represent the most abundant peak (**21**) in barley root exudates, albeit in a different ratio in yeast and barley (**Fig. 2C, Fig. S10, Fig. S13, Tab. S10**). These results show that we could reconstitute to a large extent the pathway for the barley diterpenoid phytoalexins in yeast.

### Hordedane diterpenoids are absent in *cps2* and *ksl4* mutants

To evaluate the contribution of *HvCPS2* and *HvKSL4* to the barley diterpenoids and to study their biological roles in the interaction with *Bs*, we performed targeted mutagenesis of *HvCPS2* and *HvKSL4* by CRISPR/Cas9 gene editing on the barley cultivar Golden Promise Fast hereafter referred to as wild-type (WT) (*25*). Two sgRNAs for each gene were used and two independent homozygous mutant lines for both *HvCPS2* and *HvKSL4 (cps2-1, cps2-2, ksl4-1, ksl4-2)* were selected for further characterization (**Fig. S11**). All mutations lead to a premature stop codon and the synthesis of truncated proteins missing essential residues for catalysis (**Fig. S10B-C**).

Furthermore, the expression of the mutated genes is strongly reduced (**Fig. S12**). We performed an infection assay with *Bs* on the four mutants and WT barley and analyzed roots and root exudates by untargeted metabolomic profiling. The induced diterpenoids in WT plants are absent in all four mutants (**Fig. 3**). These results show that these diterpenoids derive from the products of the two diterpene synthases encoded by *HvCPS2* and *HvKSL4* in the chromosome 2 gene cluster.

**Fig. 3:**
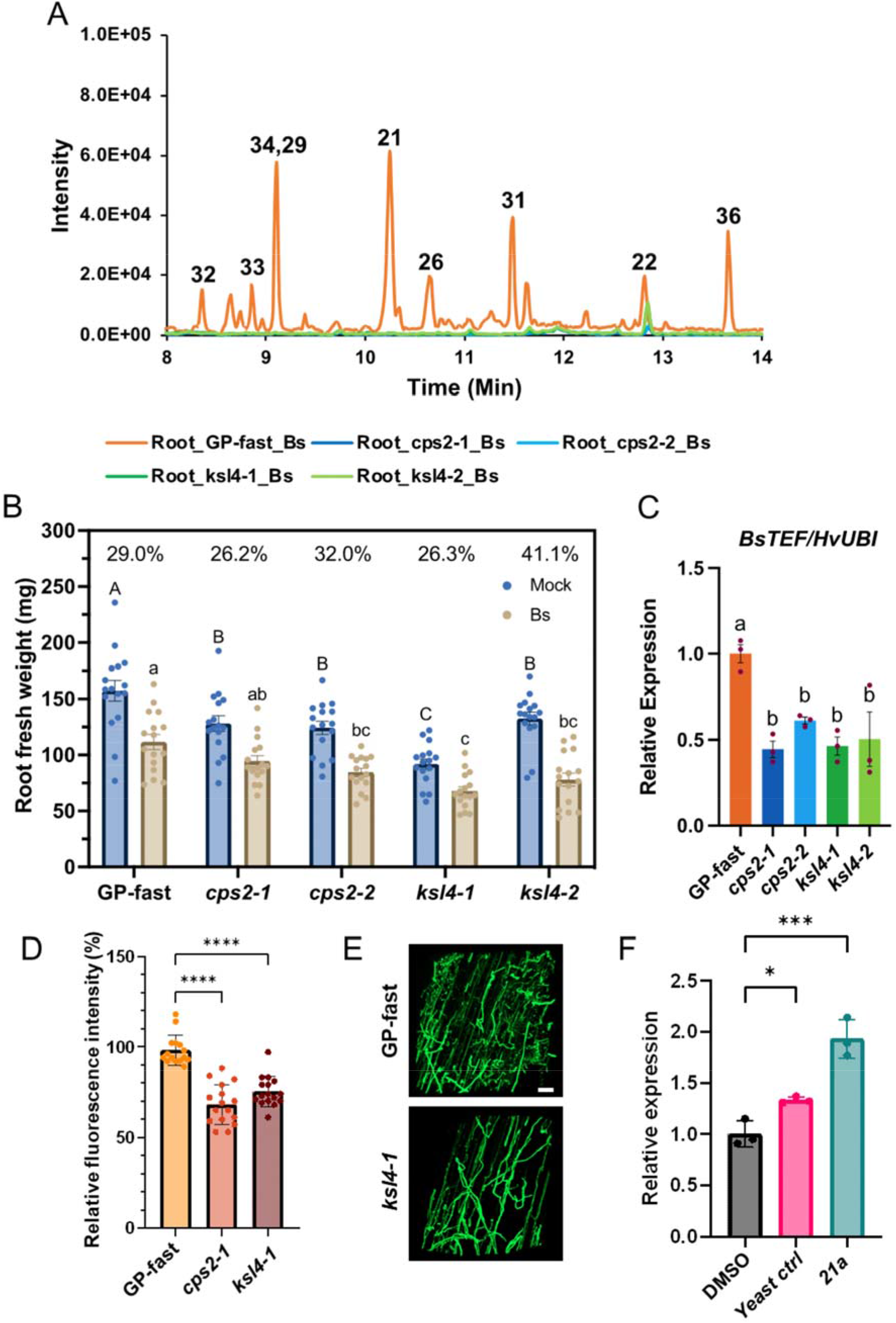
Characterization of *cps2* and *ksl4* barley mutants. **A)** Chromatograms (LC-MS/MS, negative mode) of diterpenoids in wild-type barley, *cps2* and *ksl4* mutants after *Bs* infection. Selected ions: m/z 313.2, 315.2, 317.2, 331.2. **B)** Fresh weight of barley roots 6 dpi with *Bs* or mock-inoculation. Bars: mean ± standard error of mean (SEM) (n = 16). The percentages represent the reduction of biomass for each genotype of barley after *Bs* infection compared with corresponding controls. It calculated by dividing the mean difference of root weight between mock- and *Bs*-inoculation by the mean root weight of mock-inoculation. The biomass reduction upon infection between lines was not significant (two-way ANOVA and Tukey’s post-hoc tests, p = 0.1197). Letters (uppercase letters for samples with mock-treatment, and lowercase letters for samples inoculated with *Bs*) represent statistically significant differences according to Tukey’s post-hoc test. Bars: mean ± SEM (n = 16). **C)** Quantification of *Bs* in the roots by qRT-PCR. The reference gene of barley is *Ubiquitin. BsTEF*: *Bipolaris sorokiniana* translation elongation factor. Statistics: one-way ANOVA and Tukey’s post-hoc test, p = 0.0039. Bars: Mean ± SEM (n = 3). **D)** Fluorescence intensity of barley roots infected by Bs and stained with WGA-AlexaFluor 488. **E)** Representative images of root segments of wild type (GP-fast) and *ksl4-1* mutant stained with WGA-AlexaFluor 488. The white bar indicates 100 µm in both images. **F)** Quantification of Bs in roots by qRT-PCR as in C). Barley *ksl4-1* mutants were inoculated by Bs in the presence of DMSO, purifed 19-OH-HTA from yeast (21a), or from a yeast strain with empty vector processed the same way as the strain producing 19-OH-HTA.

### Reduced root colonization of *cps2* and *ksl4* mutants by *B. sorokiniana*

Next, we compared the performance of hordedane-less barley mutants with WT plants upon infection by *Bs*. We measured the fresh weight of roots at 6 dpi for infected and non-infected plants. Infection by *Bs* leads to growth reduction in all genotypes but we observed no significant difference in the extent of growth reduction of roots between WT and *cps2* or *ksl4* mutants (two-way ANOVA, P value=0.1197) (**Fig. 3B**). We then quantified the relative amount of *Bs* in the roots of all genotypes by qRT-PCR and, unexpectedly, detected less *Bs* in all four mutants compared with the WT (**Fig. 3C)**. The reduced *Bs* colonization was confirmed by measuring the fluorescence after staining with a fungal-specific fluorescent marker (**Fig. 3D-E**) These results suggest that in the context of the interaction between the pathogenic *Bs* and barley roots, the presence of the induced barley diterpenoids increases the ability of *Bs* to colonize barley roots. We therefore sought to investigate the effect of the diterpenoids on *Bs*.

### 19-OH-HTA reverses the reduced colonization of diterpenoid-free mutants

We tested the ability of 19-OH-HTA to reverse the reduced colonization of *ksl4* mutants. For this, we added 19-OH-HTA at a concentration of 50 µM to the agar medium used for the colonization assays and compared it to a mock addition or to an extract from the background yeast strain, i.e. processed in the same way as for the purification of 19-OH-HTA. Quantification of *Bs* abundance qRT-PCR shows that colonization levels were significantly increased in the presence of 19-OH-HTA (**Fig. 3F**).

### 19-OH-HTA enhances the growth of *Bs* and reverses the reduced colonization of diterpenoid-free mutants

We took advantage of the availability of a yeast strain producing 19-OH-HTA and small amounts of 3-OH-HTA to purify the mixture and use it for bioassays. Because 19-OH-HTA is the major compound of this mixture, we refer to it as 19-OH-HTA for simplification. We tested 19-OH-HTA on five fungi, including three phytopathogens, *Bs, Verticilium dahliae, Fusarium culmorum*, and two beneficial fungal endophytes, *Serendipita indica* and *S. vermifera*. 19-OH-HTA strongly inhibited the spore germination of *F. culmorum* (5.2% spore germination in the presence of 19-OH-HTA versus 93.7% without it), and reduced its hyphal growth significantly (**Fig. 4**). Similar inhibitory effects were also observed on the two beneficial fungal endophytes, indicating a broad activity spectrum, a feature frequently reported for other phytoalexins (*18, 19*). No effect was observed on the germination and growth of *V. dahliae*. By contrast, 19-OH-HTA enhanced the spore germination of *Bs* (**Fig. 4B**). Furthermore, incubation with 19-OH-HTA at 25-50 µM concentrations leads to significantly enhanced growth of *Bs* in a concentration-dependent manner (**Fig. 4C**).

**Fig. 4:**
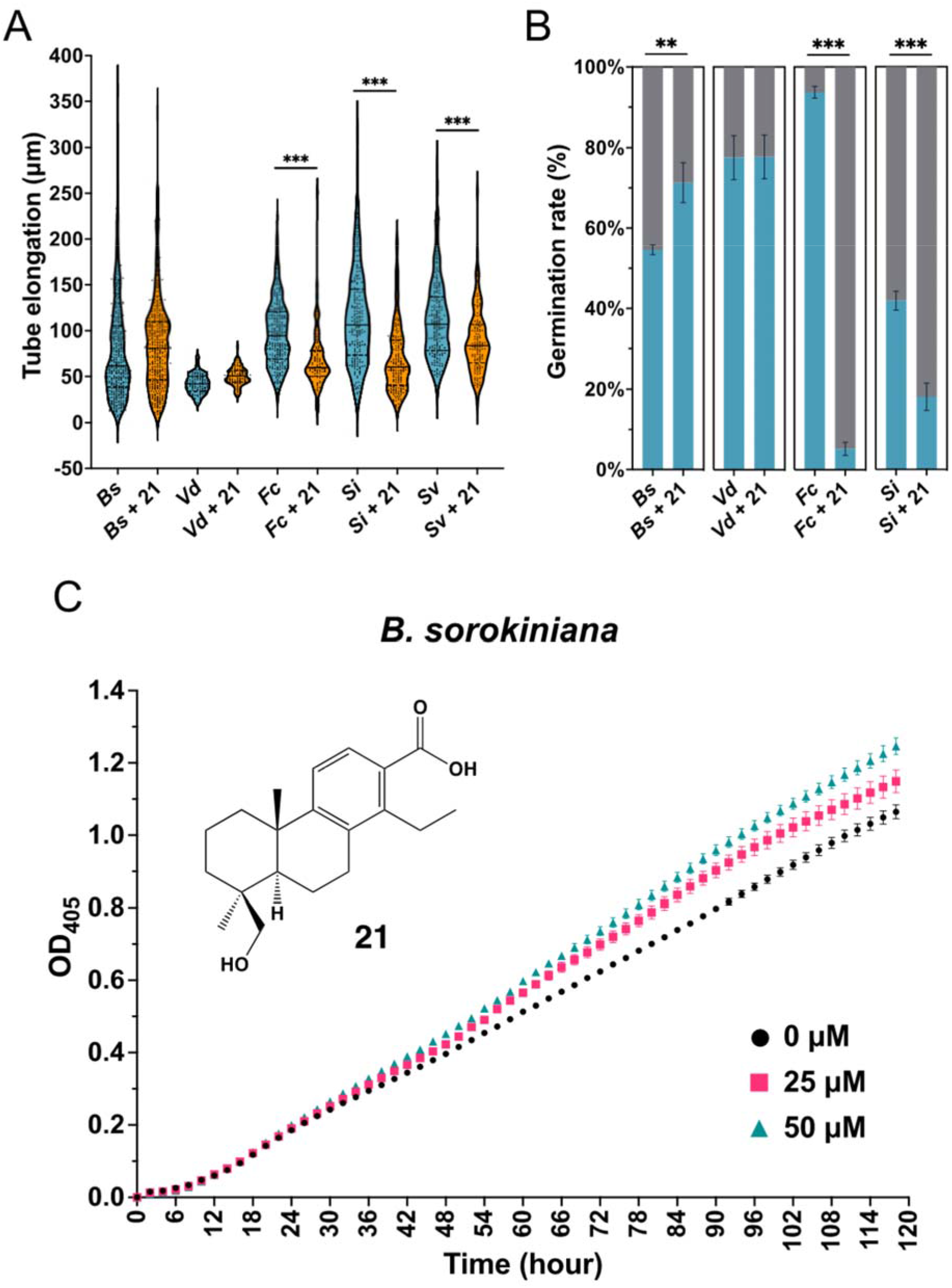
Effect of 19-OH-HTA application on different fungi. **A)** Tube elongation/hyphal length of *Bipolaris sorokiniana* (*Bs*, 3 hpi), *Verticilium dahlia* (*Vd*, 9 hpi), *Fusarium culmorum* (*Fc*, 9 hpi), *Serendipita indica* (*Si*, 12 hpi) and *Serendipita vermifera* (*Sv*, 12 hpi) after application of 0 or 50 µM 19-OH-HTA to the growth media. **B)** Percentage of germinated (blue) and not germinated (grey) spores of *Bs* (3 hpi), *Vd* (9 hpi), *Fc* (9 hpi) and *Si* (12 hpi) after application of 0 or 50 µM 19-OH-HTA to the growth media. The incubation time depended on the speed of fungal spore germination. Bars: Mean ± SD (n = 3). **C)** *Bs* growth time course in media containing 19-OH-HTA at the concentration of 0, 25 and 50 µM respectively. Bars: Mean ± SEM (n = 8-9). Asterisks indicate significant differences between control and diterpenoid-treated samples (Student’s t-test; * = P < 0.05, ** = P < 0.01, *** = P <0.001).

### Barley diterpenoids are modified by *B. sorokiniana*

To further investigate the growth-enhancing effect of 19-OH-HTA on *Bs*, we performed metabolite profiling of axenic *Bs* cultures in the presence of 19-OH-HTA. Untargeted profiling of the whole culture revealed two independent modifications of 19-OH-HTA. The first pathway is through oxidation, resulting in two metabolites with m/z of 331 (compound **37**) and 329 (compound **38**), indicating the addition of a hydroxyl group (m/z 331) or further oxidation to an aldehyde or a ketone (m/z 329) (**Fig. 5A** and **Fig. S13**). We also identified a group of metabolites with m/z of 549 (compounds **39, 40, 41** and **42**). Their mass spectrum indicates that they are conjugates of 19-OH-HTA and a metabolite with the mass of 252 (**Fig. 5B**). Upon negative electrospray ionization, we detect several peaks with an m/z of 251 and the most abundant (compound **43**) was identified as helminthosproric acid based on published data (*26, 27*). Helminthosporic acid is a major sesquiterpenoid metabolite isolated from *Bs* (*27*). Compound **42** was inferred to be an ester of 19-OH-HTA and helminthosporic acid. The other three conjugates are likely to be isomers of **42**, because *Bs* extracts contain several metabolites with the mass of 252, i.e. bipolenin-type isomers of helminthosporic acid (*28*) that most likely can be conjugated to 19-OH-HTA as well. Importantly, the two oxidized diterpenoids and the conjugates are present in the exudates of barley roots infected with *Bs* (**Fig. 5A**), providing evidence that the same modifications of 19-OH-HTA occur during the interaction of barley roots with *Bs*. Notably, during a time course experiment, the appearance of the derivatives of 19-OH-HTA coincided with the enhancement of growth of *Bs* compared to the control (**Fig. S14**).

**Fig. 5.**
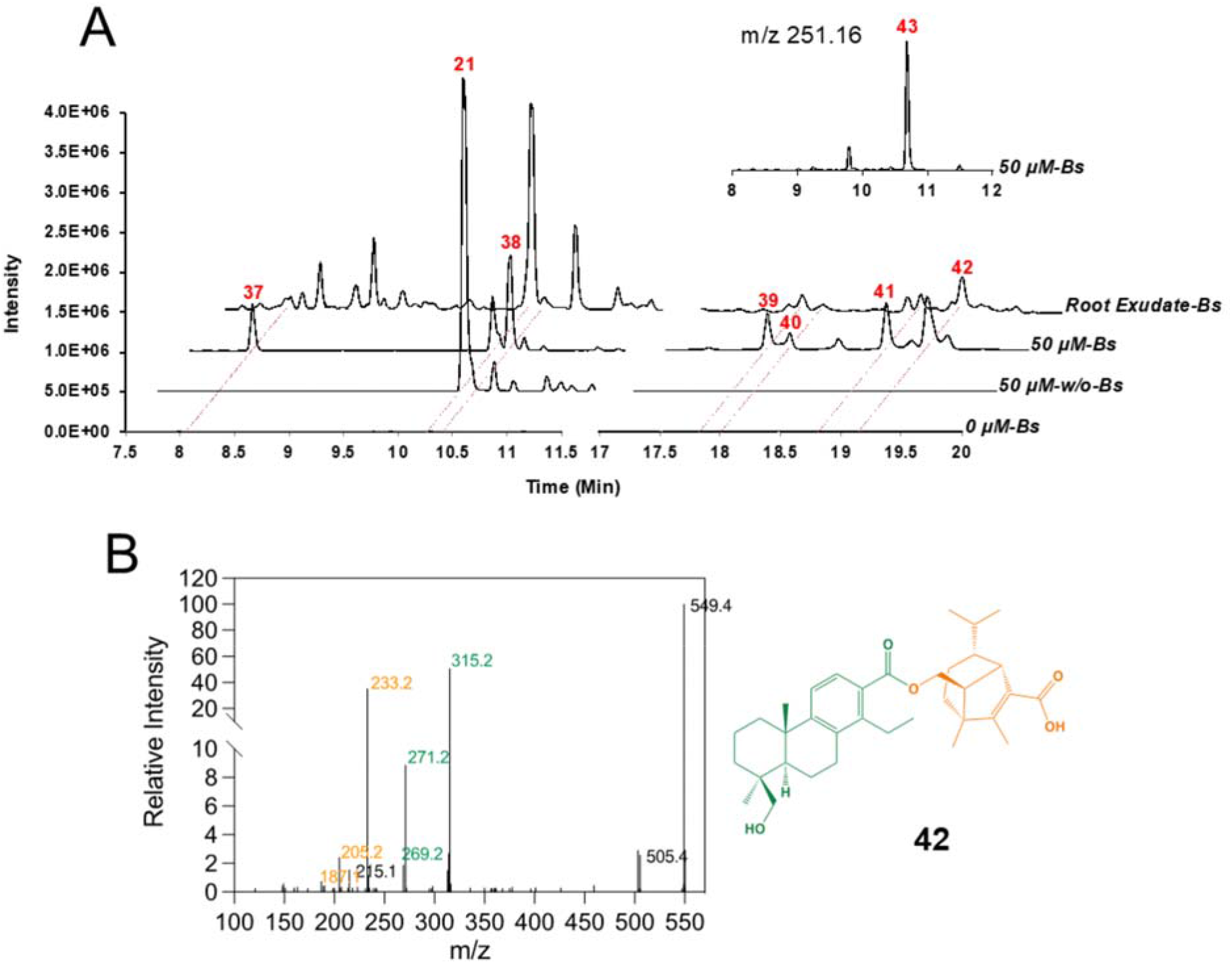
Metabolism of barley diterpenoid 21 by *Bipolaris sorokiniana*. **A)** LC-HRMS (negative mode) chromatogram of extracts from whole *Bs* culture or barly root exudates. *Bs* was grown in the media with or without 19-OH-HTA (**21a**) respectively for three days, and a sample with **21a** at the same concentration but without *Bs* present was used as a control. All experiments were performed in triplicates and were repeated for more than one time. The ions used for the extracted ion chromatograms are m/z 315.2, 329.2, 331.2 and 549.3. The insert shows a part of the chromatogram with ion m/z 251.16 selected, with the peak for compound **43** from *Bs*, which is helminthosporic acid. **B)** MS/MS spectra of **42**. The fragments putatively derived from **21a** are colored in green whilst those from **43** are colored in orange.

### Diterpenoid gene clusters are conserved in Poaceae

Similar gene clusters have been identified in rice and maize, and one would expect to find them in species more closely related to barley, such as wheat. To address this, we performed a synteny analysis of the rice, barley and wheat genomes focusing on the barley diterpenoid cluster (**Fig. S15**). The figure shows strong conservation between the barley hordedane and the rice momilactone cluster, particularly with the presence of CPS, KSL and CYP99 encoding genes. Interestingly, the synteny in wheat is also conserved and distributed over the three genomes (**Fig. S15**). The wheat clusters on the three chromosome 2 all contain a CPS and a KSL, but differ in their content of CYPs, with no CYPs on Chr. 2B, and CYP99s but also CYP71s on chromosome 2A and 2D. This analysis also revealed the presence of several glycosyltransferases, an ABC transporter and a chalcone synthase. We could detect glycosylated 19-OH-HTA in barley (**Fig. S16**), so it is possible that some of these genes are involved in the diterpenoid pathway. Based on these data, it is likely that wheat also produces a set of diterpenoid phytoalexins. To explore this, we inoculated wheat with *Bs* and performed metabolic profiling as for barley diterpenoids. We could detect a number of compounds whose m/z signals and fragmentation patterns are consistent with those of diterpenoids (**Fig. S17)**. Diterpene synthases (a CPS and a KSL) from these clusters have been characterized and most likely provide a backbone for the diterpenoids observed (*29-31*). Beyond wheat, the conservation of this cluster across distant monocots such as rice, barley wheat, but not maize, suggests that many Poaceae are likely to contain similar clusters and thereby contribute to the diversity of the diterpenoid chemical space.

## Discussion

A broadly reported mechanism by which microbial pathogens circumvent the toxicity of phytoalexins is via enzymatic degradation or modification (*32*). Much less frequent is the modification of phytoalexins to modulate plant defense. To our knowledge, the only documented example is that of α-tomatine, a steroidal glycoalkaloid from tomato, which is partially deglycosylated to β_2_-tomatine by tomatinase from the fungal pathogen *Septoria lycopersici* (*33*). In addition to detoxifying α-tomatine, tomatinase and β_2_-tomatine suppress plant defense (*34*). This however was shown in a heterologous host, *N. benthamiana*, and the mechanism by which β_2_-tomatine and tomatinase repress plant defenses is still unknown. Here we show that the barley diterpenoids can be modified by *Bs* and that the growth of *Bs* is enhanced in the presence of 19-OH-HTA. Furthermore, the colonization of barley by *Bs* is reduced on mutants unable to produce the diterpenoids, a phenotype which is reversed by providing 19-OH-HTA to *ksl4* mutants. These data suggest that *Bs* is modifying the barley diterpenoids and using them to increase its growth and capacity to colonize barley. The low concentrations of diterpenoid used and detected in our assays (around or below 50 µM) indicate that they are unlikely to serve as growth substrates. Therefore, a more plausible hypothesis is that these diterpenoids and/or their derivatives act as signalling molecule on the fungus and possibly on the plant host as well. Moreover, because the growth of other fungi is strongly inhibited by 19-OH-HTA, we hypothesize that the modification of diterpenoids is a specific adaptation of *Bs* to the barley metabolic defenses.

Here we show that distinct fungi react differently to the diterpenoid phytoalexin: some are inhibited, another is indifferent and *Bs* is using them to its own advantage. Considering that diterpenoids are only one group of compounds produced in response to pathogens, our work illustrates the complexity of interactions at the metabolite level between pathogens and plant hosts. Deciphering these specific interactions will allow us to better understand plant defense mechanisms besides the classical gene-for-gene resistance.

## Acknowledgments

This work was funded by grants TI 800/7-1 and TI 800/7-2 from the Deutsche Forschungsgemeinschaft to AT. We would like to thank the gardeners of the IPB for caring about the barley plants in the IPB greenhouses.

## Conflict of interest

The authors have no conflict of interest to declare.

## Author contributions

YL performed the metabolite profiling, the cloning, expression in yeast and *N. benthamiana*, purified several diterpenoids and prepared figures and materials. LKM prepared inoculated barley material and performed fungal growth assays. AP and PS performed NMR spectroscopy and analysis. DE performed *Bs* growth assays, fluorescence staining and quantification and complementation experiments 19-OH-HTA. AS-H assisted with metabolite profiling experiments. UB supervised the yeast engineering experiments. IFA produced the CRISPR/Cas knock-outs. AZ provided materials and co-initiated the project. GUB supervised the metabolite profiling, analysed the LC-MS data and performed compound purification. AT conceived and designed the project, acquired funding and wrote the manuscript.

## Notes

### Competing Interest Statement

The authors have declared no competing interest.

## References

1. T. Miedaner, P. Juroszek, Climate change will influence disease resistance breeding in wheat in Northwestern Europe. Theor. Appl. Genet., (2021).

2. U. R. Rosyara, S. Subedi, E. Duveiller, R. C. Sharma, The effect of spot blotch and heat stress on variation of canopy temperature depression, chlorophyll fluorescence and chlorophyll content of hexaploid wheat genotypes. Euphytica 174, 377–390 (2010).

3. D. Sarkar et al., The inconspicuous gatekeeper: endophytic Serendipita vermifera acts as extended plant protection barrier in the rhizosphere. New Phytol 224, 886–901 (2019).

4. I. Ahuja, R. Kissen, A. M. Bones, Phytoalexins in defense against pathogens. Trends in Plant Science 17, 73–90 (2012).

5. N. Ube et al., Evolutionary changes in defensive specialized metabolism in the genus Hordeum. Phytochemistry 141, 1–10 (2017).

6. N. Ube et al., Identification of methoxylchalcones produced in response to CuCl2 treatment and pathogen infection in barley. Phytochemistry 184, 10 (2021).

7. A. Ishihara et al., Induced accumulation of tyramine, serotonin, and related amines in response to Bipolaris sorokiniana infection in barley. Bioscience, Biotechnology, and Biochemistry 81, 1090–1098 (2017).

8. T. Kato et al., Momilactones, growth inhibitors from rice, oryza sativa L. Tetrahedron Letters 14, 3861–3864 (1973).

9. D. Cartwright, P. Langcake, R. J. Pryce, D. P. Leworthy, J. P. Ride, Chemical activation of host defence mechanisms as a basis for crop protection. Nature 267, 511–513 (1977).

10. T. Akatsuka, O. Kodama, H. Kato, Y. Kono, S. Takeuchi, Short Communication 3-Hydroxy-7-oxo-sandaracopimaradiene (Oryzalexin A), a New Phytoalexin Isolated from Rice Blast Leaves. Agricultural and Biological Chemistry 47, 445–447 (1983).

11. Y. Kono, S. Takeuchi, O. Kodama, T. Akatsuka, Absolute Configuration of Oryzalexin A and Structures of Its Related Phytoalexins Isolated from Rice Blast Leaves Infected with <i>Pyricularia oryzae</i>. Agricultural and Biological Chemistry 48, 253–255 (1984).

12. H. Sekido et al., Oryzalexin D (3, 7-Dihydroxy-(+)-sandaracopimaradiene), a New Phytoalexin Isolated from Blast-infected Rice Leaves. Journal of Pesticide Science 11, 369–372 (1986).

13. H. Kato, O. Kodama, T. Akatsuka, Oryzalexin E, A diterpene phytoalexin from UV-irradiated rice leaves. Phytochemistry 33, 79–81 (1993).

14. H. Kato, O. Kodama, T. Akatsuka, Oryzalexin F, a diterpene phytoalexin from UV-irradiated rice leaves. Phytochemistry 36, 299–301 (1994).

15. M. Watanabe et al., Novel C19-Kaurane Type of Diterpene (Oryzalide A), a New Antimicrobial Compound Isolated from Healthy Leaves of a Bacterial Leaf Blight-resistant Cultivar of Rice Plant. Agricultural and Biological Chemistry 54, 1103–1105 (1990).

16. Y. Kono et al., Structures of Oryzalides A and B, and Oryzalic Acid A, a Group of Novel Antimicrobial Diterpenes, Isolated from Healthy Leaves of a Bacterial Leaf Blight-resistant Cultivar of Rice Plant. Agricultural and Biological Chemistry 55, 803–811 (1991).

17. Y. Inoue et al., Identification of a Novel Casbane-Type Diterpene Phytoalexin, <i>ent</i>-10-Oxodepressin, from Rice Leaves. Bioscience, Biotechnology, and Biochemistry 77, 760–765 (2013).

18. E. A. Schmelz et al., Identity, regulation, and activity of inducible diterpenoid phytoalexins in maize. Proceedings of the National Academy of Sciences 108, 5455–5460 (2011).

19. A. Huffaker et al., Novel Acidic Sesquiterpenoids Constitute a Dominant Class of Pathogen-Induced Phytoalexins in Maize Plant Physiology 156, 2082–2097 (2011).

20. K. Shimura et al., Identification of a Biosynthetic Gene Cluster in Rice for Momilactones. Journal of Biological Chemistry 282, 34013–34018 (2007).

21. U. Scheler et al., Elucidation of the biosynthesis of carnosic acid and its reconstitution in yeast. Nat Commun 7, 12942 (2016).

22. K. Bruckner, A. Tissier, High-level diterpene production by transient expression in Nicotiana benthamiana. Plant Methods 9, (2013).

23. Y. Liu et al., A barley gene cluster for the biosynthesis of diterpenoid phytoalexins. bioRxiv, 2021.2005.2021.445084 (2021).

24. J. Liang et al., Deceptive Complexity in Formation of Cleistantha-8,12-diene. Org Lett 24, 2646–2649 (2022).

25. L. Gol, E. B. Haraldsson, M. von Korff, Ppd-H1 integrates drought stress signals to control spike development and flowering time in barley. Journal of Experimental Botany 72, 122–136 (2020).

26. C. S. Phan et al., Bipolenins K-N: New sesquiterpenoids from the fungal plant pathogen Bipolaris sorokiniana. Beilstein J. Org. Chem. 15, 2020–2028 (2019).

27. Y.-Y. Li et al., Bioactive seco-Sativene Sesquiterpenoids from an Artemisia desertorum Endophytic Fungus, Cochliobolus sativus. Journal of Natural Products 83, 1488–1494 (2020).

28. C.-S. Phan et al., Bipolenins K–N: New sesquiterpenoids from the fungal plant pathogen Bipolaris sorokiniana. Beilstein J. Org. Chem. 15, 2020–2028 (2019).

29. Y. Wu et al., Functional characterization of wheat copalyl diphosphate synthases sheds light on the early evolution of labdane-related diterpenoid metabolism in the cereals. Phytochemistry 84, 40–46 (2012).

30. G. Polturak et al., Pathogen-induced biosynthetic pathways encode defense-related molecules in bread wheat. Proc Natl Acad Sci U S A 119, e2123299119 (2022).

31. K. Zhou et al., Functional characterization of wheat ent-kaurene(-like) synthases indicates continuing evolution of labdane-related diterpenoid metabolism in the cereals. Phytochemistry 84, 47–55 (2012).

32. N. M. Westrick, D. L. Smith, M. Kabbage, Disarming the Host: Detoxification of Plant Defense Compounds During Fungal Necrotrophy. Front. Plant Sci. 12, (2021).

33. A. Osbourn, P. Bowyer, P. Lunness, B. Clarke, M. Daniels, Fungal pathogens of oat roots and tomato leaves employ closely related enzymes to detoxify different host plant saponins. Mol Plant Microbe Interact 8, 971–978 (1995).

34. K. Bouarab, R. Melton, J. Peart, D. Baulcombe, A. Osbourn, A saponin-detoxifying enzyme mediates suppression of plant defences. Nature 418, 889–892 (2002).

35. S. Beier et al., Construction of a map-based reference genome sequence for barley, Hordeum vulgare L. Scientific Data 4, 170044 (2017).

36. C. Monat et al., TRITEX: chromosome-scale sequence assembly of Triticeae genomes with open-source tools. Genome Biology 20, 284 (2019).

37. M. Mascher et al., Long-read sequence assembly: a technical evaluation in barley. The Plant Cell 33, 1888–1906 (2021).

