## Supplementary Materials for "Metabolic arms race between a plant and a fungal pathogen"

**The PDF file includes:**

Materials and Methods
Supplementary Text
Figs. S1 to S17
Tables S1 to S13
References

### Materials and Methods

Subhead

Type or paste text here. You can break this section up into subheads as needed (e.g., one on "Materials" and one on "Methods").

Plant growth and fungal inoculations

Barley seeds (*Hordeum vulgare* L. cv Golden Promise) were sterilized in 70% ethanol for 1 min, followed by washing with sterile distilled water and 1.5 h incubation in 12% sodium hypochloride under continuous shaking. After 3 times 30 min washing, the seeds were placed on wet filter paper in darkness and at room temperature for 4 days for germination. Four seedlings were transferred to 1/10 PNM (Plant Nutrition Medium, pH 5.7) (*1*) in sterile glass jars (Height: 147mm, Mouth: 100 mm, WECK) and grown in a day/night cycle of 16/8 h at 22/18 °C, 60% humidity under 108 µmol $m^{-2}$ $s^{-1}$ light intensity.

*Bipolaris sorokiniana* (ND90Pr, *Bs*) was used in this study. *Bs* was propagated on modified CM medium (*2*) with 1.5% agar in the dark at 28 °C for 21 days before inoculation. *Bs* conidia were collected according to the procedures which were described in (*2*). Barley roots were inoculated with 5 ml of *Bs* conidia (5000 spores/ml) per jar for 6 days. Sterile water was used as a mock treatment. Roots washed thoroughly and the corresponding medium were collected and snap-frozen in liquid nitrogen for extraction of metabolites.

Phylogenetic analysis

Phylogenetic analysis was conducted using MEGA X (*3*). Protein sequences (Tab. S2, S3 and S4) were first aligned using ClustalW algorithm (*4*), then phylogenetic trees were generated from the alignment using the maximum likelihood method with a bootstrap of 1,000. For the other parameters, default settings were used.

qRT-PCR

RNA isolation from roots was performed using the Spectrum Plant Total RNA kit (Sigma-Aldrich). The complementary DNA (cDNA) was synthesized using ProtoScript II First Strand cDNA Synthesis Kit (New England Biolabs) following the manufacturer's instructions with primer $d(T)_{23}$ VN. Quantitative real-time PCR was performed in triplicates using 10-20 ng cDNA as template and gene specific primer pairs shown in **Tab. S11** in CFX Connect Real-Time PCR System (Bio-Rad). The PCR conditions were 95 °C for 15 min; 40 cycles of 95 °C for 15 s, 60 °C for 30 s; 95 °C for 10 s. The melting curve was measured from 65 °C to 95 °C with a step of 0.1 °C per second. Relative expression of targeted genes was calculated using delta Ct method (*5*) and the barley ubiquitin gene as reference (*6*).

Heterologous expression of diterpenoids in yeast

Plasmids containing *GGPPS* and *ATR1* were kindly provided by colleagues in the group and have been described previously (*7*). Codon-optimized DNA sequences of *HvCPS2, HvKSL4*, *HvCYP89E31*, *HvCYP99A66*, *HvCYP99A67* and *HvCYP99A68*, were synthesized by Thermo Fisher Scientific Inc. for yeast expression. Each gene was further cloned into Golden Gate compatible yeast expression level 1 vector, together with a synthetic galactose-inducible promoter and a terminator. Different gene combinations were finally assembled into one yeast expression level M vector by a 50-cycle restriction–ligation reaction with BpiI and T4-Ligase.

Constructs were then transformed into *Saccharomyces cerevisiae* strain INVSc1 (Thermo Fisher Scientific Inc.) and plated out onto uracil-free (Ura-) selection medium (1 g $l^{-1}$ Yeast Synthetic Drop-out Medium Supplements without uracil (Sigma-Aldrich), 6.7 g $l^{-1}$ Yeast Nitrogen Base with Amino Acids (Sigma-Aldrich) and 20 g $l^{-1}$ Micro Agar (Duchefa Biochemie)). Three positive colonies were picked and inoculated into 5 ml yeast extract-peptone-dextrose (YPD) medium (20 g $l^{-1}$ tryptone and 10 g $l^{-1}$ yeast extract) containing 2% of glucose and grown for 24 h with shaking at 30 °C. To induce protein expression, the cell pellet was resuspended in fresh YPD medium containing 2% galactose. After another 24 h of growth, the whole culture was extracted with 2 ml *n*-hexane.

Transient expression in *Nicotiana Benthamiana*

Transit peptides of protein HvCPS2 and HvKSL4 were predicted by two online tools, ChloroP 1.1 (http://www.cbs.dtu.dk/services/ChloroP/) and LOCALIZER (http://localizer.csiro.au/). Truncated sequences without the predicted transit peptides of these two genes were generated by PCR reactions using designed primers (**Tab. S11**) and subsequently sequenced. The cDNAs of truncated *HMG reductase*, *GGPPS* in plasmids have been described previously (*7, 8*). The *trHMG reductase*, *GGPPS*, *trHvCPS2*, *trHvKSL4*, *HvCYP89E31*, *HvCYP99A66*, *HvCYP99A67* and *HvCYP99A68* were cloned into T-DNA vectors (binary vector pL1F-1) driven by the 35S promoter and flanked by the Ocs terminator (*9*). The resulting T-DNA plasmids were transformed into *Agrobacterium tumefaciens* strain GV3101::pMP90 and plated out onto LB agar plates with appropriate antibiotics. Bacteria were harvested and resuspended in infiltration medium (10 mM $MgCl_2$, 10 mM MES, 20 μM acetosyringone, pH=5.6) after 48 h inoculation at 28 °C. To co-infiltrate several genes, each bacteria suspension was diluted to a final $OD_{600}$ of 0.4, then all strains were mixed equally to an appropriate volume for infiltration. The suspension was infiltrated into the abaxial side of several leaves in three individual 4-week-old *N. benthamiana* plants using a syringe without needle. After treatment, the plants were cultivated in a climate controlled phytochamber for 4 days. Three leaf discs (9 mm diameter) per infiltrated spot were harvested and extracted by 2 ml *n*-hexane, followed by drying down under nitrogen flow and GC/LC-MS analysis.

Microsome isolation and *in vitro* enzyme assay

A protocol from the literature with slight modification was used for microsome isolation (*7, 10*). The construct carrying *HvCYP89E31* and *ATR1* were transformed into yeast strain INVSc1. A single positive colony was picked to inoculate 5 ml of Ura-medium with 2% glucose and grown for 24 h at 30 °C with shaking. The culture was then used to inoculate 100 ml of Ura-medium with 2% glucose in a 500 ml flask at 30 °C for 24 h. The cells were then collected by centrifugation, resuspended in 100 ml fresh YPD medium with 2% galactose to induce protein expression and inoculated under shaking for another 24 h at 30 °C. All the following steps were carried out at 4 °C. The cells were harvested by centrifugation and resuspended in 30 ml of pre-chilled TEK buffer (50 mM Tris-HCl pH 7.5, 1 mM EDTA, 100 mM KCL), centrifuged again and resuspended in 2 ml TES buffer (50 mM Tris-HCl pH 7.5, 600 mM sorbitol, 10 g $l^{-1}$ BSA, 1.5 mM β-mercaptoethanol) and transferred to a 50 ml tube. Acid-washed autoclaved 450–600 μm diameter glass beads were added into the tube until the surface of the cell suspension are reached. The suspension was shaken vigorously by hand for 1 min and returned to ice for 1 min. This step was repeated four times. The glass beads were washed by 5 ml TES buffer three times, and the supernatant was collected and combined to a new tube, followed by centrifugation at

7,500 g for 10 min. The supernatant was transferred to ultracentrifugation tubes and centrifuged for 2 h at 100 000 g. The pellet was gently washed successively with 5 ml TES and 2.5 ml TEG buffer (50 mM Tris-HCl pH 7.5, 1 mM EDTA and 30% glycerol) after the supernatant was removed, then transferred to a Potter homogenizer with a spatula. 2 ml TEG buffer was added to the homogenizer and the pellet was carefully homogenized. 100 µl aliquots were transferred to 1.5 ml microtubes and stored at −80 °C until used.

*In vitro* CYP enzyme assays were performed in a 600 µl reaction volume, containing 40 µl of microsome preparation, 100 µM substrate, 1 mM NADPH, 50 mM sodium phosphate pH 7.4. The solution was incubated at 30 °C for 2 h with gentle shaking. Products were extracted with 1 ml *n*-hexane under strong agitation (vortex). After centrifugation, the organic phase was collected, then dried under a $N_2$ stream and resuspended in 100 µl *n*-hexane for GC-MS analysis.

Purification of diterpenoids from yeast culture by silica gel column chromatography or SPE

For the purification of hordediene (**1**), 1 L of yeast culture was grown and extracted with 1 L *n*-hexane. The raw extracts were dried in a rotary evaporator and resuspended in 4 ml *n*-hexane, then loaded into two properly conditioned SiOH SPE cartridges (500 mg, MACHEREY-NAGEL). The cartridges were then washed with 2 ml n-hexane. The breakthrough and washing fraction were collected and combined. After drying down under nitrogen stream, an aliquot was measured by GC-MS to check the purity of the product and the rest, with an amount of around 2 mg was used for NMR structure elucidation.

Diterpenoids **5, 6** and **8** were first extracted from three liters of yeast culture. After concentration, the raw extracts were dissolved in 4 ml *n*-hexane and loaded into a pre-conditioned self-packed silica gel column (5 g, 15 mm x 100 mm). The column was then eluted by *n*-hexane and successively by 99:1, 98:2, 97:3, 96:4, 95:5 *n*-hexane: ethyl acetate solutions. The volume of every elution solution was 10 ml, and each elution was sequentially collected into five 2 ml microtubes. An aliquot of each fraction was measured by GC-MS and the fractions with the same product were combined and then used for NMR structure elucidation. The yields of diterpenoids **5** and **8** were around 0.5 mg, and **6** was about 1 mg.

Diterpenoids **19** and **21** were first extracted from three to five liters of yeast culture by the same volume of extraction solution (95:5 *n*-hexane: ethyl acetate). After concentration, the raw extracts were dissolved in 4 ml *n*-hexane and loaded into two properly conditioned SiOH SPE cartridges (500 mg, MACHEREY-NAGEL). The column was then washed by 2 ml *n*-hexane and successively by 2 ml 95:5, 90:10, 85:15 *n*-hexane: ethyl acetate solutions, followed by elution using 80:20 *n*-hexane: ethyl acetate for **19** or 75:25 *n*-hexane: ethyl acetate for **21**. The volume of elution solution was 10 ml, but the elution was separately collected into five 2 ml microtubes. An aliquot of each fraction was measured by LC-MS and the fractions with the same product were combined and then used for NMR structure elucidation. The yields of diterpenoids **19** and **21** were around 1 mg and 0.5 mg respectively.

Purification of diterpenoid 35 from barley root exudates

Purification of semi-polar diterpenes was achieved by micro-SPE-UPLC-MS on a prototype device (Axel Semrau GmbH, Sprockhövel, Germany). In brief, 400 µL of methanolic barley or yeast extracts (dissolved in 80% methanol acidified to pH2.4) were injected to micro solid phase extraction (SPE) cartridges (CHROspe Polymer DVB 10 x 2 mm, 25-35 µM, SparkHolland B.V., Emmen, The Netherlands) under concomitant dilution by a second pump with an excess of water. Next, adsorbed diterpenoids were transferred from the SPE cartridge to an UPLC column

(Nucleoshell RP18, 2 mm x 100 mm x 2.7 µm) with 400 µL of 80% acetonitrile used for desorption. During the transfer, the eluate was diluted with water again to give a final concentration of 8% acetonitrile on column throughout the transfer process. Barley and yeast-derived diterpenoids were chromatographically separated using the gradient described above but fractions were collected from the UPLC effluent. Micro-SPE was again used for fraction collection, and the LC effluent was again diluted with excess water to retain the desired products. Finally, the SPE cartridges were desorbed with pure methanol and the eluates were dried in a nitrogen stream.

Nuclear magnetic resonance (NMR) conditions

$^{1}H$, $^{13}C$, 2D ($^{1}H$,$^{1}H$ gDQCOSY; $^{1}H$,$^{1}H$ zTOCSY;$^{1}H$,$^{1}H$ ROESYAD; $^{1}H$,$^{13}C$ gHSQCAD; $^{1}H$,$^{13}C$ gHMBCAD), selective ($^{1}H$,$^{1}H$ zTOCSY1D; $^{1}H$,$^{1}H$ ROESY1D), and band selective ($^{1}H$,$^{13}C$ bsHMBC) NMR spectra were measured with an Agilent VNMRS 600 instrument at 599.83 MHz ($^{1}H$) and 150.84 MHz ($^{13}C$) using standard CHEMPACK 8.1 pulse sequences implemented in the VNMRJ 4.2A spectrometer software. TOCSY mixing time: 80 ms; ROESY mixing time: 300 ms; HSQC optimized for $^{1}J_{CH}$ = 146 Hz; HMBC optimized for $^{n}J_{CH}$ = 8 Hz. All spectra were obtained with $C_6D_6$ + 0.03% TMS as solvent at +25°C. Chemical shifts were referenced to internal TMS (δ $^{1}H$ = 0 ppm) and internal $C_6D_6$ (δ $^{13}C$ = 128.0 ppm).

Metabolites extraction from barley roots and PNM medium

100 mg (fresh weight) of frozen and cryo-ground root matter was extracted using 900 µl dichloromethane/ethanol (2:1, v/v) and 100 µl hydrochloric acid solution (pH 1.4). Extraction and duplicate removal of hydrophilic metabolites was achieved by 1 min FastPrep bead milling (FastPrep24, MP Biomedicals) followed by phase separation during centrifugation. For extraction 1.6 ml wall-reinforced cryo-tubes (Biozyme) each containing steel and glass beads were used. The upper aqueous phase was discarded and replaced for a second round of bead mill extraction/centrifugation. Thereafter, the aqueous phase was removed, and the lower organic phase was collected. Subsequently 600 ul tetrahydrofuran (THF) was used for exhaustive extraction (FastPrep). After centrifugation the organic THF extract was combined with the first extract and dried under a stream of $N_2$.

Root exudates were extracted from 60 ml of Gelrite media. For this, the gel was distributed into two 50 ml Falcon tubes. To each tube, 4 ml ethyl acetate were added. The tubes were thoroughly shaken by hand and centrifuged. The upper phase (organic extract) was collected before fresh ethyl acetate was added for another two consecutive extractions. The combined extracts of three extraction rounds were combined and dried in a stream of $N_2$.

GC-MS

Dried extracts were suspended in 200 µl n-hexane. The analysis of yeast and plant extracts was carried out using a Trace GC Ultra gas chromatograph (Thermo Scientific) coupled to an ATAS Optic 3 injector and an ISQ single quadrupole mass spectrometer (Thermo Scientific) with electron impact ionization. Chromatographic separation was performed on a ZB-5MS capillary column (30 m × 0.25 mm x 0.25 mm, Phenomenex) using splitless injection and an injection volume of 1 µl. The injection temperature rose from 60 °C to 250 °C with 7 °C $s^{-1}$ and the flow rate of helium was 2 ml $min^{-1}$. The GC oven temperature ramp was as follows: 50 °C for 1 min, 50 to 300 °C with 7 °C $min^{-1}$, 300 to 330 °C with 20 °C $min^{-1}$ and 330 °C for 5 min.

Mass spectrometry was performed at 70 eV, in a full scan mode with m/z from 50 to 450. Data analysis was done with the device specific software Xcalibur (Thermo Scientific).

Some samples were analyzed on a ZB-5HT capillary column (30 m × 0.25 mm x 0.25 mm, Phenomenex) using splitless injection and an injection volume of 1 µl. The injection temperature rose from 130 °C to 280 °C with 5 °C $s^{-1}$ and the flow rate of helium was 2 ml $min^{-1}$. The GC oven temperature ramp was as follows: 130 °C for 2 min, 130 to 250 °C with 8 °C $min^{-1}$, 250 to 310 °C with 10 °C $min^{-1}$ and 310 °C for 5 min. MS spectra were acquired using the same parameters which are described above.

RP-UPLC-ESI-MS/MS

For UPLC-MS/MS analysis dried extracts were suspended in 180 µl 80% methanol/ 20% water. Separation of medium polar metabolites was performed on a Nucleoshell RP18 (2.1 × 150 mm, particle size 2.1 µm, Macherey & Nagel, GmbH, Düren, Germany) using a Waters ACQUITY UPLC System, equipped with a Binary Solvent Manager and Sample Manager (20 µl sample loop, partial loop injection mode, 5 µl injection volume, Waters GmbH Eschborn, Germany). Eluents A and B were aqueous 0.3 mmol/l NH4HCOO (adjusted to pH 3.5 with formic acid) and acetonitrile, respectively. Elution was performed isocratically for 2 min at 5% eluent B, from 2 to 19 min with a linear gradient to 95% B, from 19-21 min isocratically at 95% B, and from 21.01 min to 24 min at 5% B. The flow rate was set to 400 µl $min^{-1}$ and the column temperature was maintained at 40 °C. Metabolites were detected by positive and negative electrospray ionization and mass spectrometry.

Mass spectrometric analysis of small molecules was performed by MS-TOF-SWATH-MS/MS (TripleToF 5600, AB Sciex GmbH, Darmstadt, Germany) operating in negative or positive ion mode and controlled by Analyst 1.7.1 software (AB Sciex GmbH, Darmstadt, Germany). The source operation parameters were as follows: ion spray voltage, −4500 V / +5500 V; nebulizing gas, 60 psi; source temperature 600 °C; drying gas, 70 psi; curtain gas, 35 psi. TripleToF instrument tuning and internal mass calibration were performed every 5 samples with the calibrant delivery system applying APCI negative or positive tuning solution (AB Sciex GmbH, Darmstadt, Germany), respectively.

TripleToF data acquisition was performed in $MS^1$-ToF mode and $MS^2$-SWATH mode. For $MS^1$ measurements, ToF masses were scanned between 65 and 1250 Dalton with an accumulation time of 50 ms and a collision energy of 10 V (−10 V). $MS^2$-SWATH experiments were divided into 26 Dalton segments of 20 ms accumulation time. Together the SWATH experiments covered the entire mass range from 65 to 1250 Dalton in 48 separate scan experiments, which allowed a cycle time of 1.1 s. Throughout all MS/MS scans a declustering potential of 35 V (or −35 V) was applied. Collision energies for all SWATH-MS/MS were set to 35 V (−35 V) and a collision energy spread of ±25 V, maximum sensitivity scanning, and elsewise default settings.

For some samples, mass spectrometric analysis of small molecules was performed by MS-TOF-IDA-MS/MS (ZenoTOF 7600, AB Sciex GmbH, Germany) operating in negative or positive ion mode and controlled by SCIEX OS software 2.1.6 (AB Sciex GmbH, Darmstadt, Germany). The source operation parameters were as follows: ion spray voltage, −4500 V / +5500 V; ion source gas 1, 60 psi; source temperature 600 °C; ion source gas 2, 70 psi; curtain gas, 35 psi; CAD gas 7. ZenoTOF instrument tuning and internal mass calibration were performed every 5 samples with the calibrant delivery system applying X500 ESI negative or positive calibration solution (AB Sciex GmbH, Germany), respectively.

ZenoTOF data acquisition was performed in $MS^1$-ToF mode and $MS^2$-IDA mode. For $MS^1$ measurements, ToF masses were scanned between 65 and 1500 Dalton with an accumulation time of 100 ms and a collision energy of 10 V (−10 V). $MS^2$-IDA experiments were performed using the following parameters: ToF mass range 65 to 1500; declustering potential of 80 V (or −80 V) with a spread of 50; maximum candidate ions of 40 with an intensity threshold of 1000 cps; collision energy of 35 V (−35 V) with a spread of 25 V; Zeno threshold 20000 cps; accumulation time of 20 ms. Total scan time of one cycle for both $MS^1$ and $MS^2$ was 1.166 s.

Mutation of *HvCPS2* and *HvKSL4* by CRISPR/Cas9 gene editing

The CRISPOR web tool (Concordet and Haeussler, 2018) Reference: https://academic.oup.com/nar/article/46/W1/W242/4995687 was used to design two single guide RNAs for each gene, as follows (PAM sequence underlined):
HvCPS2_sgRNA1: 5'-GAAGAGTAGGGTCGTTGGTATGG-3'
HvCPS2_gRNA2: 5'-AAGTTAATCTCGAAGCCACATGG-3'
HvKSL4_gRNA1: 5'-GGCTGGTGAGTCAAATTCACCGG-3'
HvKSL4_gRNA2: 5'-TTAATGTGATAGTTCGCCATCGG-3'
Golden Gate cloning was used to load each pair of guide sequences from complementary oligonucleotides into shuttle vectors pMGE625 and pMGE627, which were then assembled into binary vector pMGE599 to create pMP222 (HvCPS2) and pMP223 (HvKSL4). Vectors and cloning protocols have been previously described (Kumar et al., 2018) Reference: https://pubmed.ncbi.nlm.nih.gov/29577542/ and were kindly provided by Johannes Stuttmann. Stable transformation of pMP222 and pMP223 in Golden Promise Fast was performed as described (Amanda et al., 2022). Reference: https://pubmed.ncbi.nlm.nih.gov/35316655/

Root staining and fluorescence quantification

After harvest, the roots were gently washed in distilled water, cut in 1cm pieces, and placed in 2ml test tube filled with 1.5ml of 50% Ethanol (EtOH) and left overnight at 4°C. After removing the EtOH, the root segments were incubated in 20% potassium hydroxide (KOH) for 10’ at 90°C. The KOH was then removed, and the samples were washed three times with distilled water. Afterwards, 0.1M hydrochloric acid (HCl) was added and then left at room temperature over 1.5h. Another round of three wash with distilled water was then carried out. The roots were then rinsed once with phosphate buffered saline (BPS) and then incubated in PBS-WGA staining solution (WGA-AlexaFluor488 final concentration of 0.2ug/ml) overnight, wrapped in aluminum foil, at 4°C.

The fluorescence of each segment was then measured in a TECAN device (Tecan Group Ltd., Switzerland) with the following parameters: excitation wavelength 485nm; emission wavelength 535nm; gain 40% calculated on a wild-type sample; mirror Dichroic 510; 30 flashes; z-position defined on a wild-type sample; 15*15 reads per well with a 2500um border. Each run, four roots section per sample were analyzed. Eight run per sample were carried, for a total of 32 segments per sample analyzed. Two sample per genotype were analyzed. After measuring, the average of each run was computed and used for statistical analysis.

Bioassay with 21 on five fungi

*Bipolaris sorokiniana* (ND90Pr; *Bs*) was grown on modified CM-Bs medium (*2*). *Serendipita vermifera* (MAFF305830; *Sv*) was grown on MYP medium (*11*). *Serendipita indica*

(DSM11827; *Si*) was grown on CM medium (*12*). *Verticilium dahlie* (JR2; *Vd*) was grown on PDA medium. *Fusarium culmorum* (KF350; *Fc*) was grown on YPSs medium (*13*). *Verticilium dahlie* was grown at room temperature in darkness for 7 days. All other fungi were grown at 28°C for 3 weeks. Spores were harvested as described previously (*2*) and diluted to a final concentration of 250,000 spores/ml for *Bs*, 700,000 spores/ml for *Si*, 552,000 spores/ml for *Fc* and 542,500 spores/ml for *Vd* respectively in 1/2 TSB (Sigma Aldrich; 15 g/l). *Sv* small mycelium fragments were harvested by adding water to the plate, removing the loose hyphae with a scalpel, and filtering the solution through miracloth. The solution was centrifuged, the supernatant was discarded, and the pellet was dissolved in 1/2 TSB medium. 400 µl of the fungal solutions were applied to each microscope chamber slide (VWR; Kat734-2050) and treated with either 1% DMSO as a control or 1% DMSO + 50 µM of the barley diterpenoid **21a**. Pictures were taken at different timepoints depending on the speed of spore germination (3 hours post inoculation (hpi) for *Bs*; 9 hpi for *Vd*; 9 hpi for *Fc*; 12 hpi for *Si*; and 12 hpi for *Sv*). Spore germination rate and tube elongation/hyphal length were quantified using imageJ (*14*).

Bioassay with 21a (19-OH-HTA) on *Bs* (Growth curve)

To measure the growth curve of *Bs* in different concentrations of **21a**, the following assay was performed. Microplate wells were filled with 200 µl of working solution with different concentration of **21**. Each concentration was tested with ten replicates. After pipetting, the plate was inoculated at 28 °C in either an incubator or TECAN device (Tecan Group Ltd., Switzerland). The value of $OD_{405}$ was measured by TECAN every one hour for 120 h.

**Components of working solution**

| **21 in Concentration** (Final) | **21a stock** (20 mM in DMSO) | **DMSO** | **CM medium** | ***Bs* Spores** ($3x10^5$/ml) |
|---|---|---|---|---|
| **50 µM** | 2.5 µl | 17.5 µl | 880 µl | 100 µl |
| **25 µM** | 1.25 µl | 18.75 µl | 880 µl | 100 µl |
| **0 µM** | 0 | 20 µl | 880 µl | 100 µl |
| **Blank** | 0 | 20 µl | 980 µl | 0 |

Note: final concentrations of **21a**: 0, 25, 50 µM in CM medium with 2% DMSO.

To measure degraded metabolites, the following extraction protocol was used. The *Bs* culture was transferred from the wells to 2 ml Eppendorf tubes. Each sample was extracted by two times of 500 µl ethyl acetate, and the extractions were combined in the end. During extraction, the ultrasonic bath and vertex device were used to improve extraction efficiency. The extracts were then analyzed by LC-MS/MS.

Complementation assay

*Growing of yeast and extraction*

Two different vectors were used, namely: pAGT1009 and pAGT9593. With the first being an empty vector a the second withall the genes required to produce the 19-OH-HTA. These were then transformed into *S. cerevisiae* strain INVSc1 and the positive colonies grown in uracil-free medium (1 g $l^{-1}$ Yeast Synthetic Drop-out Medium Supplements without uracil (Sigma-Aldrich), 6.7 g $l^{-1}$ Yeast Nitrogen Base with Amino Acids (Sigma-Aldrich)) containing 2% of glucose over two days, shaking at 30°C. The inoculum was then transferred to fresh media containing 2% galactose to induce protein expression and grown over three days in the same conditions. A total of 2.4 liters of media were used for each strain. The whole culture was then let to sediment overnight at 4°C and both the supernatant and the pellet were extracted three times with 0.4 volumes of 30/70 Ethyl-acetate/n-Hexane (EtAc/Hex) HPLC grade. The organic fractions were then combined and dried using a rotary evaporator.

*Purification*

The dried extract, of both yeast with pAGT1009 (sample y1009) and yeast with pAGT9593 (sample y9593), was resuspended in 1ml of 10/90 EtAc/Hex, centrifuged at maximum speed for 3 minutes, 5°C, and loaded in a pre-conditioned normal-phase (NP) SiOH solid-phase-extraction (SPE) cartridges (500 mg, MACHEREY-NAGEL). The residual pellet was then washed another two times in the same way. After washing the cartridges with 2ml of 10/90 EtAc/Hex and 2ml of 20/80 EtAc/Hex, the sample was eluted from it with 4ml 30/70 EtAc/Hex and dried. This whole procedure was carried twice to reduce the number of contaminants present. The final amount of **21a** in the sample y9593 (around 100 µg) was estimated via LC-MS.

*Preparation of the glass jars*

60ml of barley media were poured into a sterile glass jar (Height: 147mm, Mouth: 100 mm, WECK) where a small glass vial was sitting in the center. This was then removed once the agar solidified, leaving behind a hole of around 5ml volume in which fresh agar, enriched with the different extracts, was later poured. Before adding the extracts to the agar, they were resuspended in pure DMSO to have a final concentration of 2% in the 5ml agar plug.

*Seeds sterilization and plant growth*

Barley (*H. vulgare* L.) seeds cv. Golden promise fast and from the mutant *ksl4-1* were sterilized with 6% hypochlorite under continuous shaking for 1h. Then washed three times with sterile distilled water and four times under continuous shaking for 30’ each round. The seeds were then placed on wet filter paper and let germinate in darkness at room temperature over three days. The seedlings were then moved to the jars (one per jar), taking care of positioning the roots over the yeast extract plug and let grow for two more days. Then the roots were inoculated with 5ml of *Bs* conidia (5000 spores/ml), collected as described before (cf. Plant growth and fungal inoculations) and grown over six days in a day/night cycle of 16/8 h at 22/18 °C, 60% humidity under 108 µmol m-2 s-1 light intensity. Sterile water was used as a mock treatment. Roots washed thoroughly and the corresponding medium were collected and snap-frozen in liquid nitrogen for extraction of metabolites and RNA extraction.

*RNA extraction, cDNA synthesis and Bs quantification via qPCR.*

See section on qRT-PCR.

Synteny analysis

The synteny analysis was performed by MCScanX (*15*), employing the parameters ‘-s 5 -w 0 -m 15’. The orthologs were identified by blastp program, using the parameters ‘E-value 1e-10; num of best hits 10’. The plot for micro-synteny was generated by MCScan (Python version) (*16*). The version of genomes used for the analysis is IRGSP-1.0 of rice, MorexV2 of barley, and IWGSC RefSeq v1.0 of wheat.

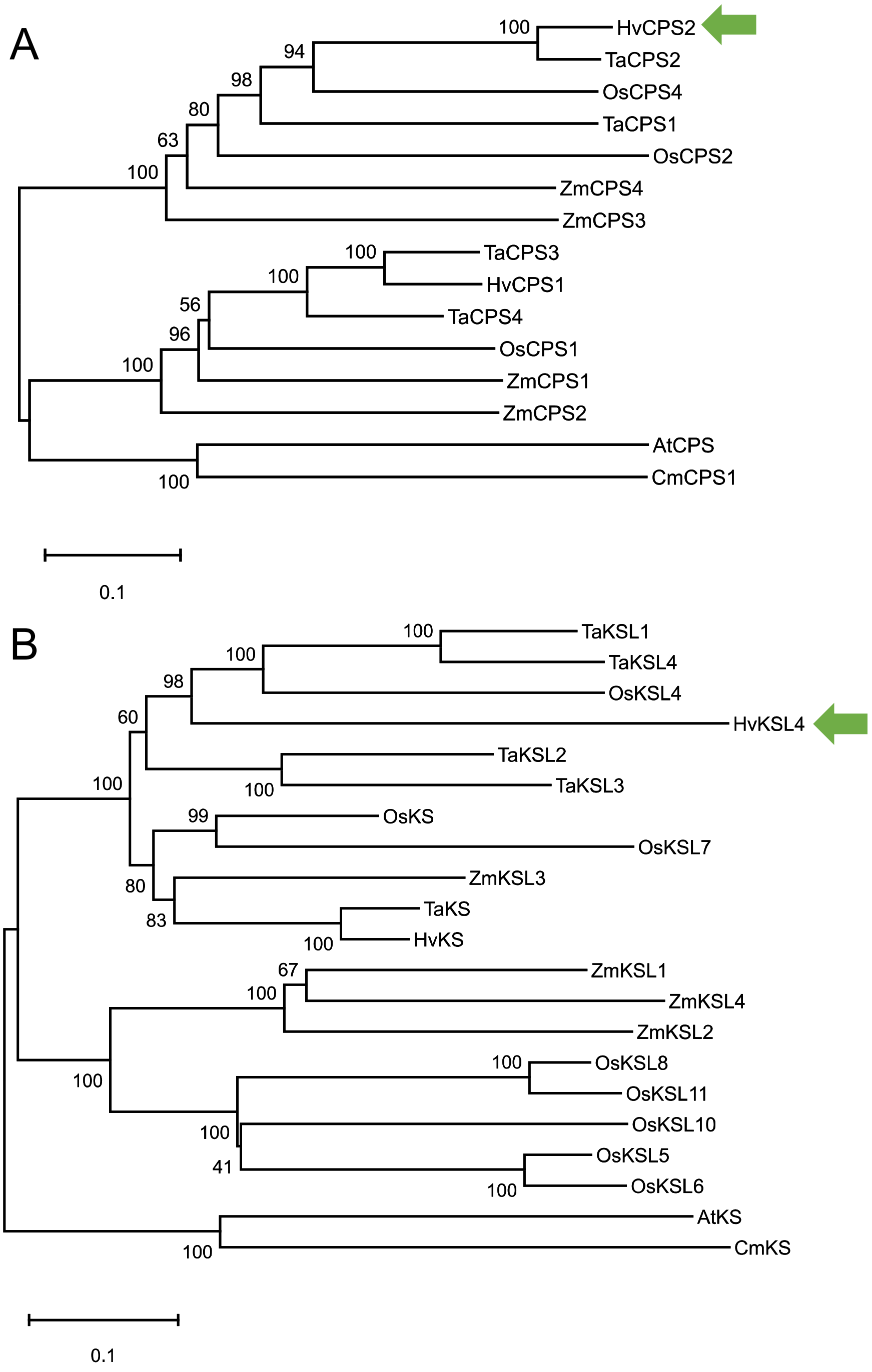

**Fig. S1. Phylogenetic analysis of HvCPS2 and HvKSL4.**

The protein sequences indicated were aligned by ClustalW (*4*) and further processed with the MEGA X software (*3*) using the maximum likelihood method with a bootstrap of 500. The tree with the highest log likelihood is shown. For other parameters, default settings were used. The HvCPS2 and HvKSL4 sequences are indicated by green arrows. The list of sequences used is provided in **Tab. S2** and **S3**.

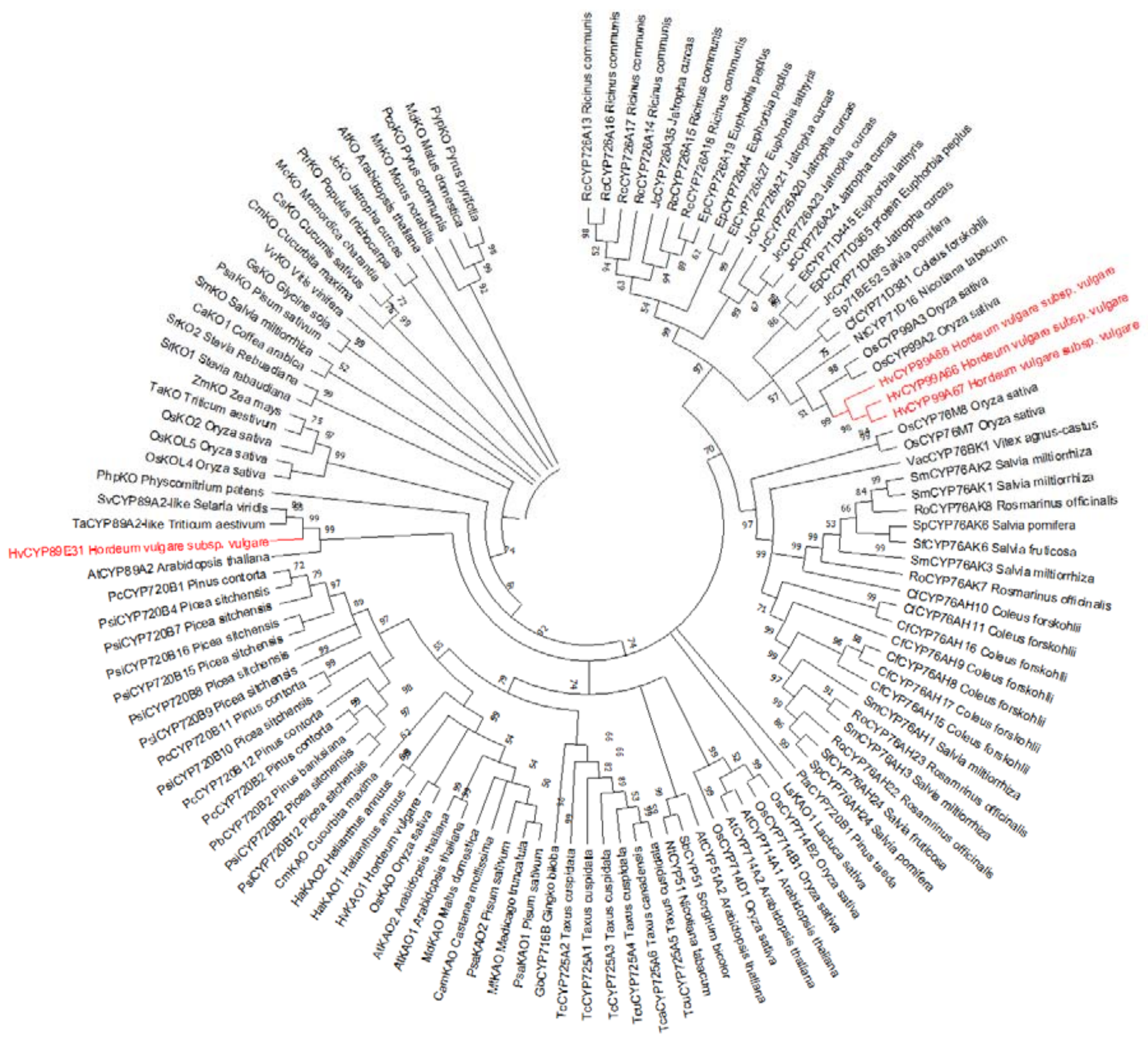

**Fig. S2. Phylogenetic analysis of CYPs from barley chromosome 2 diterpenoid cluster.**
The evolutionary history was inferred using the Maximum Likelihood method and Poisson correction model. The bootstrap consensus tree inferred from 1000 replicates is taken to represent the evolutionary history of the taxa analyzed. Branches corresponding to partitions reproduced in less than 50% of bootstrap replicates are collapsed. Evolutionary analyses were conducted in MEGA X (*3*). The list of sequences used is provided in **Tab. S4**. The barley CYPs are highlighted in red.

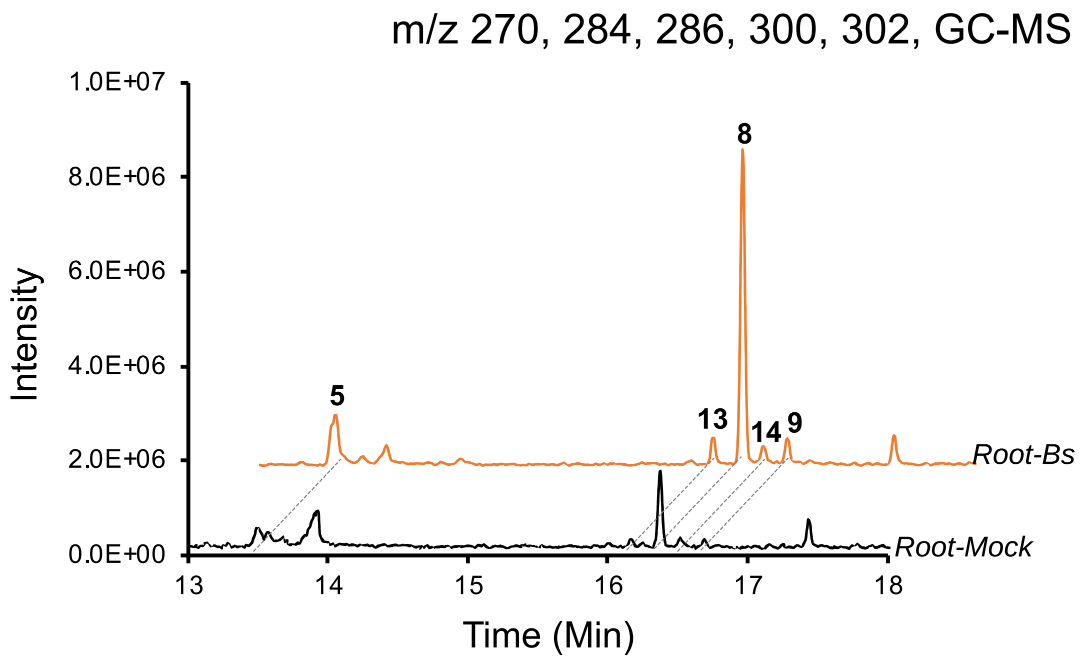

**Fig. S3. GC-MS chromatograms of extracts from barley roots inoculated with *Bs* or mock treatment.**

Extracted ion chromatograms (m/z 270, 284, 286, 300, 302) of extracts from barley roots infected with *Bs* (Root-Bs) or mock-inoculated (Root-Mock).

Type or paste caption here. Create a page break and paste in the figure above the caption.

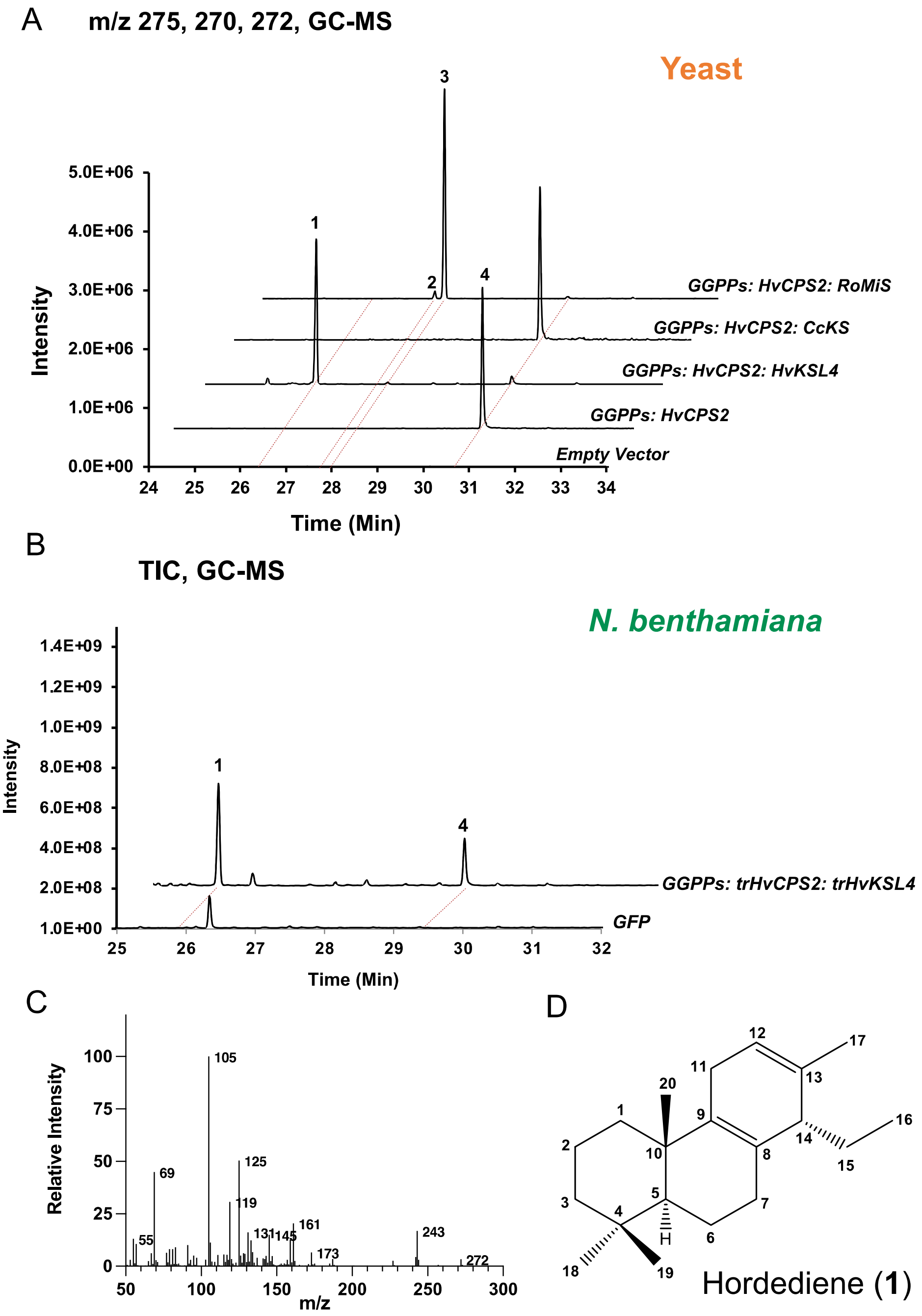

**Fig. S4. Characterization of HvCPS2 and HvKSL4 in yeast and *N. benthamiana*.**

**A)** GC-MS chromatograms of extracts of yeast strains expressing the gene combinations indicated on the right. CcKS: ent-kaurene synthase from coffee (*Coffea canephora*); RoMiS, miltiradiene synthase from rosemary (*Rosmarinus officinalis*). **1**: hordediene; **2**: miltiradiene; **3**: abietatriene; **4**: (+)-copalol. Selected ions, m/z 270, 272, and 275. **B)** GC-MS analysis of transient expression in *N. benthamiana*. The genes that were expressed are indicated on the right side of the chromatogram. Total ion chromatograms (TIC) are shown. **C)** EI mass spectrum of hordediene. **D)** Structure of hordediene determined by NMR.

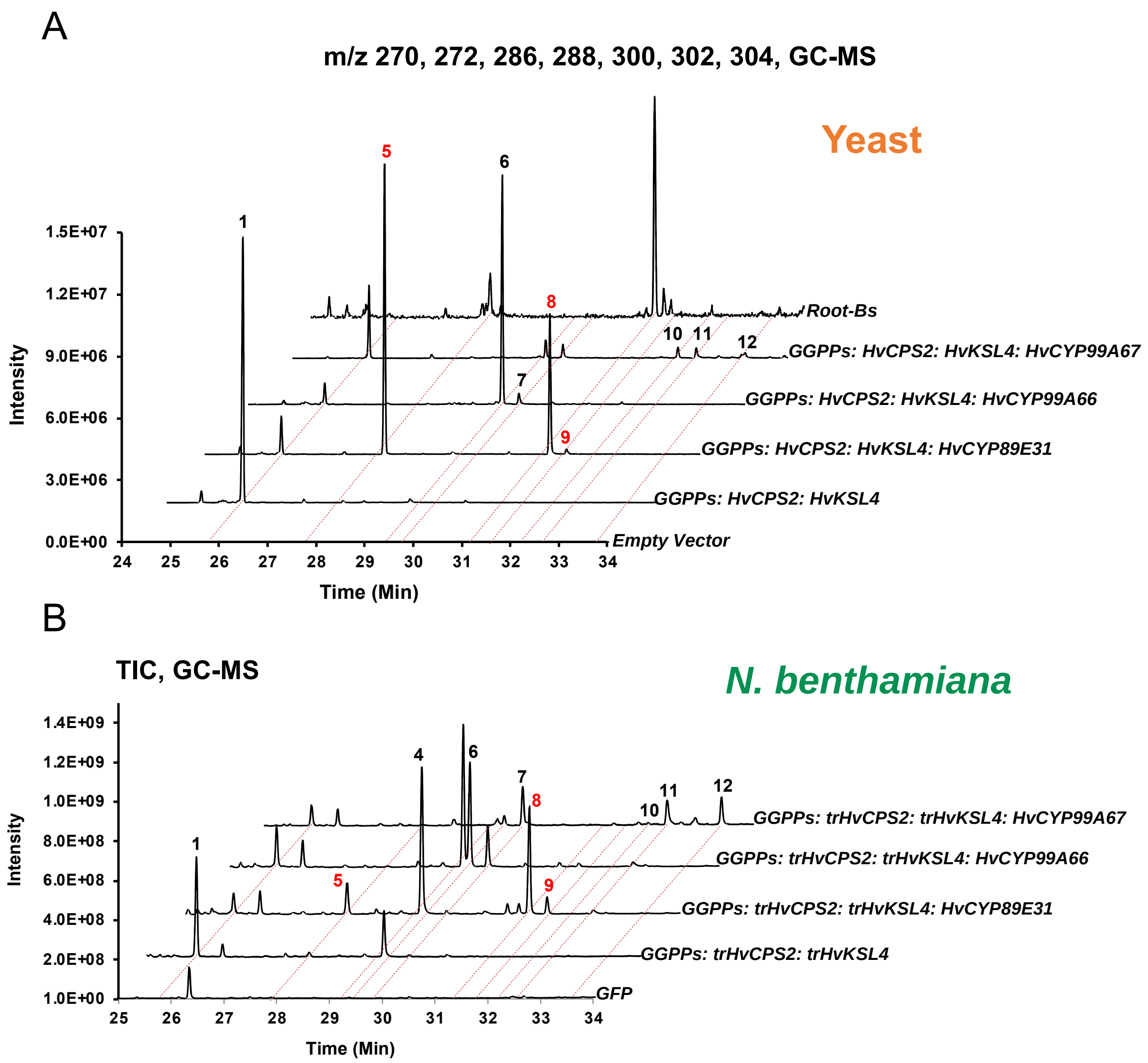

**Fig. S1: Characterization of HvCYP89E31, HvCYP99A66 and HvCYP99A67 in yeast and *N. benthamiana* (GC-MS).**

**A)** GC-MS analysis of products from yeast strains expressing the gene combinations indicated on the right. Selected ions, m/z 270, 272, 286, 288, 300, 302, 304.

**B)** GC-MS analysis of extracts from leaf discs expressing the gene combinations indicated on the right. TIC, Total ion chromatogram. Products that were also detected in barley roots are highlighted in red.

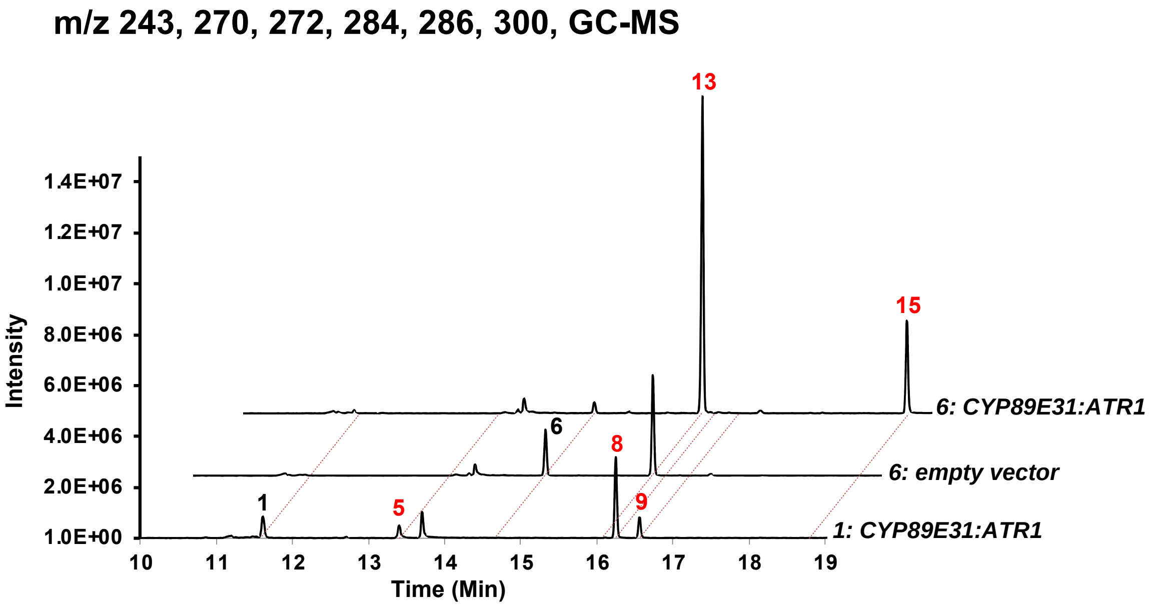

**Fig. S2: *In vitro* enzyme assays using microsomal preparation expressing CYP89E31 and ATR1.**

Microsomes expressing CYP89E31 and ATR1 and empty vector were incubated with **1** or **6**. Products were extracted then analysed by GC-MS. Selected ion (m/z 243, 270, 272, 284, 286, 300) chromatograms are presented. Converted products from **1** or **6** are highlighted in red.

A

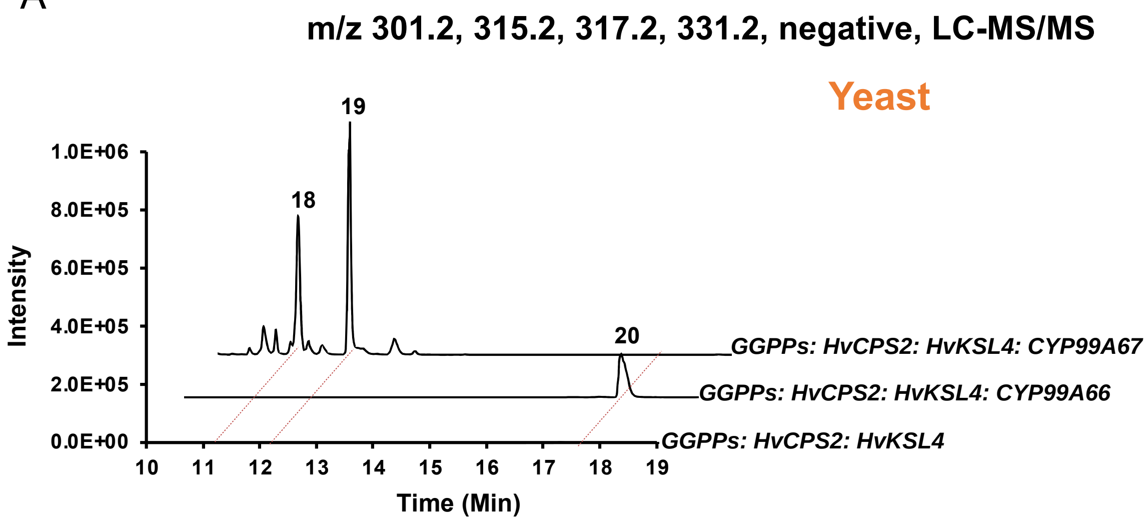

B

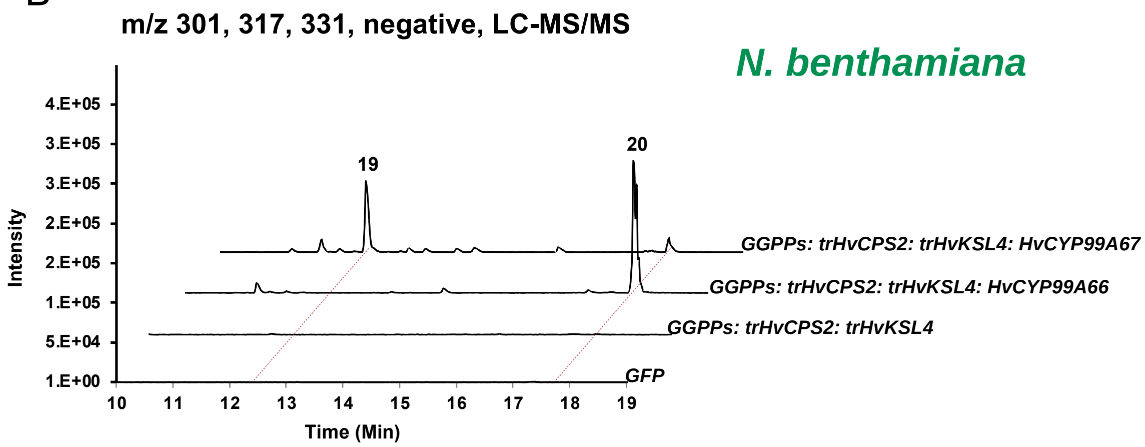

**Fig. S3: Characterization of HvCYP99A66 and HvCYP99A67 in yeast and *N. benthamiana* (LC-MS).**

**A)** LC-HRMS (negative) analysis of products from yeast strains expressing the gene combinations indicated on the right. Selected ions**,** m/z 301.2, 315.2, 317.2, 331.2.
**B)** LC-HRMS (negative) analysis of extracts from leaf discs expressing the gene combinations indicated on the right. Selected ions, m/z 301.2, 317.2, 331.2.
Products that were also detected in barley roots are highlighted in red.

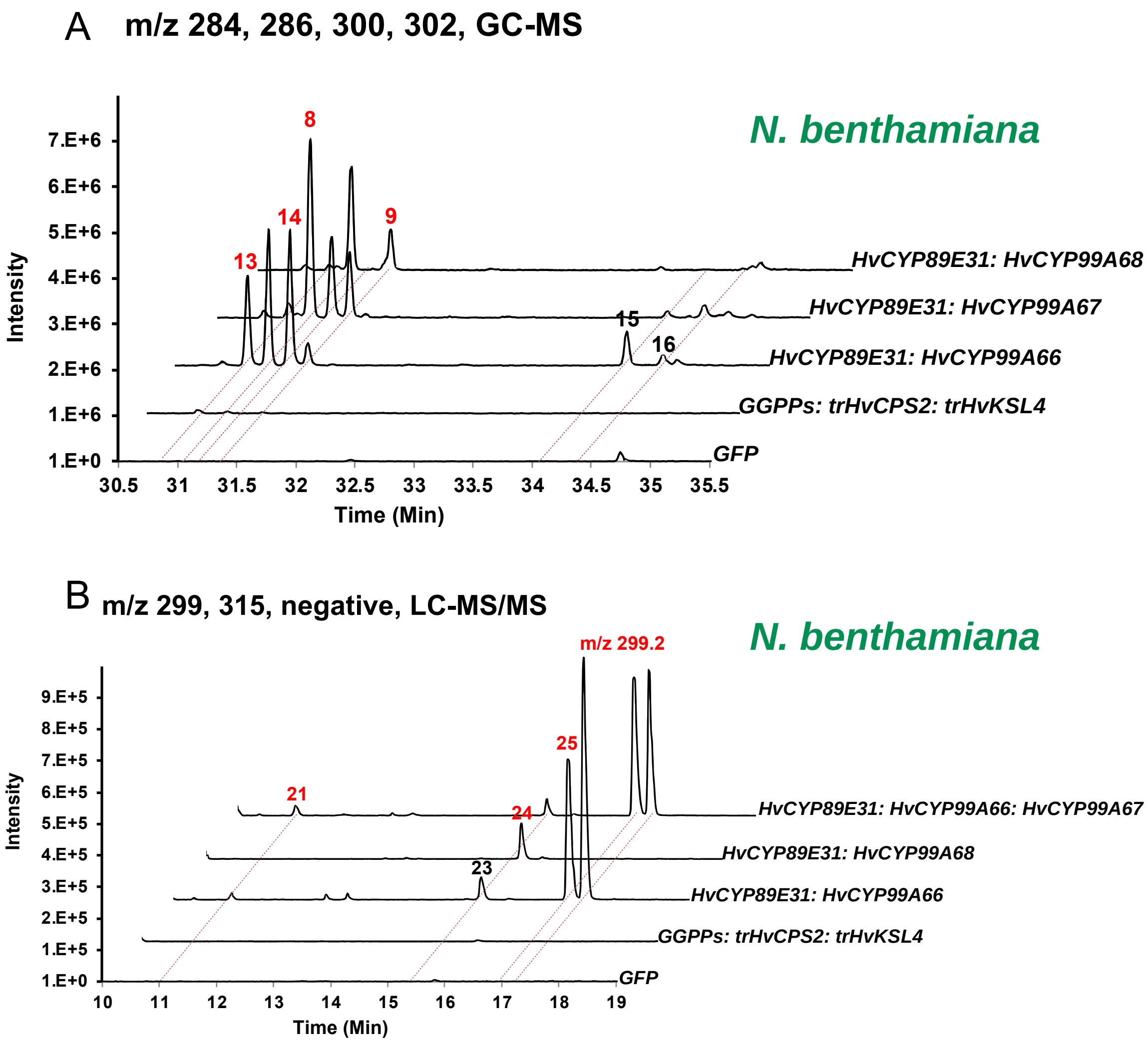

**Fig. S4: Co-expression of two or three CYPs in *N. benthamiana*.**

**A)** GC-MS analysis of extracts from leaf discs expressing the gene combinations indicated on the right. Selected ions, m/z 284, 286, 300, 302.

**B)** LC-HRMS (negative) analysis of extracts from leaf discs expressing the gene combinations indicated on the right. Selected ion, m/z 299.2, 315.2.

When CYPs were expressed, they were always co-expressed together with trHvCPS2, trHvKSL4 and GGPPs, though it was not written in the figure. Products that were also detected in barley roots are colored red.

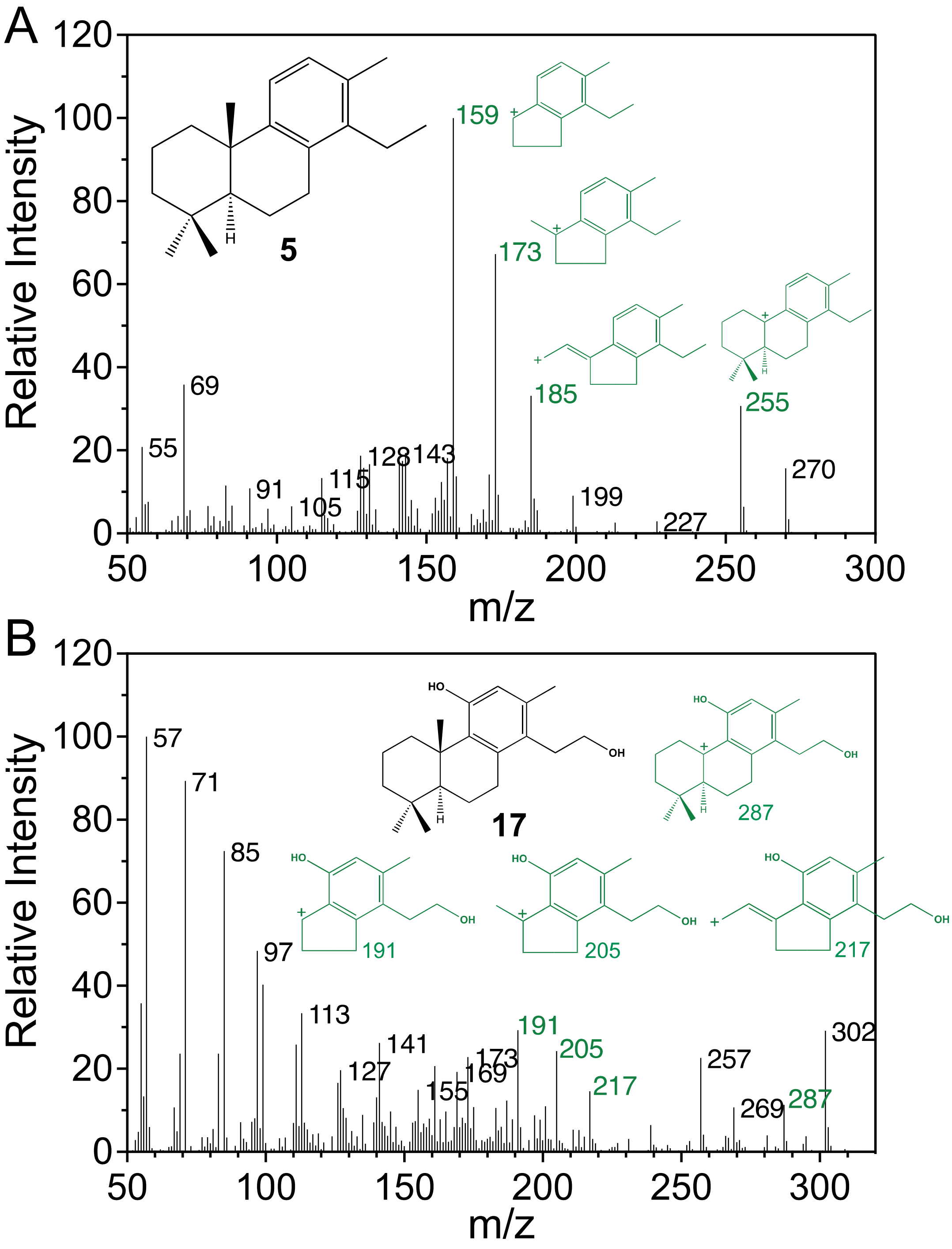

**Fig. S5: Fragment annotation of EI spectra of 5 and 17.**

A) EI spectrum of **5**, the annotation of fragments was recreated from (*17*).

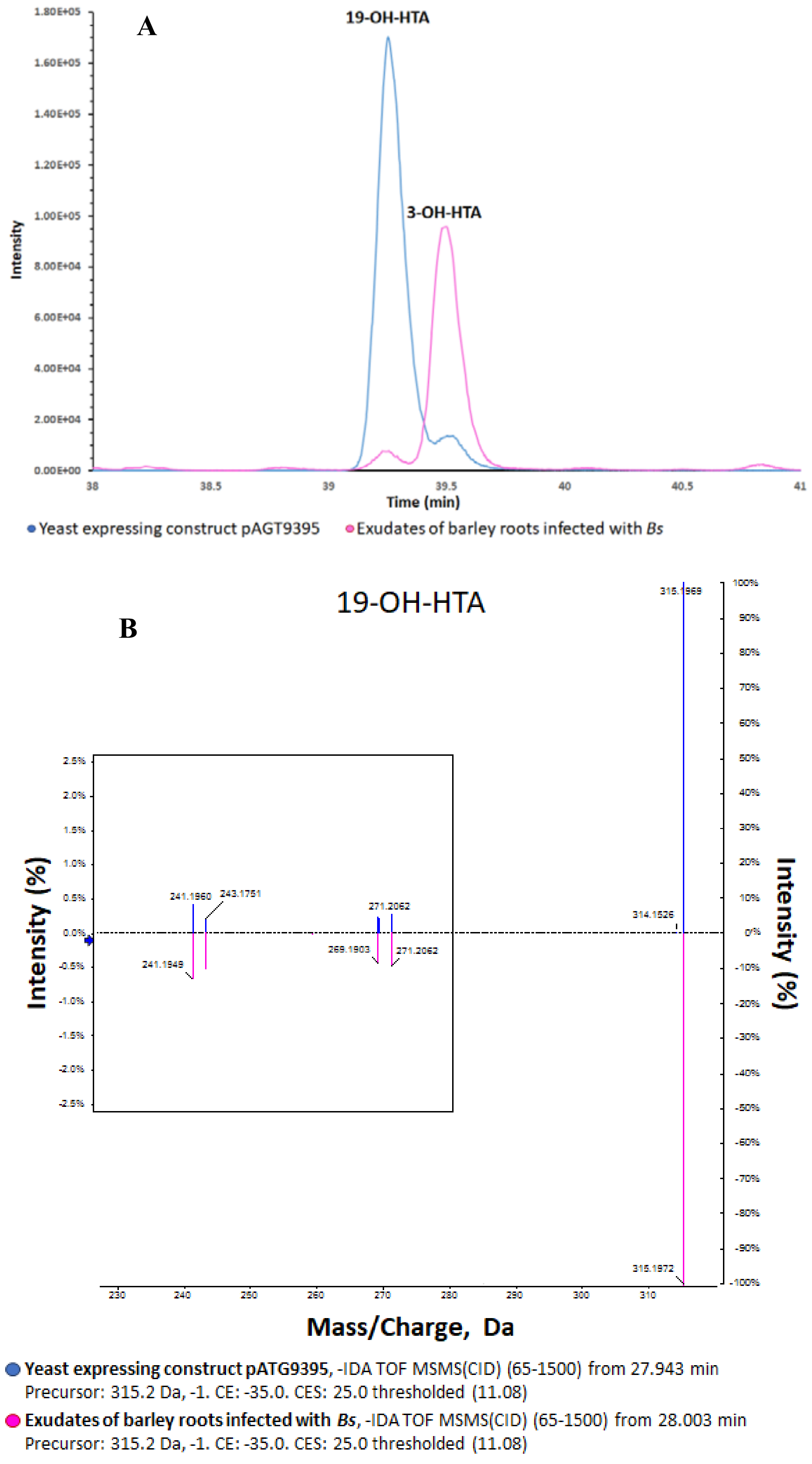
A
19-OH-HTA
3-OH-HTA
Intensity
Time (min)
Yeast expressing construct pAGT9395
Exudates of barley roots infected with *Bs*
B
19-OH-HTA
Intensity (%)
Intensity (%)
Mass/Charge, Da
241.1960
243.1751
241.1949
271.2062
269.1903
271.2062
314.1526
315.1968
315.1972
Yeast expressing construct pATG9395, -IDA TOF MSMS(CID) (65-1500) from 27.943 min
Precursor: 315.2 Da, -1. CE: -35.0. CES: 25.0 thresholded (11.08)
Exudates of barley roots infected with *Bs*, -IDA TOF MSMS(CID) (65-1500) from 28.003 min
Precursor: 315.2 Da, -1. CE: -35.0. CES: 25.0 thresholded (11.08)

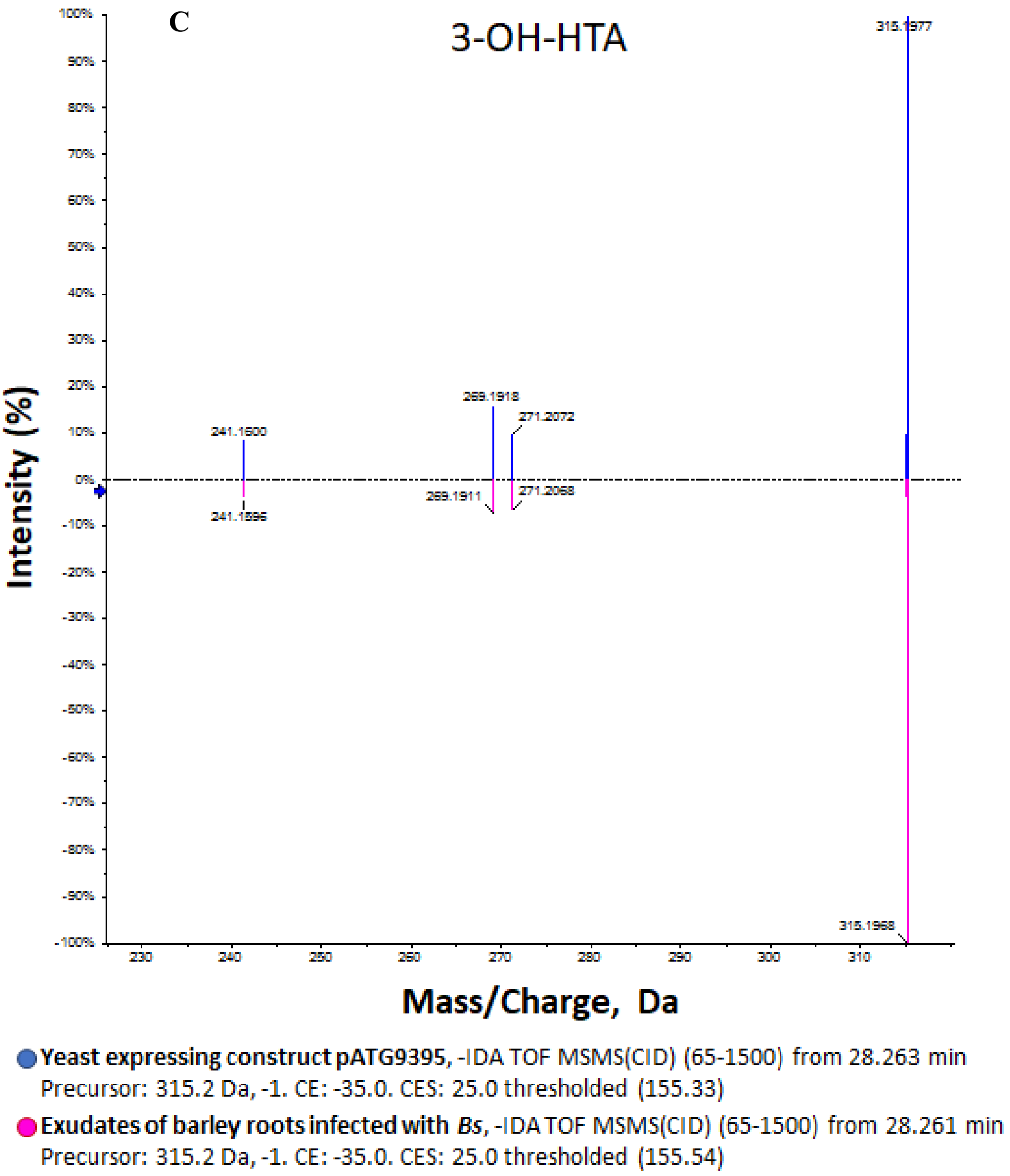

**Fig. S6: Separation of peak 21 in compound 21a and 21b**

**A)**. Using a longer gradient, two peaks could be separated under peak 21. The first peak (compound **21a,** NMR data **Table S10**) is the major product produced in yeast with plasmid pAGT9593 expressing CYP89E31, CYP99A66 and CYP99A67 in addition to GGPPS, HvCPS2 and HvKSL4. Compound **21a** is 19-β-hydroxy-hordetrien-17-oic acid (19-OH-HTA). The second peak (compound **21b**) is the major product found in exudates barley roots infected with *B. sorokiniana*. Compound **21b** is 3-β-hydroxy-hordetrien-17-oic acid (3-OH-HTA, NMR data Table S11). **B)** MS/MS spectra of compound **21a** produced in yeast (top panel, blue) and from barley root exudates (bottom panel, pink). **C)** MS/MS spectra of compound **21b** produced in yeast (top panel blue) and from barley root exudates (bottom panel, pink).

**A**

| ***HvCPS2*-sgRNA1** | |
|---|---|
| WT | CCATAC–CAACGACCCTACTCTTCA |
| *cps2-1* | CCATACTCAACGACCCTACTCTTCA |
| *cps2-2* | CCATACTCAACGACCCTACTCTTCA |

| ***HVCPS2*-sgRNA2** | |
|---|---|
| WT | CCATGT–GGCTTCGAGATTAACTTC |
| *cps2-1* | CCATGTGGGCTTCGAGATTAACTTC |
| *cps2-2* | CCATGT–GGCTTCGAGATTAACTTC |

| ***HvKSL4*-sgRNA** | |
|---|---|
| WT | CCGGTG–AATTTGACTCACCAGCCA |
| *ksl4-1* | CCGGTGAAATTTGACTCACCAGCCA |
| *ksl4-2* | CCGGTGTAATTTGACTCACCAGCCA |

**B**

```
             1 ........10........20........30........40........50........60........70
HvCPS2       1 MLTFTAAFRHVPVLDHPTAEPWRRLSLHLHSQHRRCGVVLSSKSPPYPDVEVGERKVHEYIRHTDEPSEM
cps2-1       1 MLTFTAAFRHVPVLDHPTAEPWRRLSLHLHSQHRRCGVVLSSKSPPYPDVEVGERKVHEYIRHTDEPSEM
cps2-2       1 MLTFTAAFRHVPVLDHPTAEPWRRLSLHLHSQHRRCGVVLSSKSPPYPDVEVGERKVHEYIRHTDEPSEM

            71 ........80........90.......100.......110.......120.......130.......140
HvCPS2      71 RQMIDAIRTTLASLGDDETSMSVSAYDTALVALVKNLDGGDGPQFTSCIDWIVQNQLPDGSWGDPDFFMV
cps2-1      71 RQMIDAIRTTLASLGDDETSMSVSAYDTALVALVKNLDGGDGPQFTSCIDWIVQNQLPDGSWGDPDFFMV
cps2-2      71 RQMIDAIRTTLASLGDDETSMSVSAYDTALVALVKNLDGGDGPQFTSCIDWIVQNQLPDGSWGDPDFFMV

           141 .......150.......160.......170.......180.......190.......200.......210
HvCPS2     141 QDRMISTLACVVALKSWNIDIDNLCDRGMLFIKENMSRLLEQEQDWMPCGFEINFPALLEKAKDLDLDIP
cps2-1     141 QDRMISTLACVVALKSWNIDIDNLCDRGMLFIKENMSRLLEQEQDWMPCGLRD*----------------
cps2-2     141 QDRMISTLACVVALKSWNIDIDNLCDRGMLFIKENMSRLLEQEQDWMPCGFEINFPALLEKAKDLDLDIP

           211 .......220.......230.......240.......250.......260.......270.......280
HvCPS2     211 YDHPVLEEIYAKKNLKLSKIPLNVLHAIPTTLLFSLEGMVDLPLDWEKLLRLRCTDGSFHSSPAATAAAL
cps2-1         ----------------------------------------------------------------------
cps2-2     211 YDHPVLEEIYAKKNLKLSKIPLNVLHAILN-------------------------DPTLQP*--------

           281 .......290.......300.......310.......320.......330.......340.......350
HvCPS2     281 SRTGDKECQAFLDRLIKKFDGGVPCSHSMDTFEQVWVVDRLMHLGISRHFTSEIDQFLEFIYRRWTNKGL
cps2-1         ----------------------------------------------------------------------
cps2-2         ----------------------------------------------------------------------

           351 .......360.......370.......380.......390.......400.......410.......420
HvCPS2     351 AHNVHCPIADIDDTAMGFRLLRQHGYEVNPSVFKQFEKDGRFVCFPMETNHASVTPMHNTYRASQFMFPG
cps2-1         ----------------------------------------------------------------------
cps2-2         ----------------------------------------------------------------------

           421 .......430.......440.......450.......460.......470.......480.......490
HvCPS2     421 DDDVLARAGRYCRAFLEERQASNNLYDKWIITKDLPGEVGYTLNFPWKASLPRIETRMYLDQYGGNTDVW
cps2-1         ----------------------------------------------------------------------
cps2-2         ----------------------------------------------------------------------

           491 .......500.......510.......520.......530.......540.......550.......560
HvCPS2     491 IAKVLYRMNLVSNDLYLKMAKADFREYQRLSRLEWNGLRKWYFRNHLQRYGGTPKSALTAYFLASANIFE
cps2-1         ----------------------------------------------------------------------
cps2-2         ----------------------------------------------------------------------

           561 .......570.......580.......590.......600.......610.......620.......630
HvCPS2     561 PGRATERLAWARMAVLAEAVTTHFRHIGGPCNSTENLEDLIDLVSFDDVSGGLREAWKQWLMAWTAKESH
cps2-1         ----------------------------------------------------------------------
cps2-2         ----------------------------------------------------------------------

           631 .......640.......650.......660.......670.......680.......690.......700
HvCPS2     631 GSIDGDTALLFVRTIEIVSGRNVAAEKKLNLWDYSQLEQLTSSICHKLATIGLAQNGGSMENTEDLQRQV
cps2-1         ----------------------------------------------------------------------
cps2-2         ----------------------------------------------------------------------

           701 .......710.......720.......730.......740.......750.......
HvCPS2     701 DLEMQELSWRIHQGCQGINRETRETFLNVVKSFYYSAHCSPETVDSHIAKVIFQDVI
cps2-1         ---------------------------------------------------------
cps2-2         ---------------------------------------------------------
```

**C**

```
             1 ........10........20........30........40........50........60........70
HvKSL4       1 MSTAKVLLPVPKSVSHHHQRNPLRLAFAHGSTEAFFPPSIRRRPSLHSIRTRSRKISACTAYVESRPLGE
ksl4-1/2     1 MSTAKVLLPVPKSVSHHHQRNPLRLAFAHGSTEAFFPPSIRRRPSLHSIRTRSRKISACTAYVESRPLGE

            71 ........80........90.......100.......110.......120.......130.......140
HvKSL4      71 RNVNRQNMDREARIRKQLQNPKFSPSPYDTAWVAMVPLPGSLQDPCFPQCVEWILQNQHGNGYWGSGEFD
ksl4-1/2    71 RNVNRQNMDREARIRKQLQNPKFSPSPYDTAWVAMVPLPGSLQDPCFPQCVEWILQNQHGNGYWGSGEI*

           141 .......150.......160.......170.......180.......190.......200.......210
HvKSL4     141 SPASRDVLLSTLASVIALKKWNVGPEEIMRGLQFIGRNFSIIMDEQTTAPIGYNLTFSSLLILAIEMGLE
ksl4-1/2   140 ----------------------------------------------------------------------

           211 .......220.......230.......240.......250.......260.......270.......280
HvKSL4     211 LPVSHADINVIVRHREMEIERLDAEKSSTKEVYSSYVAEGLVNVLDLSEVMTFQKKNGSLFNSPSATAAA
ksl4-1/2       ----------------------------------------------------------------------

           281 .......290.......300.......310.......320.......330.......340.......350
HvKSL4     281 LIRNYDLGAFEYLNLIVSEFGSAVPAMYPLTVHYQLSMVDTLQKVGISRFFSSEINHILDKTYSLLLKRD
ksl4-1/2       ----------------------------------------------------------------------

           351 .......360.......370.......380.......390.......400.......410.......420
HvKSL4     351 DEIMSNVETCAVAFRILRMNRYDVSSDVLSHVAEASTLLDPPQEYVTDTKSLLELYKASKVILSQNELVL
ksl4-1/2       ----------------------------------------------------------------------

           421 .......430.......440.......450.......460.......470.......480.......490
HvKSL4     421 EKIEKWSRSLLKEIMCSDMAQRIPAVIEEVEYALKIPFYATVDPLDHKWSIEHFDAMDSHMMKTKYIIRS
ksl4-1/2       ----------------------------------------------------------------------

           491 .......500.......510.......520.......530.......540.......550.......560
HvKSL4     491 CGVNKDILALAVEDFCVSQSIYQREVEHLDSWEKECRLGELQFARQKMKYCYLCAAASITPHELSEARVA
ksl4-1/2       ----------------------------------------------------------------------

           561 .......570.......580.......590.......600.......610.......620.......630
HvKSL4     561 CAKATILTIVIDDFFDSGAGSPEALANLISLAEKWEDPHEDDFHSEEVKILFYALYKTMNQIAAAASPLQ
ksl4-1/2       ----------------------------------------------------------------------

           631 .......640.......650.......660.......670.......680.......690.......700
HvKSL4     631 NRDVTKELVETWLALMRTEMTEVEWRNSKHLPSFEEYMKVAYVSFALGPIVHTPMYFLGVNFPEHVLRDE
ksl4-1/2       ----------------------------------------------------------------------

           701 .......710.......720.......730.......740.......750.......760.......770
HvKSL4     701 EYDELFTIMSSCCRLLNDIRGFERELSHGKLNSTLLLVRHSGGSLSIEEAKQEIQRSIASLTTDLLRLLL
ksl4-1/2       ----------------------------------------------------------------------

           771 .......780.......790.......800.......810.......820.......830.....
HvKSL4     771 REDKVVPMPCKEIFWRFNQTGHLFYTRIDGFTSPEEMVGAVNAVVYDPLKLQGGTVPSLSAQLEN
ksl4-1/2       -----------------------------------------------------------------
```

**Fig. S7: Generation and sequence of *cps2* and *ksl4* mutants.**

**A)** DNA sequences of *HvCPS2* or *HvKSL4* in wild-type or mutant plants. Mutation in the DNA sequence is colored in red. PAM sequence recognized by Cas9 is colored in green. **B**) Alignment of protein sequences of *HvCPS2*, and *cps2* mutants. Amino acids identical among the three protein sequences are shaded in black. The DxDD motif that is critical for the cyclization of GGPP (*18*) is highlighted in red. **C)** Alignment of protein sequences of *HvKSL4*, and *ksl4* mutants. The amino acid that is identical between the two protein sequences is shaded in black. The DDxxD motif that is conserved for class I diterpene synthase (*18*) is highlighted in red.

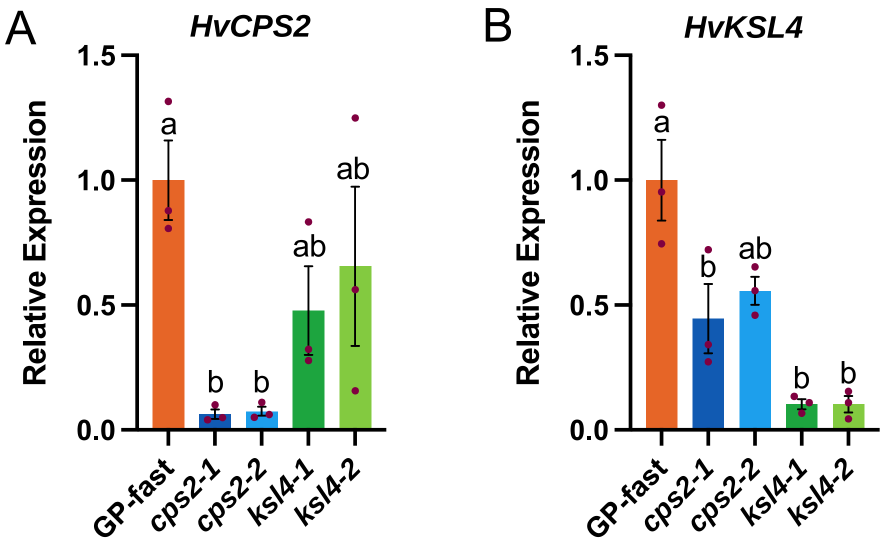

**Fig. S8: Gene expression of *HvCPS2* and *HvKSL4* in the corresponding mutants.**
**A)** and **B)** Quantification of the expression of *HvCPS2* and *HvKSL4* respectively in wild-type barley and mutants by qRT-PCR. Barley *ubiquitin* (*UBI*) was used as the reference gene. All expression data were normalized to that of GP-fast. Letters represent statistically significant differences using one-way ANOVA and Tukey's post-hoc test. Bars, Mean ± SEM (n=3).

**Fig. S9: MS or MS/MS spectra of terpenoids shown in this research and their structures or putative structures.**

**1**

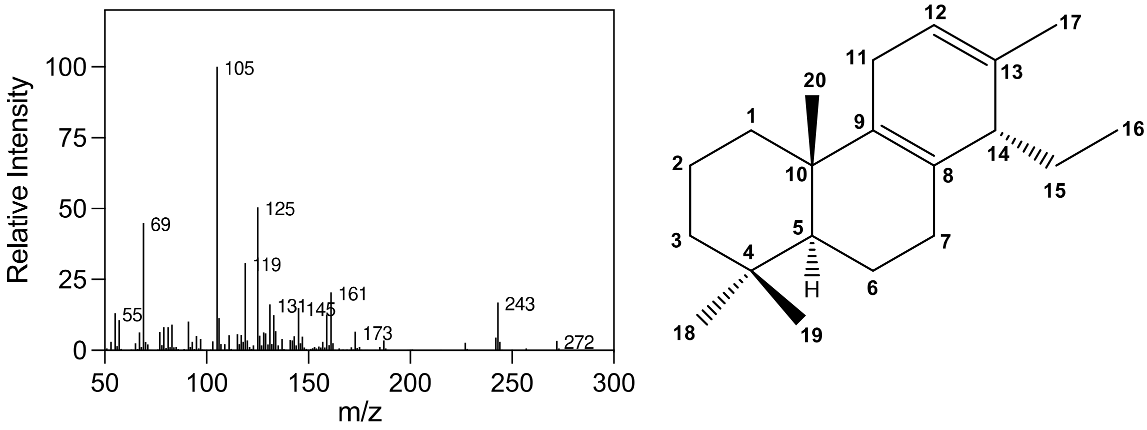

**2**

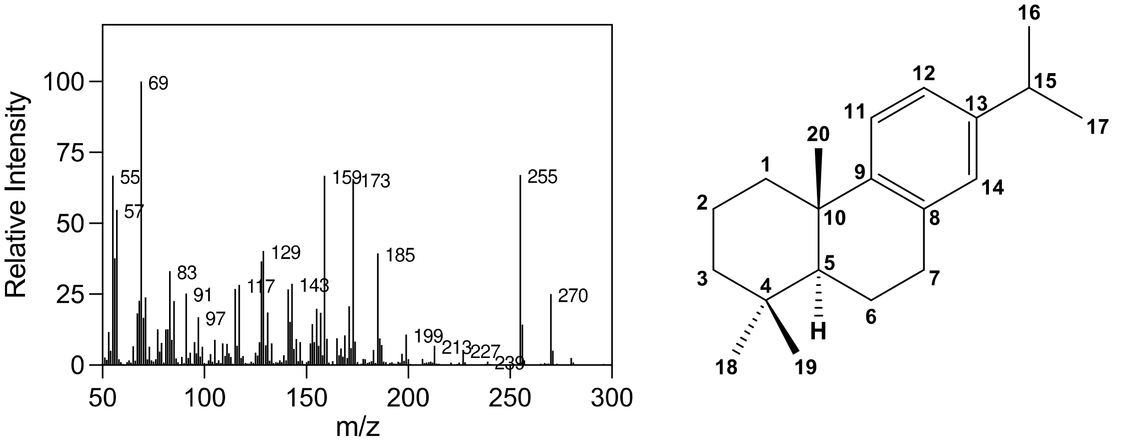

**3**

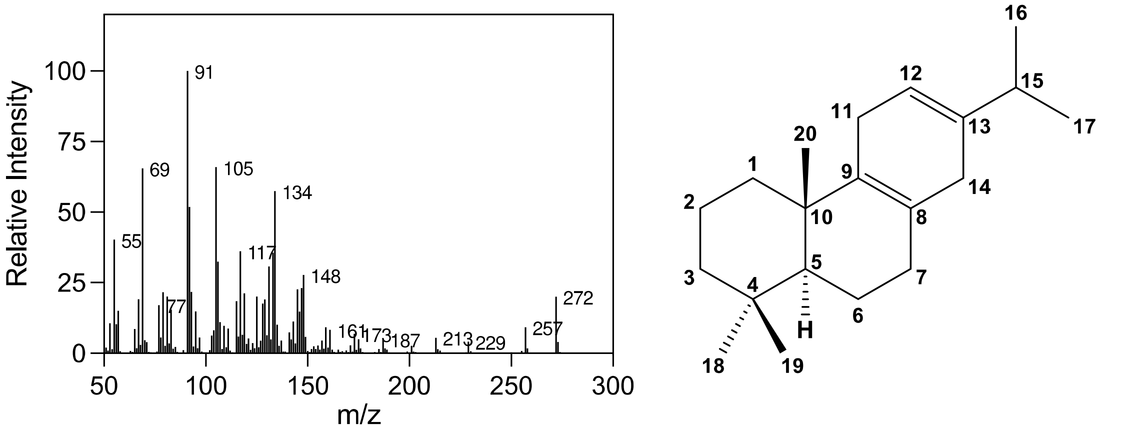

# 4

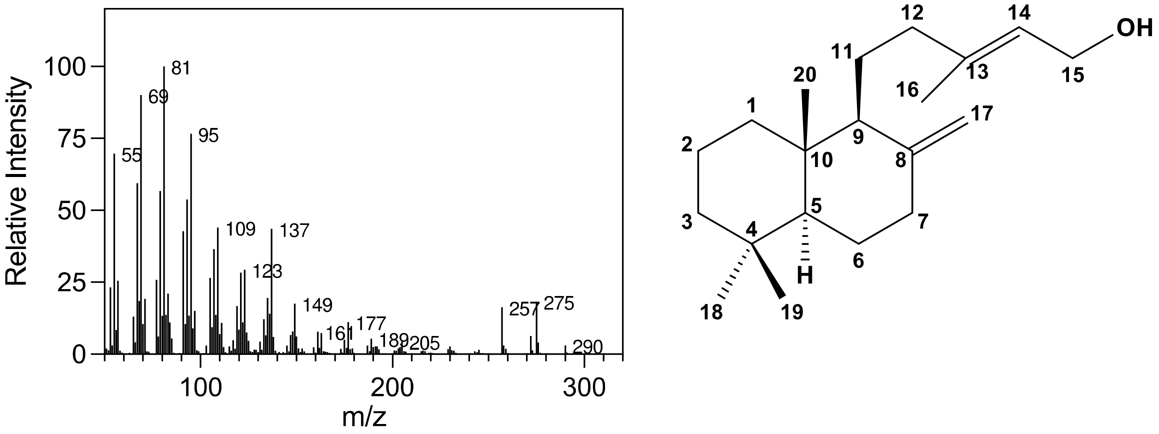
Relative Intensity
m/z
81
69
95
55
109
137
123
149
257
275
177
161
189
205
290
OH

# 5

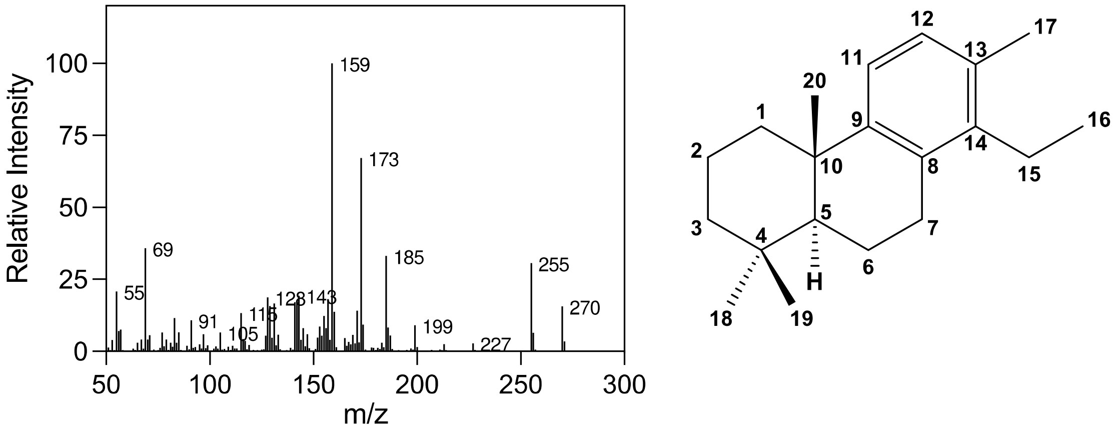
Relative Intensity
m/z
159
173
69
185
255
55
270
91
105
199
227

# 6

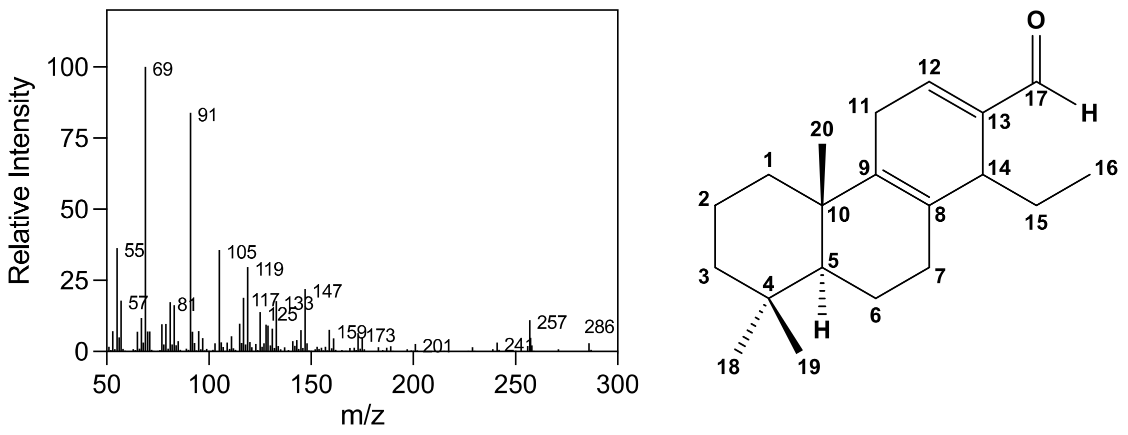
Relative Intensity
m/z
69
91
55
105
119
57
81
117
125
133
147
159
173
201
241
257
286
O
H

# 7

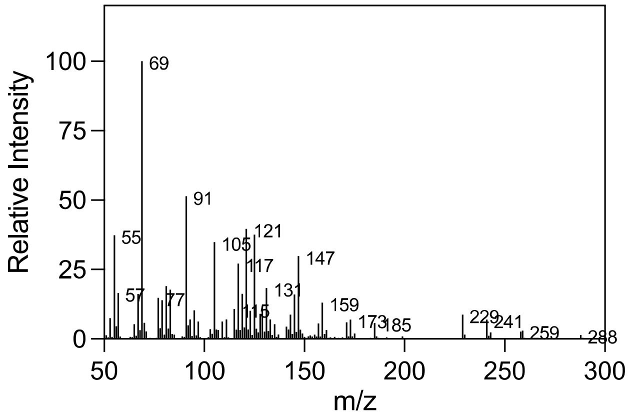

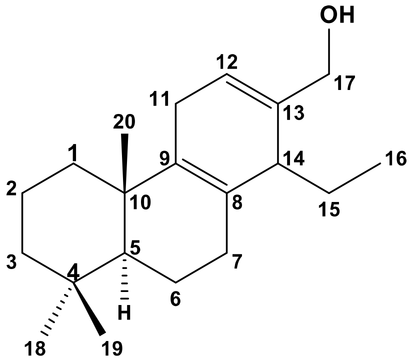

**Putative, NO NMR**

# 8

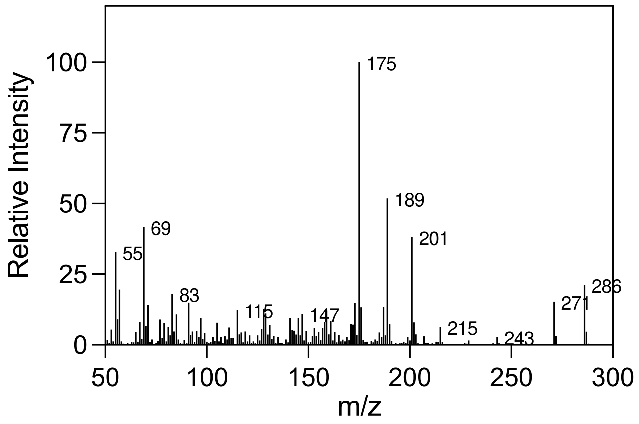

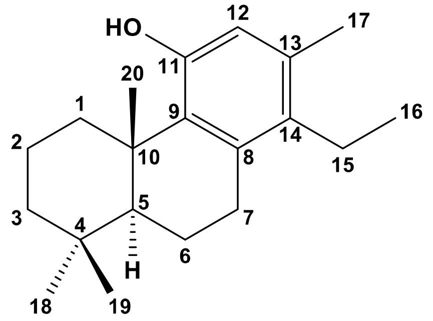

# 9

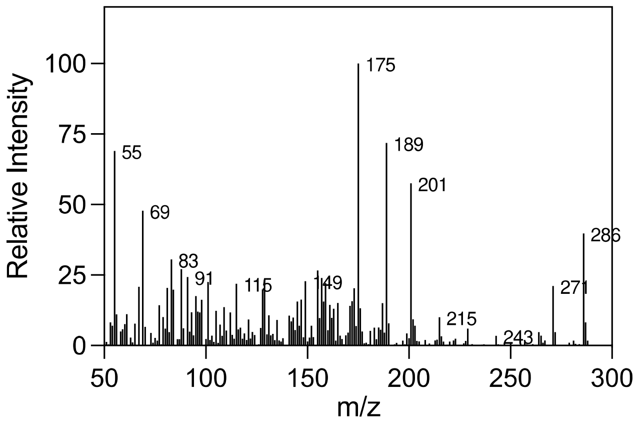

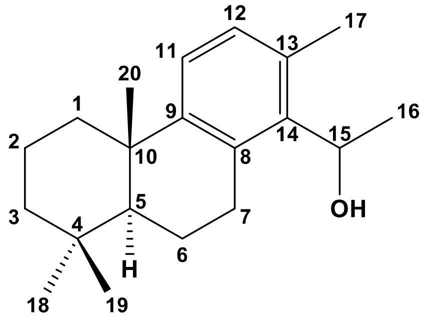

**Putative, NO NMR**

# 10

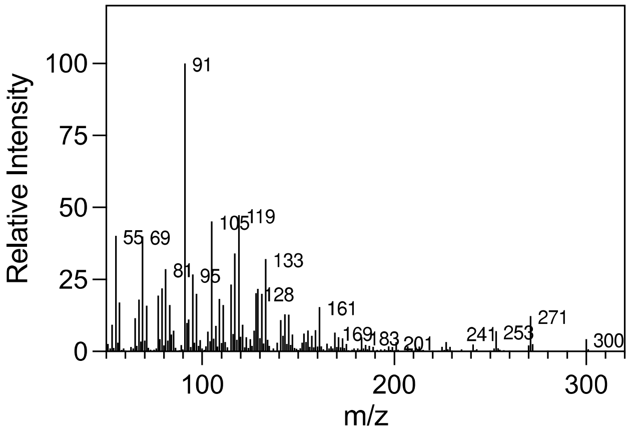

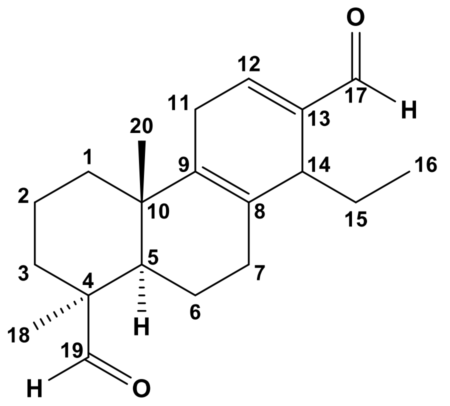

**Putative, NO NMR**

# 11

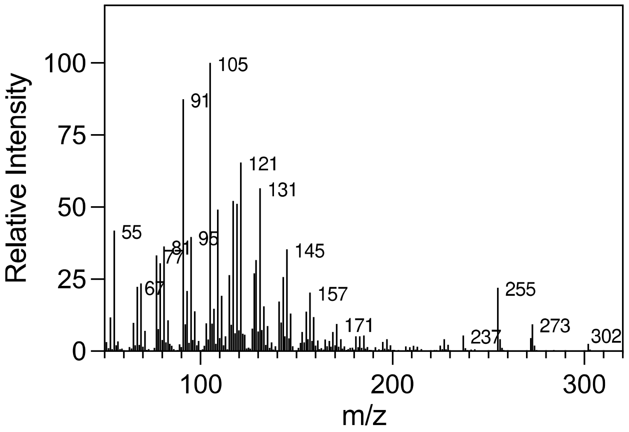

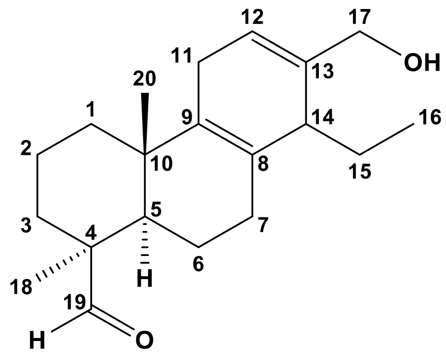

**Putative, NO NMR**

# 12

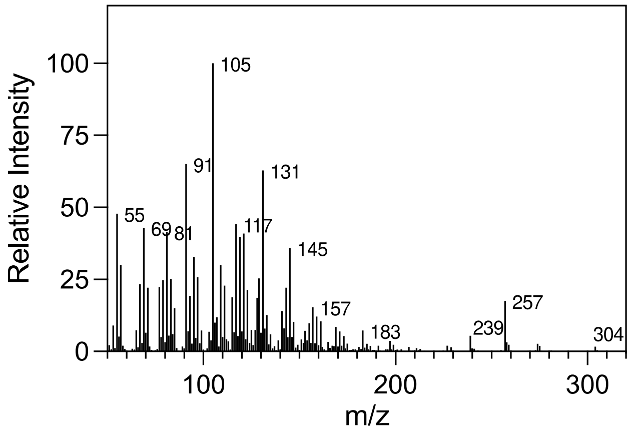

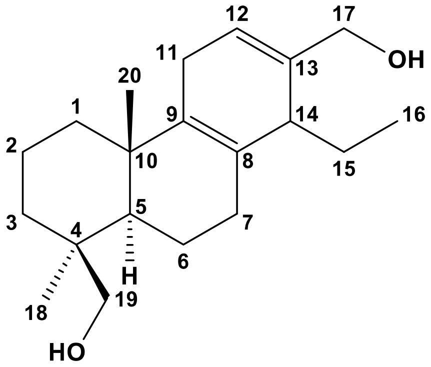

**Putative, NO NMR**

# 13

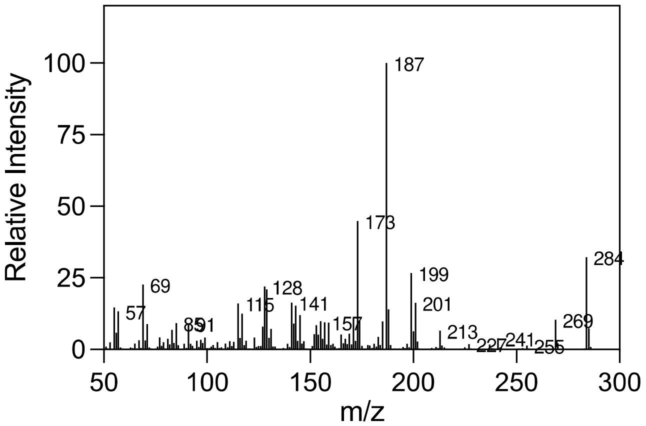

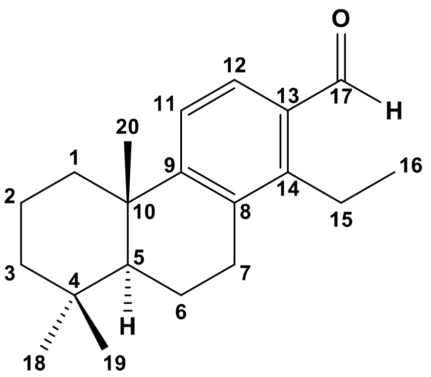

**Putative, NO NMR**

# 14

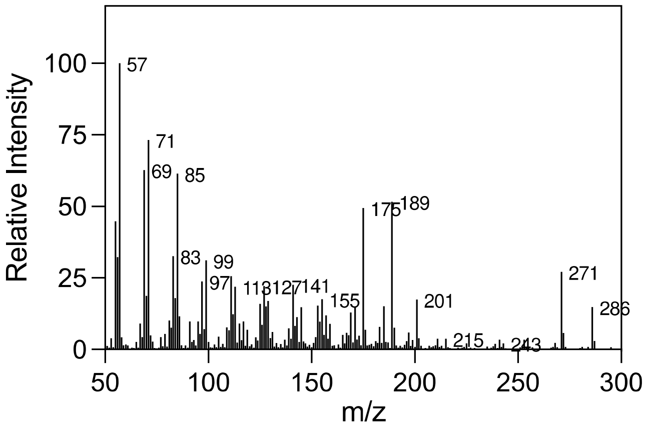

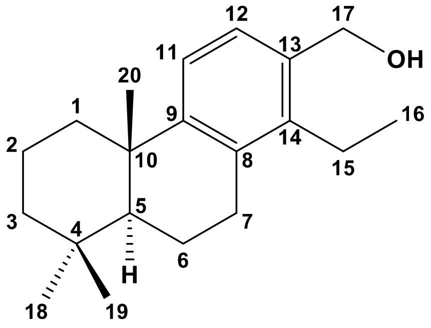

**Putative, NO NMR**

# 15

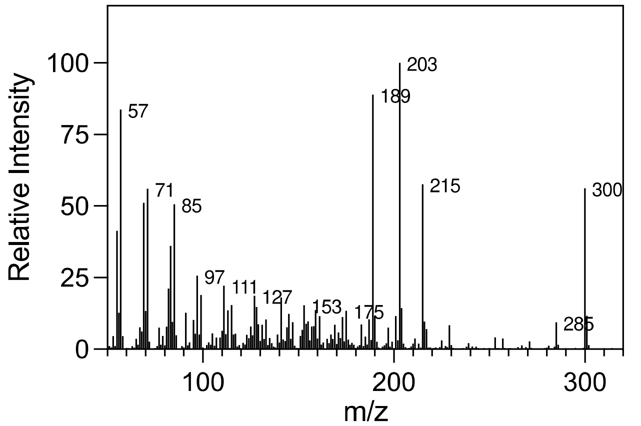

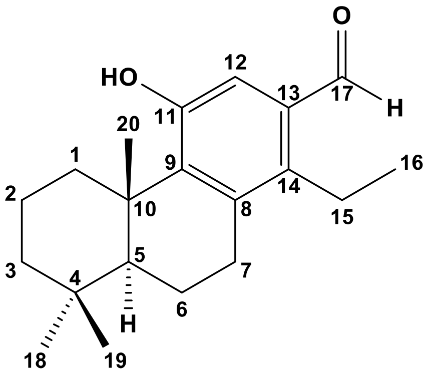

**Putative, NO NMR**

# 16

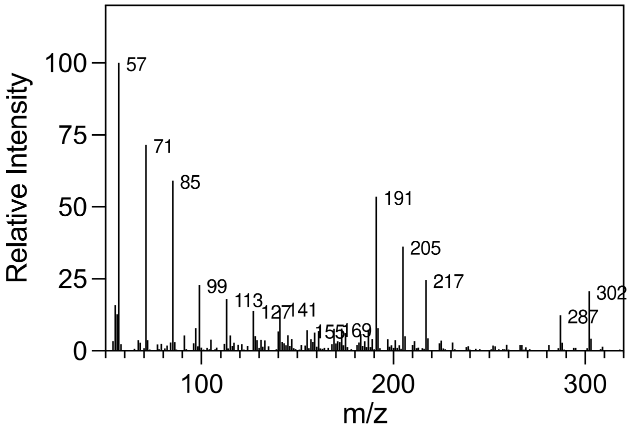

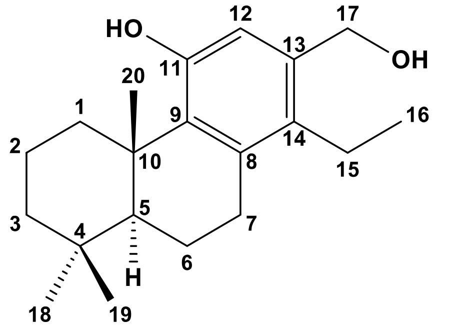

**Putative, NO NMR**

# 17

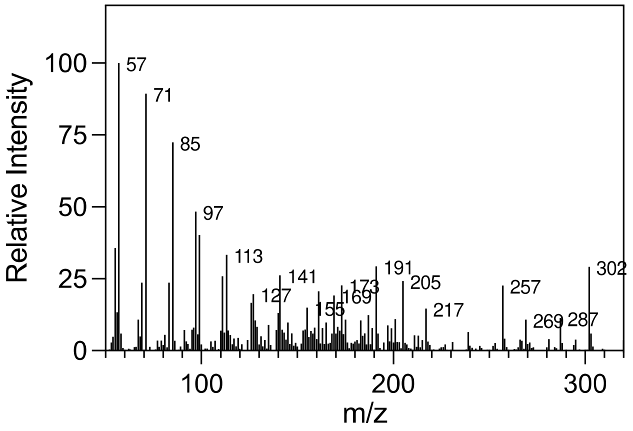

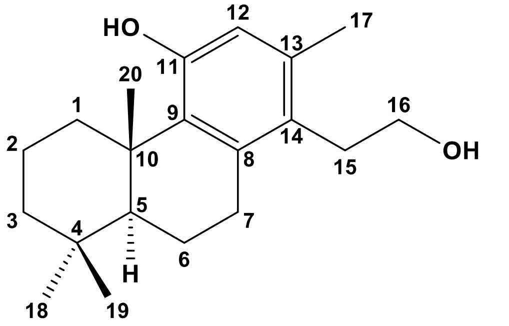

**Putative, NO NMR**

# 18

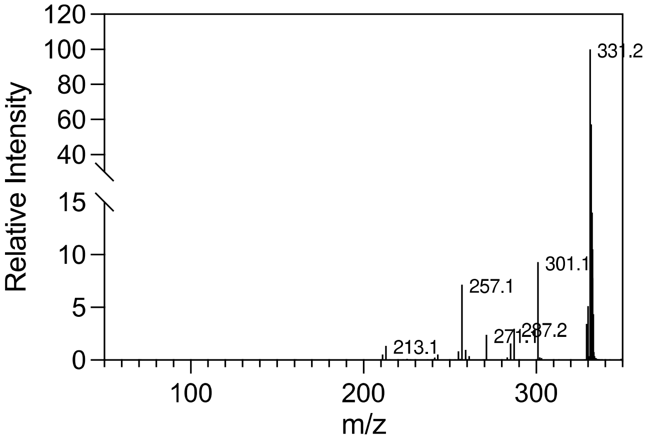

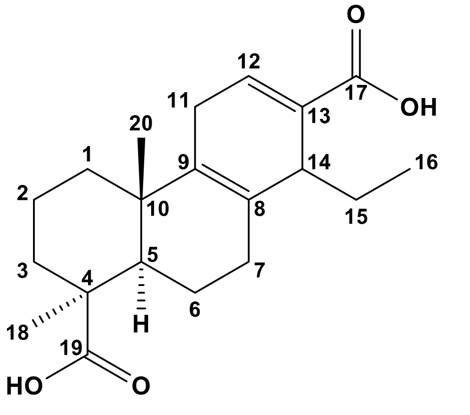

**Putative, NO NMR**

# 19

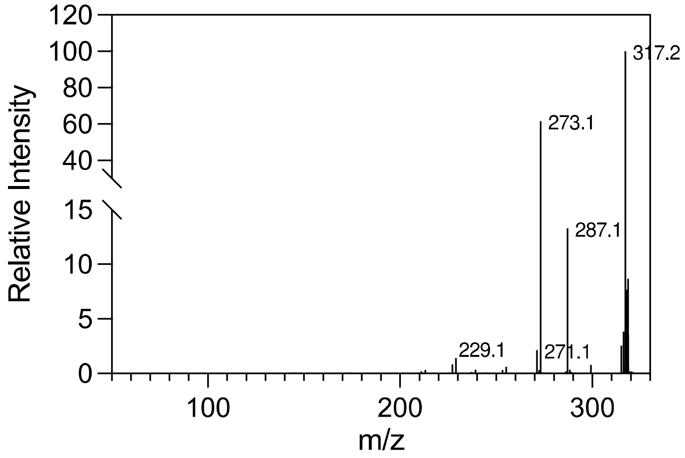

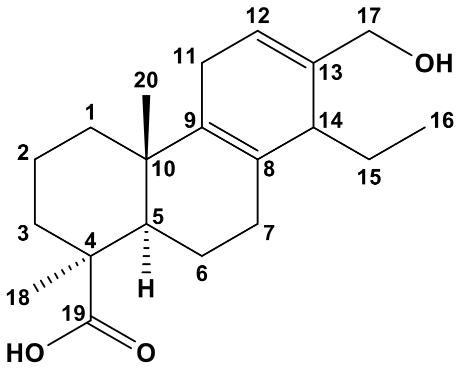

# 20

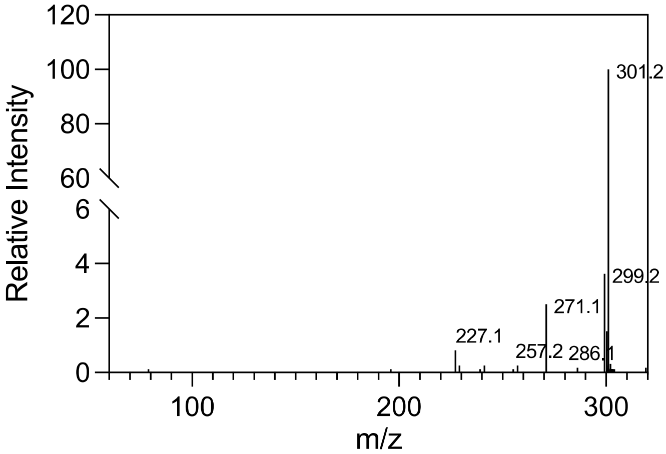

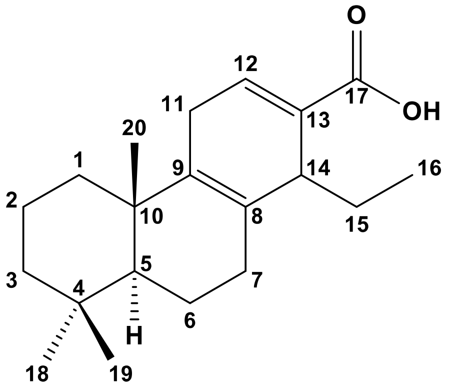

**Putative, NO NMR**

# 21

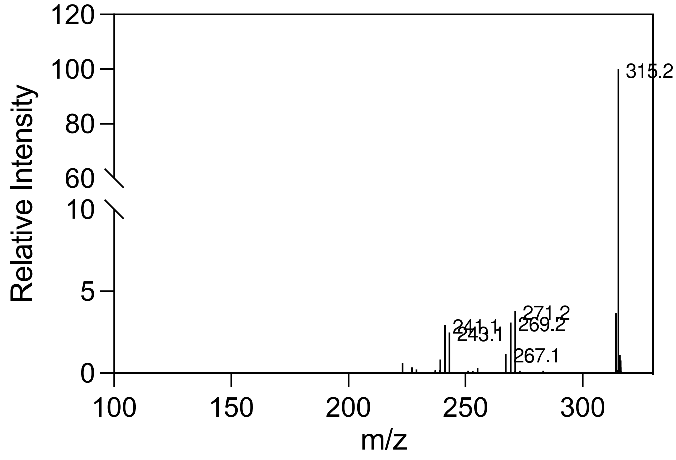

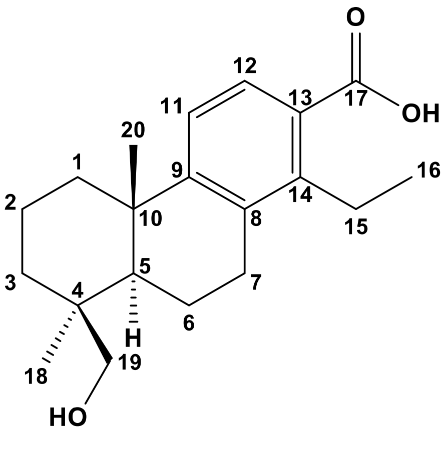

## 22

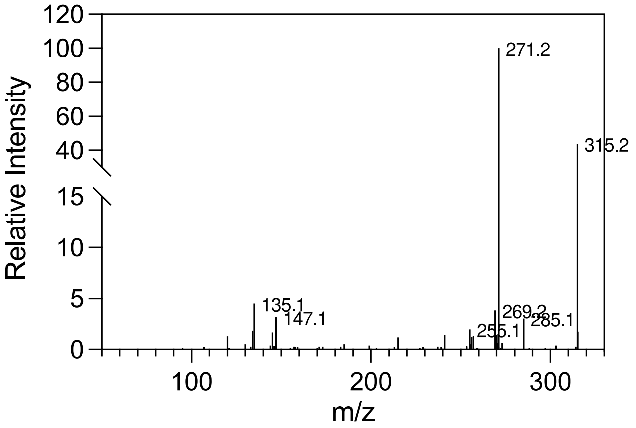

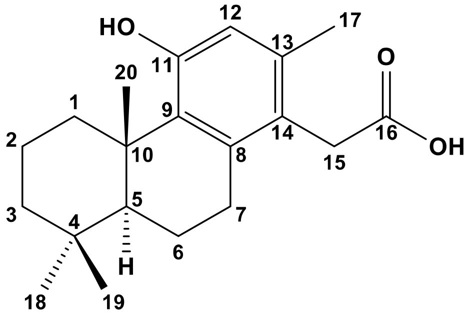

**Putative, NO NMR**

## 23

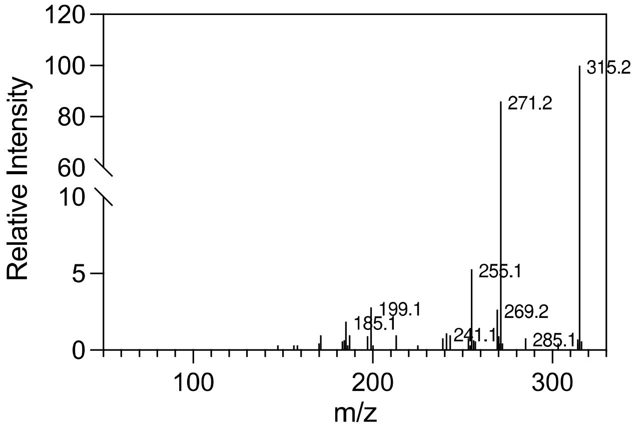

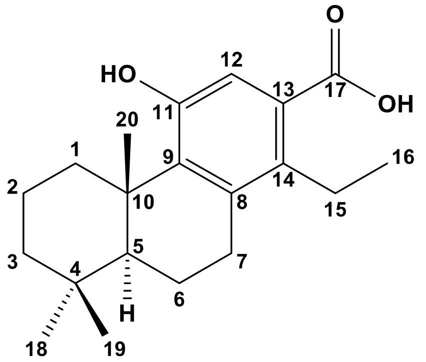

**Putative, NO NMR**

## 24

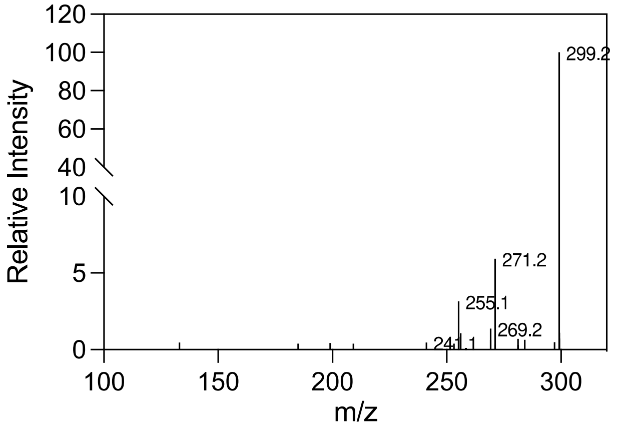

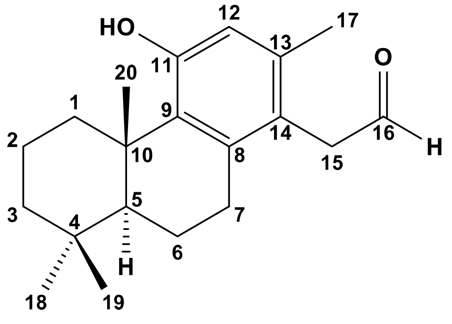

**Putative, NO NMR**

**25**

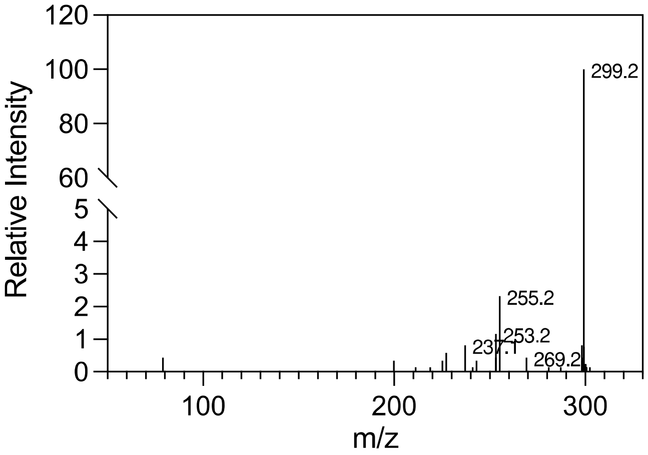

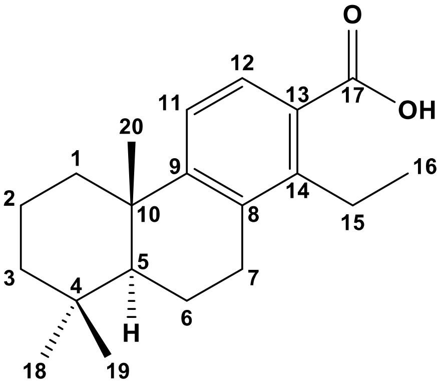

**Putative, NO NMR**

**26**

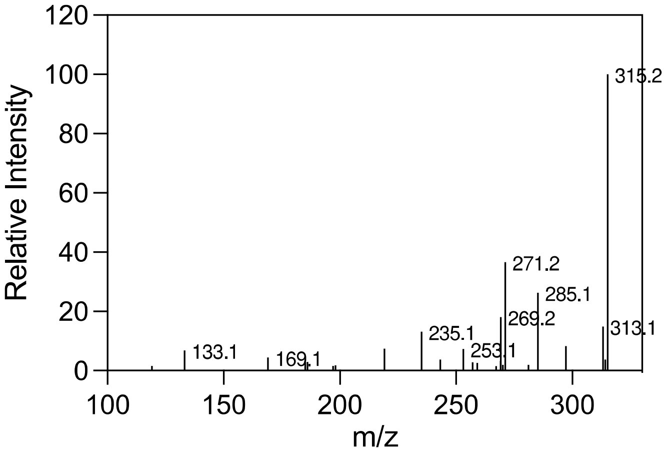

**27**

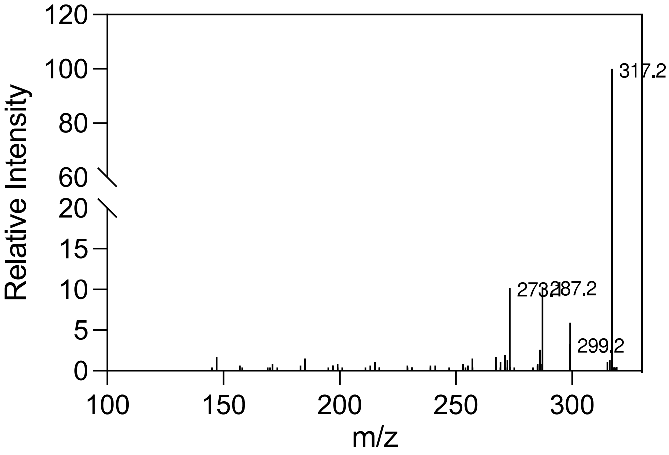

**28**

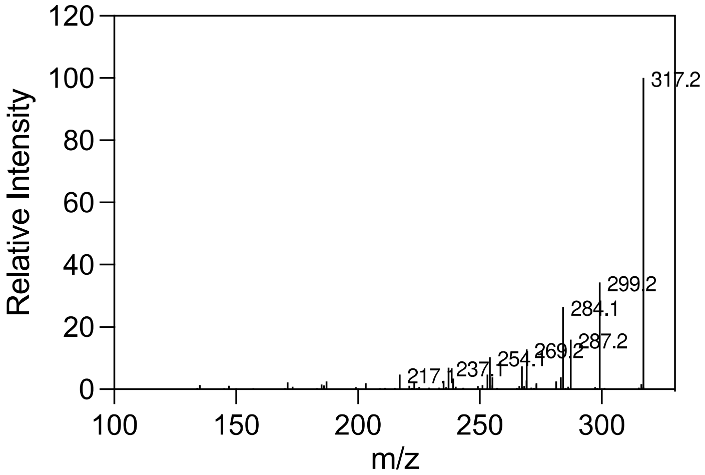

Relative Intensity
m/z
317.2
299.2
284.1
287.2
269.2
254.1
237.1
217.1

**29**

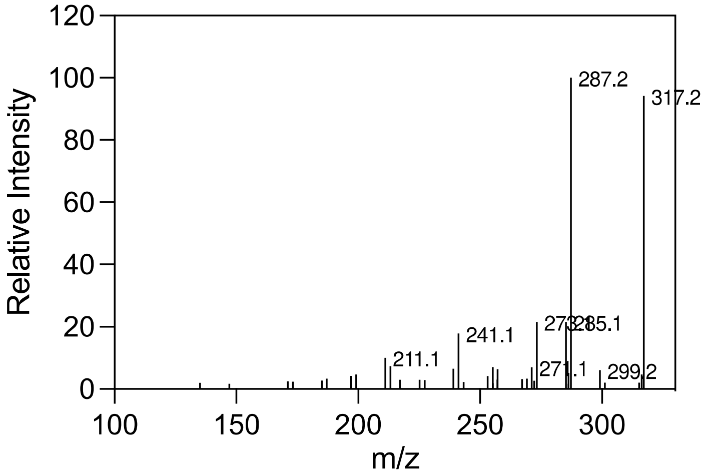

Relative Intensity
m/z
287.2
317.2
241.1
211.1
285.1
271.1
299.2

**30**

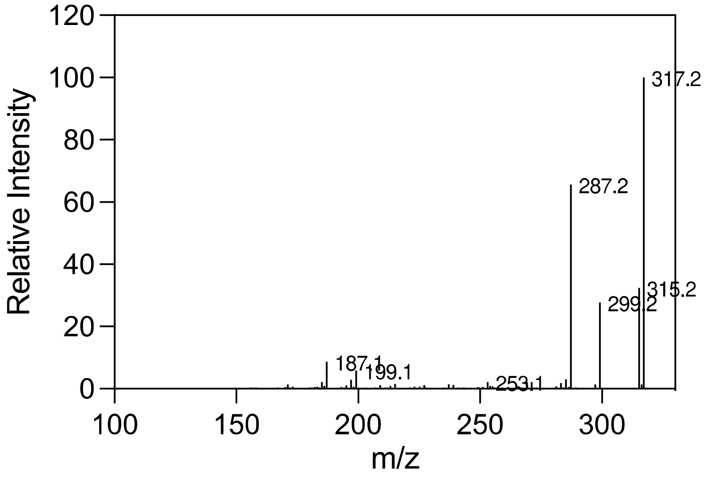

Relative Intensity
m/z
317.2
287.2
315.2
299.2
187.1
199.1
253.1

**31**

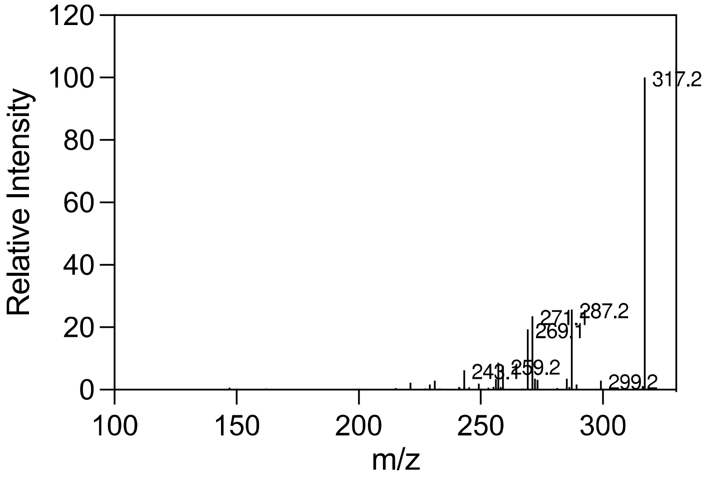
Relative Intensity
m/z
317.2
287.2
271.1
269.1
243.1
259.2
299.2

**32**

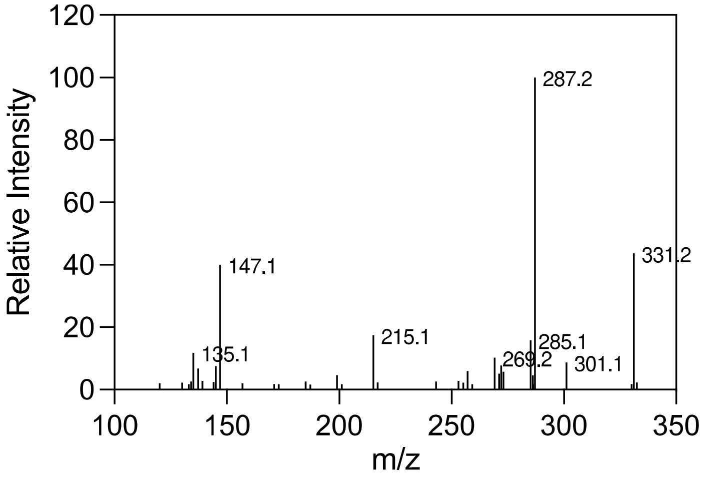
Relative Intensity
m/z
287.2
331.2
147.1
215.1
135.1
285.1
269.2
301.1

**33**

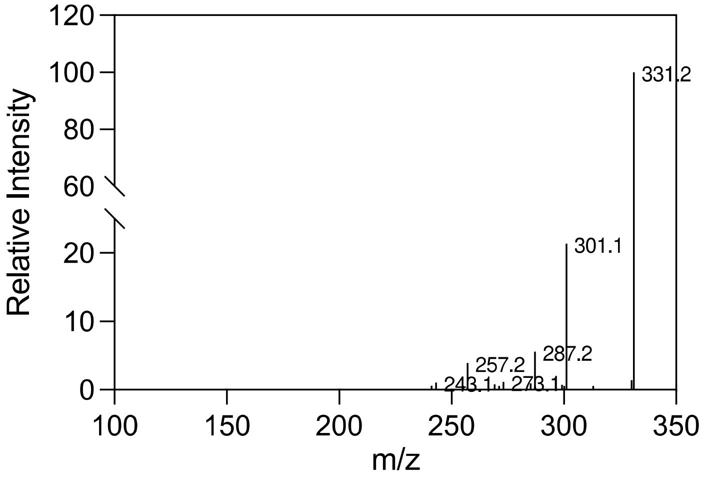
Relative Intensity
m/z
331.2
301.1
287.2
257.2
243.1
273.1

# 34

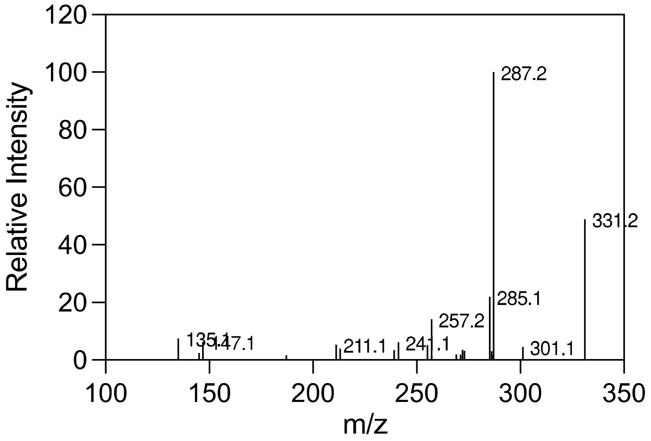
Relative Intensity
m/z
287.2
331.2
285.1
257.2
135.1
147.1
211.1
241.1
301.1

# 35

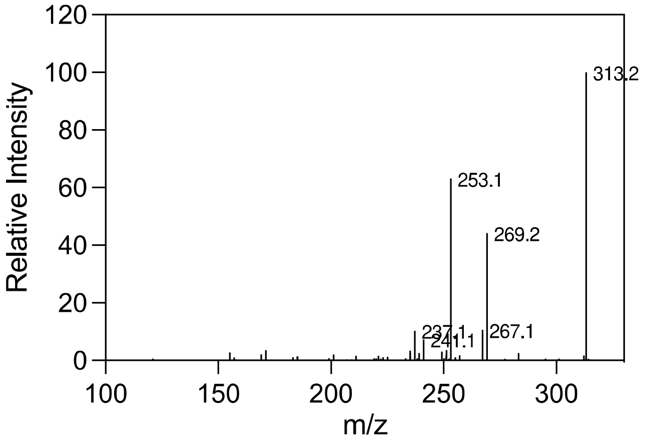
Relative Intensity
m/z
313.2
253.1
269.2
237.1
241.1
267.1

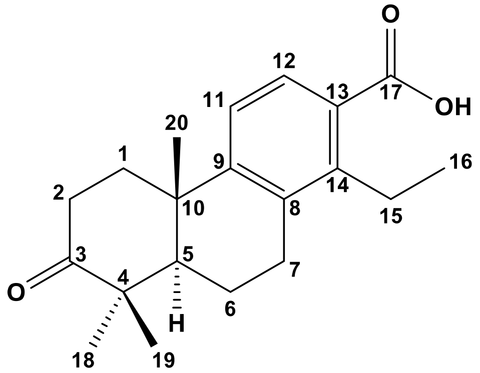
O
OH
H
O

# 36

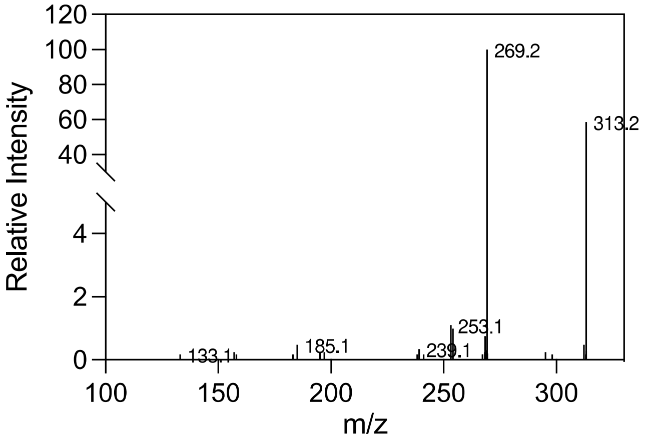
Relative Intensity
m/z
269.2
313.2
253.1
185.1
133.1
239.1

**37**

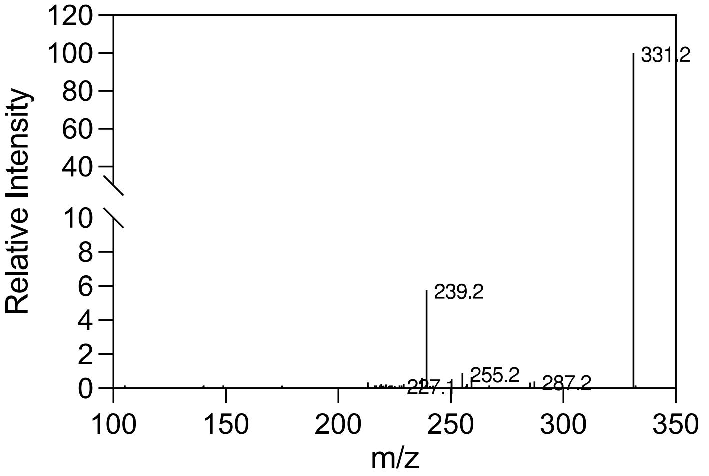

Relative Intensity
m/z
331.2
239.2
227.1
255.2
287.2

**38**

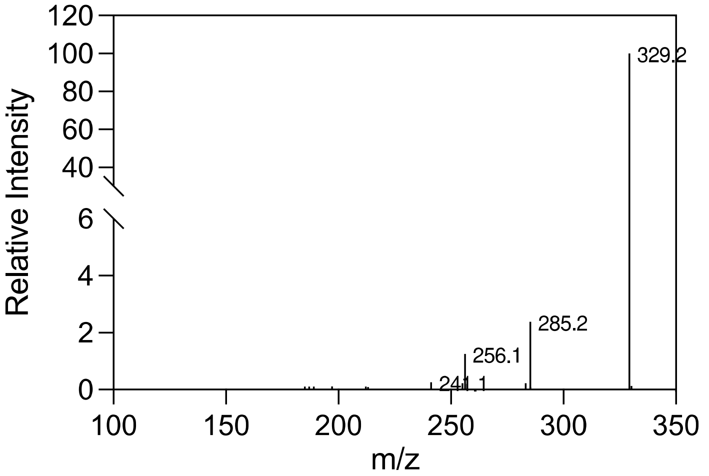

Relative Intensity
m/z
329.2
285.2
256.1
241.1

**39**

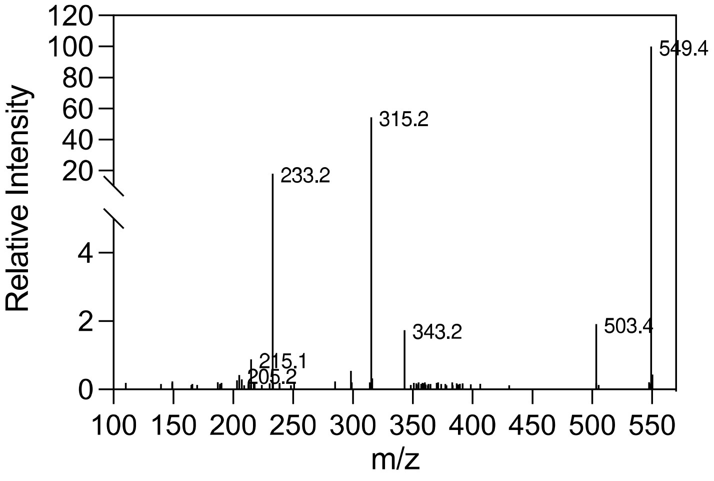

Relative Intensity
m/z
549.4
315.2
233.2
343.2
503.4
215.1
205.2

## 40

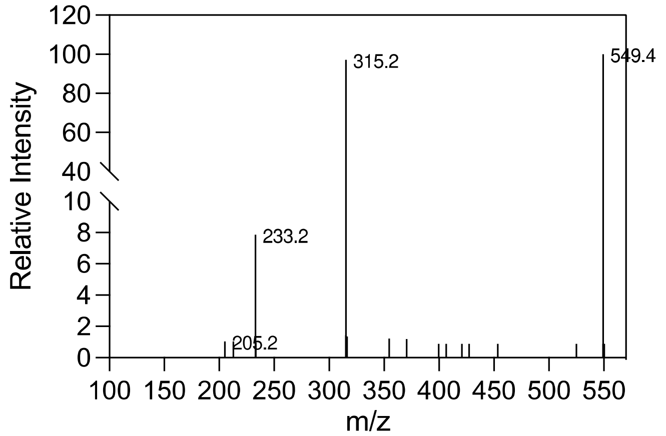

## 41

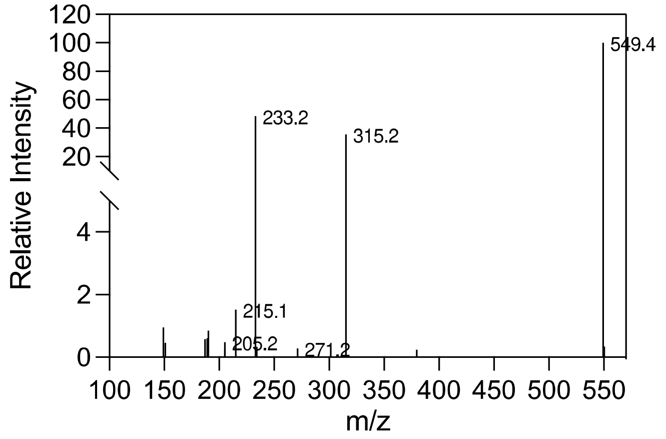

## 42

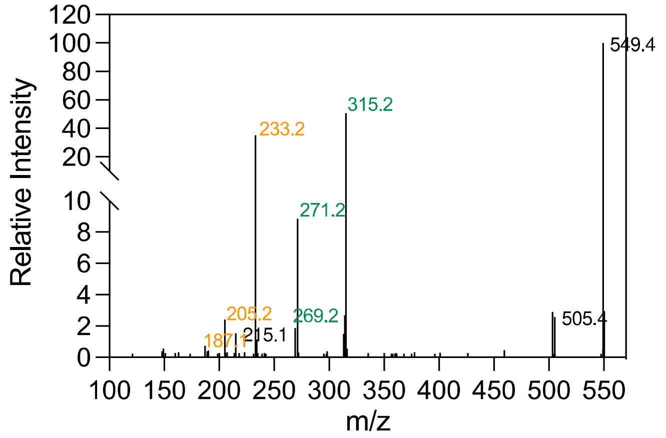

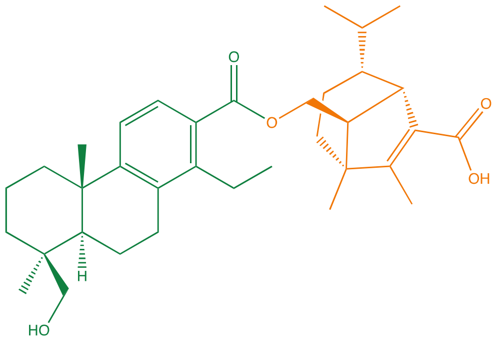

**Putative, NO NMR**

# 43

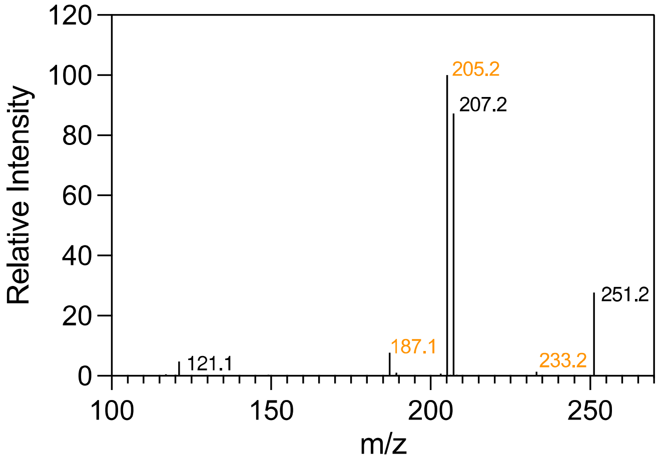

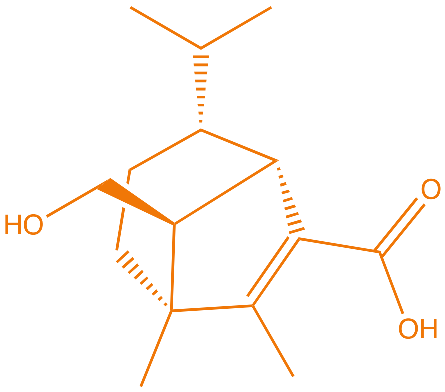

**Putative, NO NMR**

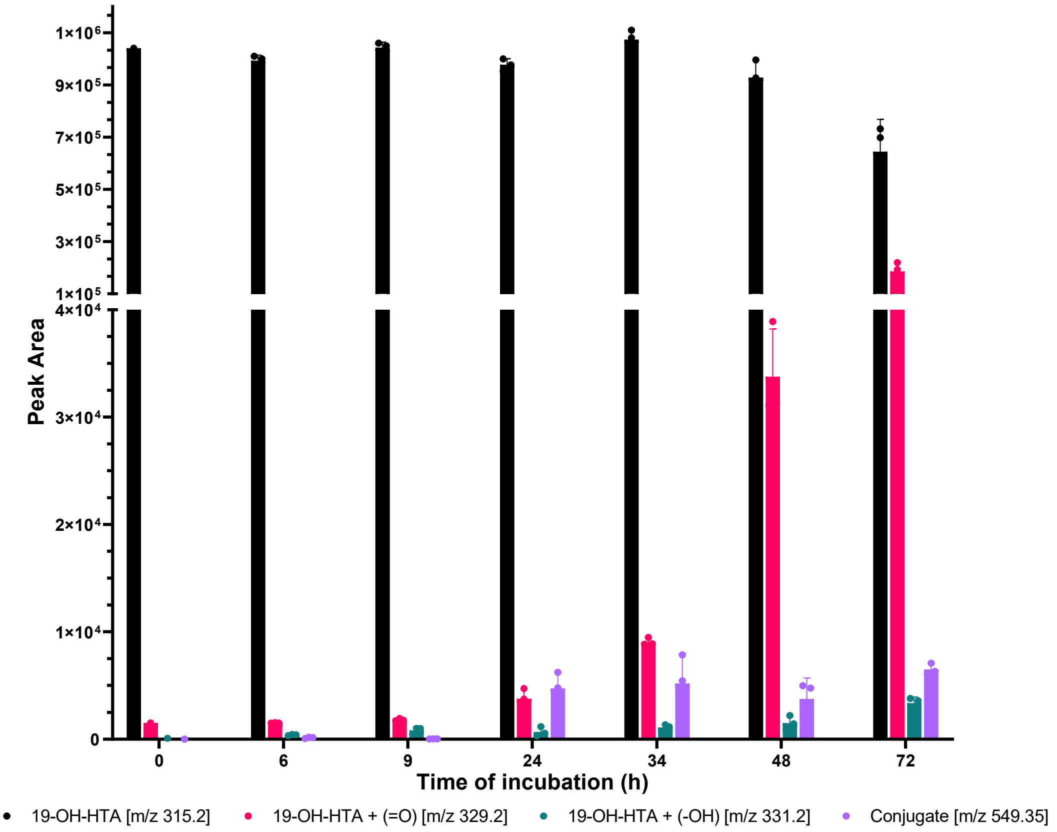

**Fig. S10: Time course of modification of 19-OH-HTA (21a) by *B. sorokiniana*.**

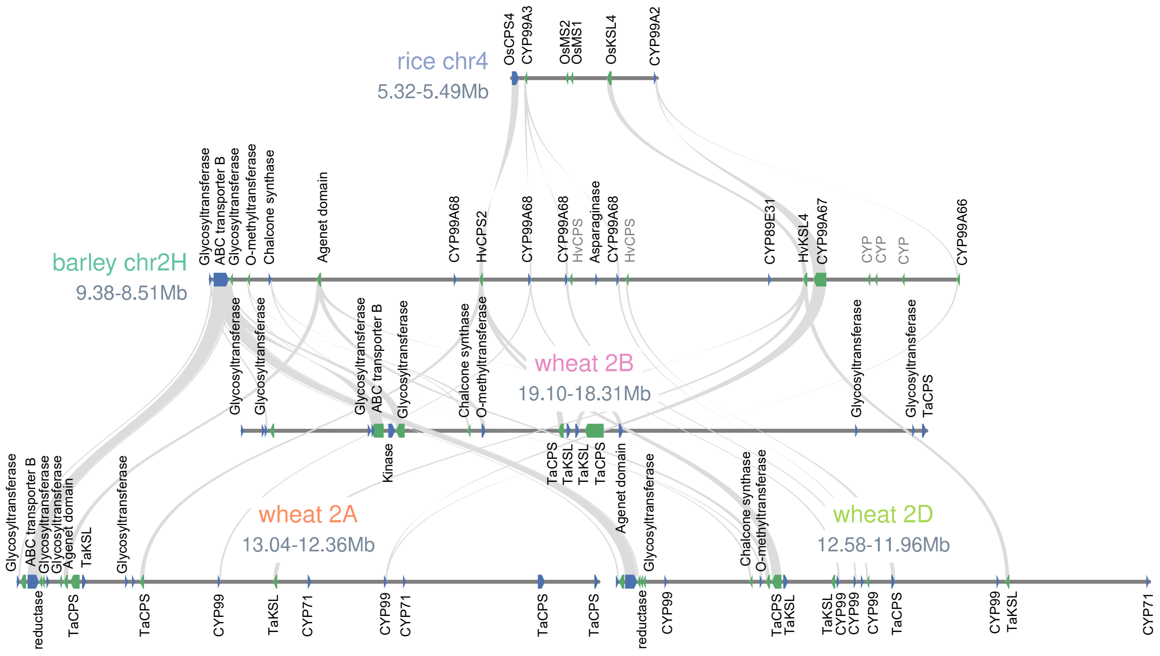

**Fig. S11: Micro-synteny of barley, rice and wheat biosynthetic gene clusters.**

Micro-synteny analysis of genes in momilactone BGC of rice, hordediene BGC of barley, and the diterpenoid BGCs of wheat. The annotation of genes in wheat BGCs derives from the functional annotation of the best hit by a blastp search against the Swiss-Prot database (*19*). BGC, biosynthetic gene cluster.

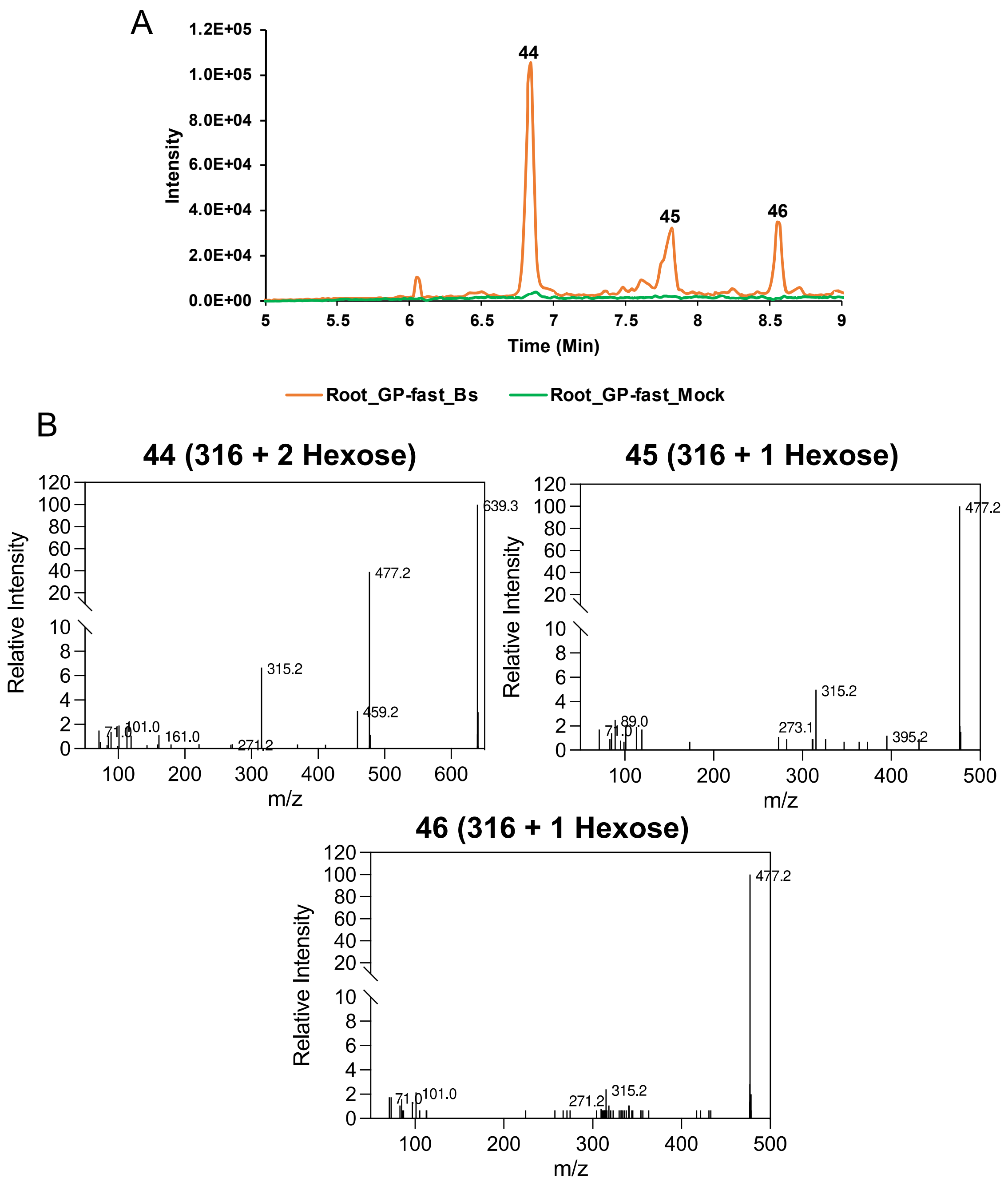

**Fig. S12: Glycosylated hordedanes detected from *Bs*-infected barley roots.**

**A)** LC-HRMS (negative mode) chromatograms of glycosylated diterpenoids from barley roots after *Bs* infection or mock treatment. Selected ions, m/z 477.2, 639.3.

**B)** MS/MS spectra of glycosylated diterpenoids presented in **(A)**.

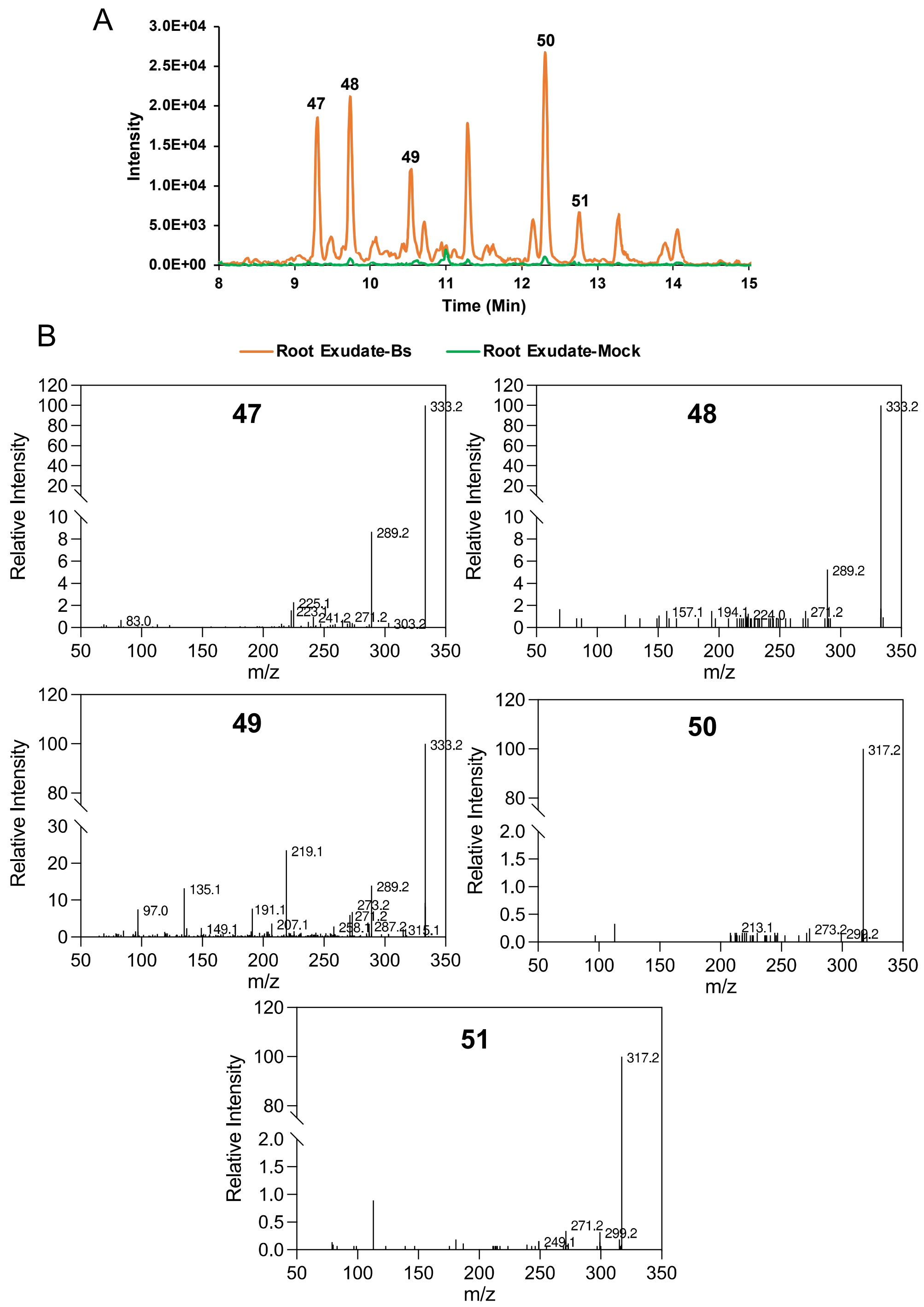

**Fig. S13: Putative diterpenoids detected from wheat root exudates after *Bs* infection.**

**A)** LC-HRMS (negative mode) chromatograms of wheat diterpenoids.Selected ions, m/z 317.2, 333.3. **B)** MS/MS spectra of diterpenoids presented in **(A)**.

| Genome version | | | Gene status | Name |
|---|---|---|---|---|
| MorexV1 (2017) | MorexV2 (2019) | MorexV3 (2021) | | |
| 2Hr1G004480 | 2HG0081780 | 2HG0099280 | FL | CYP99A66 |
| 2Hr1G004510 | 2HG0081840 | 2HG0099340 | FL | |
| 2Hr1G004520 | 2HG0081840 | NP | Pseudo | |
| 2Hr1G004530 | 2HG0081840 | 2HG0099350 | FL | CYP99A67 |
| 2Hr1G004540 | 2HG0081850 | 2HG0099360 | FL | HvKSL4 |
| 2Hr1G004550 | 2HG0081860<br>UnG0627890<br>UnG0631950 | 2HG0099370[a]<br>2HG0099420[a]<br>2HG0099430[a] | FL | CYP89E31 |
| NP | 2HG0081880 | 2HG0099470 | Pseudo | ψCPS |
| NP | 2HG0081920 | 2HG0099500 | Pseudo | ψCPS |
| 2Hr1G004600 | 2HG0081930 | 2HG0099550 | FL | CYP99A68[b] |
| 2Hr1G004610 | 2HG0081980 | 2HG0099550 | FL | CYP99A68[b] |
| 2Hr1G004620 | 2HG0082000 | 2HG0099570 | FL | HvCPS2 |
| 2Hr1G004640<br>2Hr1G004650 | 2HG0081890<br>2HG0082010<br>2HG0081930<br>UnG0636400 | 2HG0099480 | FL | CYP99A68[b] |

**Tab. S1: List of genes from the chromosome 2 diterpenoid phytoalexin cluster.**

Notes: (a) the amino acid sequences of these two genes are identical. The sequences only differ in the 5'-UTR. (b) the amino acid sequences are identical except for one amino acid change in 2Hr1G004600. NP: not present, FL: full length. (ψ) pseudogene.

| Protein name | Species | Protein ID |
|---|---|---|
| AtCPS | *Arabidopsis thaliana* | NP_192187 |
| CmCPS1 | *Cucurbita maxima* | AAD04292 |
| OsCPS1 | *Orzya sativa* | BAF08464 |
| OsCPS2 | *Orzya sativa* | BAH91759 |
| OsCPS4 | *Orzya sativa* | NP_001052171 |
| TaCPS1 | *Triticum aestivum* | BAH56558 |
| TaCPS2 | *Triticum aestivum* | BAH56559 |
| TaCPS3 | *Triticum aestivum* | BAH56560 |
| TaCPS4 | *Triticum aestivum* | BAP01383 |
| HvCPS1 | *Hordeum vulgare* | AAT49065 |
| ZmCPS1 | *Zea mays* | NP_001105329 |
| ZmCPS2 | *Zea mays* | NP_001105257 |
| ZmCPS3 | *Zea mays* | AFW57228 |
| ZmCPS4 | *Zea mays* | AFW60403 |
| HvCPS2 | *Hordeum vulgare* | BAJ95441 |

**Tab. S2: CPS sequences used for phylogenetic analysis in Fig. S1.**

| **Protein name** | **Species** | **Protein ID** |
|---|---|---|
| AtKS | *Arabidopsis thaliana* | AAC39443 |
| CmKS | *Cucurbita maxima* | Q39548 |
| OsKS | *Orzya sativa* | NP_001053841 |
| OsKSL4 | *Orzya sativa* | NP_001052175 |
| OsKSL5 | *Orzya sativa* | NP_001047190 |
| OsKSL6 | *Orzya sativa* | ABH10733 |
| OsKSL7 | *Orzya sativa* | NP_001047186 |
| OsKSL8 | *Orzya sativa* | NP_001067887 |
| OsKSL10 | *Orzya sativa* | NP_001066799 |
| OsKSL11 | *Orzya sativa* | Q1AHB2 |
| TaKS | *Triticum aestivum* | BAL41693 |
| TaKSL1 | *Triticum aestivum* | BAL41688 |
| TaKSL2 | *Triticum aestivum* | BAL41689 |
| TaKSL3 | *Triticum aestivum* | BAL41690 |
| TaKSL4 | *Triticum aestivum* | BAL41691 |
| HvKS | *Hordeum vulgare* | AAT49066 |
| ZmKSL1 | *Zea mays* | AFW61735 |
| ZmKSL2 | *Zea mays* | DAA54948 |
| ZmKSL3 | *Zea mays* | DAA36069 |
| ZmKSL4 | *Zea mays* | DAA49845 |
| HvKSL4 | *Hordeum vulgare* | BAK01991 |

**Tab. S3: KS and KSL sequences used for phylogenetic analysis in Fig. S1.**

| **Accession no.** | **Name** | **Organism** | **Function** |
|---|---|---|---|
| AB014459 | AtCYP51A2 | *Arabidopsis thaliana* | |
| Q93Z79 | AtCYP714A1 | *Arabidopsis thaliana* | |
| Q6NKZ8 | AtCYP714A2 | *Arabidopsis thaliana* | |
| AF318500 | AtKAO1 | *Arabidopsis thaliana* | Kaurenoic acid oxidase (KAO) |
| AF318501 | AtKAO2 | *Arabidopsis thaliana* | Kaurenoic acid oxidase (KAO) |
| AF047719 | AtKO | *Arabidopsis thaliana* | Kaurene oxidase (KO) |
| Q42602.2 | AtCYP89A2 | *Arabidopsis thaliana* | unknown function |
| HQ658173 | CamKAO | *Castanea mollissima* | |
| ACQ99375 | CaKO | *Coffea arabica* | Kaurene oxidase (KO) |
| KT382342 | CfCYP71D381 | *Coleus forskohlii* | Forskolin biosynthesis |
| KT382346 | CfCYP76AH10 | *Coleus forskohlii* | Forskolin biosynthesis |
| KT382349 | CfCYP76AH11 | *Coleus forskohlii* | Forskolin biosynthesis |
| KT382358 | CfCYP76AH15 | *Coleus forskohlii* | Forskolin biosynthesis |
| KT382359 | CfCYP76AH16 | *Coleus forskohlii* | Forskolin biosynthesis |
| KT382360 | CfCYP76AH17 | *Coleus forskohlii* | Forskolin biosynthesis |
| KT382348 | CfCYP76AH8 | *Coleus forskohlii* | Forskolin biosynthesis |
| KT382347 | CfCYP76AH9 | *Coleus forskohlii* | Forskolin biosynthesis |
| NP_001267703 | CsKO | *Cucumis sativus* | Kaurene oxidase (KO) |
| AF212991 | CumKAO | *Cucurbita maxima* | Kaurenoic acid oxidase (KAO) |
| AF212990 | CumKO | *Cucurbita maxima* | Kaurene oxidase (KO) |
| KR350668 | ElCYP71D445 | *Euphorbia lathyris* | Casbene oxidase |
| KR350669 | ElCYP726A27 | *Euphorbia lathyris* | Casbene oxidase |
| KX428471 | EpCYP71D365 | *Euphorbia peplus* | Casbene oxidase |
| KJ026362 | EpCYP726A19 | *Euphorbia peplus* | Casbene synthase |
| KF986823.1 | EpCYP726A4 | *Euphorbia peplus* | Casbene oxidase |
| KF773141 | GbCYP716B | *Ginkgo biloba* | taxoid-9α-hydroxylase |
| KHN31869 | GsKO | *Glycine soja* | Kaurene oxidase (KO) |
| FR666915 | HaKAO1 | *Helianthus annuus* | Kaurenoic acid oxidase (KAO) |
| FR666916 | HaKAO2 | *Helianthus annuus* | Kaurenoic acid oxidase (KAO) |
| AF326277 | HvKAO1 | *Hordeum vulgare* | Kaurenoic acid oxidase (KAO) |
| A0A287GY22 | HvCYP89E31 | *Hordeum vulgare* | Hordediene oxidase |
| A0A287GY21 | HvCYP99A66 | *Hordeum vulgare* | Hordediene oxidase |
| A0A287GY30 | HvCYP99A67 | *Hordeum vulgare* | Hordediene oxidase |
| AK249794 | HvCYP99A68 | *Hordeum vulgare* | Hordetriene oxidase |
| KX060559 | JcCYP71D495 | *Jatropha curcas* | Casbene oxidase |
| KF986815 | JcCYP726A20 | *Jatropha curcas* | Casbene oxidase |
| KF986816 | JcCYP726A21 | *Jatropha curcas* | |
| KF986818 | JcCYP726A23 | *Jatropha curcas* | |

| | | | |
|---|---|---|---|
| KF986819 | JcCYP726A24 | *Jatropha curcas* | |
| KX060558 | JcCYP726A35 | *Jatropha curcas* | Casbene oxidase |
| JF929910 | JcKO | *Jatropha curcas* | Kaurene oxidase (KO) |
| AB370238 | LsKAO | *Lactuca sativa* | Kaurenoic acid oxidase (KAO) |
| KF437682 | MdKAO | *Malus domestica* | Kaurenoic acid oxidase (KAO) |
| AY563549 | MdKO | *Malus domestica* | Kaurene oxidase (KO) |
| XM_013607618 | MtKAO | *Medicago truncatula* | Kaurenoic acid oxidase (KAO) |
| ADE06669 | McKO | *Momordica charantia* | Kaurene oxidase (KO) |
| XP_010089925 | MnKO | *Morus notabilis* | Kaurene oxidase (KO) |
| AF116915 | NtCYP51 | *Nicotiana tabacum* | obtusifoliol 14-alpha demethylase |
| AF166332 | NtCYP71D16 | *Nicotiana tabacum* | CBT-ol oxidase |
| Q7XHW5 | OsCYP714B1 | *Oryza sativa* | |
| Q0DS59 | OsCYP714B2 | *Oryza sativa* | |
| AK109526 | OsCYP714D1 | *Oryza sativa* | |
| AK107418 | OsCYP71Z6 | *Oryza sativa* | |
| AK070167 | OsCYP71Z7 | *Oryza sativa* | Cassadiene oxidase |
| AK059010 | OsCYP76M5 | *Oryza sativa* | |
| AK101003 | OsCYP76M6 | *Oryza sativa* | Oryzalexin synthase |
| AK105913 | OsCYP76M7 | *Oryza sativa* | |
| AK069701 | OsCYP76M8 | *Oryza sativa* | Oryzalexin synthase |
| AK071864 | OsCYP99A2 | *Oryza sativa* | Pimaradiene oxidase |
| AK071546 | OsCYP99A3 | *Oryza sativa* | Pimaradiene oxidase |
| Q5VRM7 | OsKAO | *Oryza sativa* | Kaurenoic acid oxidase (KAO) |
| Q5Z5R4 | OsKO2 | *Oryza sativa* | Kaurene oxidase (KO) |
| AY579214 | OsKOL4 | *Oryza sativa* | Kaurene oxidase (KO) |
| AY660664 | OsKOL5 | *Oryza sativa* | Kaurene oxidase (KO) |
| BAK19917 | PhpKO | *Physcomitrella patens* | Kaurene oxidase (KO) |
| HM245408 | PsiCYP720B10 | *Picea sitchensis* | |
| HM245397 | PsiCYP720B12 | *Picea sitchensis* | Hydroxyl abietene oxidase |
| HM245398 | PsiCYP720B15 | *Picea sitchensis* | |
| HM245399 | PsiCYP720B16 | *Picea sitchensis* | |
| HM245402 | PsiCYP720B2 | *Picea sitchensis* | Hydroxyl abietene oxidase |
| HM245403 | PsiCYP720B4 | *Picea sitchensis* | Diterpene C-18 oxidase |
| HM245406 | PsiCYP720B7 | *Picea sitchensis* | |
| HM245407 | PsiCYP720B8 | *Picea sitchensis* | |
| HM245410 | PsiCYP720B9 | *Picea sitchensis* | |
| KJ845667 | PbCYP720B2 | *Pinus banksiana* | Hydroxyl abietene oxidase |
| KJ845671 | PcCYP720B1 | *Pinus contorta* | Abietadienol/abietadienal oxidase |
| KJ845675 | PcCYP720B11 | *Pinus contorta* | |
| KJ845676 | PcCYP720B12 | *Pinus contorta* | Hydroxyl abietene oxidase |
| KJ845672 | PcCYP720B2 | *Pinus contorta* | Hydroxyl abietene oxidase |
| AY779541 | PtaCYP720B1 | *Pinus taeda* | Abietadienol/abietadienal oxidase |
| AF537321 | PsaKAO1 | *Pisum sativum* | Kaurenoic acid oxidase (KAO) |

| | | | |
|---|---|---|---|
| AF537322 | PsaKAO2 | *Pisum sativum* | Kaurenoic acid oxidase (KAO) |
| AAP69988 | PsaKO | *Pisum sativum* | Kaurene oxidase (KO) |
| XP_006386514 | PtrKO | *Populus trichocarpa* | Kaurene oxidase (KO) |
| AEK01241 | PcKO | *Pyrus communis* | Kaurene oxidase (KO) |
| HM003112 | PypKO | *Pyrus pyrifolia* | Kaurene oxidase (KO) |
| KF986809 | RcCYP726A13 | *Ricinus communis* | |
| KF986810 | RcCYP726A14 | *Ricinus communis* | Casbene oxidase |
| KF986811 | RcCYP726A15 | *Ricinus communis* | Neocembrene oxidase |
| KF986812 | RcCYP726A16 | *Ricinus communis* | Casbene oxidase |
| KF986813 | RcCYP726A17 | *Ricinus communis* | Casbene oxidase |
| KF986814 | RcCYP726A18 | *Ricinus communis* | Casbene oxidase |
| KP091843 | RoCYP76AH22 | *Rosmariuns officinalis* | Hydroxyferruginol synthase |
| KP091844 | RoCYP76AH23 | *Rosmariuns officinalis* | Hydroxyferruginol synthase |
| | RoCYP76AH4 | *Rosmariuns officinalis* | Hydroxyferruginol synthase |
| KX431219 | RoCYP76AK7 | *Rosmariuns officinalis* | C20 oxidase |
| KX431220 | RoCYP76AK8 | *Rosmariuns officinalis* | C20 oxidase |
| KP091842 | SfCYP76AH24 | *Salvia fruticosa* | Hydroxyferruginol synthase |
| KX431218 | SfCYP76AK6 | *Salvia fruticosa* | C20 oxidase |
| JX422213 | SmCYP76AH1 | *Salvia miltiorrhiza* | Ferruginol synthase |
| KR140168 | SmCYP76AH3 | *Salvia miltiorrhiza* | Hydroxyferruginol synthase |
| KR140169 | SmCYP76AK1 | *Salvia miltiorrhiza* | C20 oxidase |
| KP337688 | SmCYP76AK2 | *Salvia miltiorrhiza* | |
| KP337689 | SmCYP76AK3 | *Salvia miltiorrhiza* | |
| KJ606394 | SmKO | *Salvia miltiorrhiza* | Kaurene oxidase (KO) |
| KT157042 | SpCYP71BE52 | *Salvia pomifera* | C2 oxidase |
| KT157044 | SpCYP76AH24 | *Salvia pomifera* | Hydroxyferruginol synthase |
| KT157045 | SpCYP76AK6 | *Salvia pomifera* | C20 oxidase |
| XP_034572432.1 | SvCYP89A2-like | *Setaria viridis* | unknown function |
| Solyc08g005650.2.1 | SlCYP71BN1 | *Solanum lycopersisum* | |
| U74319 | SbCYP51 | *Sorghum bicolor* | obtusifoliol 14-alpha demethylase |
| AY364317 | SrKO1 | *Stevia rebaudiana* | Kaurene oxidase (KO) |
| AY995178 | SrKO2 | *Stevia rebaudiana* | Kaurene oxidase (KO) |
| AY518383 | TcaCYP725A6, T2OH | *Taxus canadensis* | taxoid-2α-hydroxylase |
| AF318211 | TcuCYP725A1, T10OH | *Taxus cuspidata* | taxoid-10β-hydroxylase |
| AY056019 | TcuCYP725A2, T13OH | *Taxus cuspidata* | taxoid-13α-hydroxylase |
| AY188177 | TcuCYP725A3, T14OH | *Taxus cuspidata* | taxoid-14β-hydroxylase |
| AY289209 | TcuCYP725A4, T5OH | *Taxus cuspidata* | |
| AY307951 | TcuCYP725A5, T7OH | *Taxus cuspidata* | taxoid-7β-hydroxylase |
| ADZ55286 | TaKO | *Triticum aestivum* | Kaurene oxidase (KO) |

| | | | |
|---|---|---|---|
| KAF6985951 | TaCYP89A2-like | *Triticum aestivum* | unknown function |
| MG696754.1 | VacCYP76BK1 | *Vitex agnus-castus* | |
| JQ086553 | VvKO | *Vitis vinifera* | Kaurene oxidase (KO) |
| ACG38493 | ZmKO | *Zea mays* | Kaurene oxidase (KO) |

**Tab. S4: List of CYP sequences used for phylogenetic analysis in Fig. S2.**

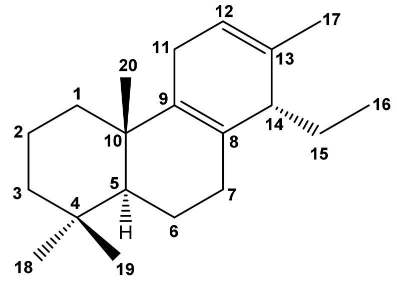

| Pos. | $^{13}$C: δ [ppm] | $^{1}$H: δ [ppm] m (J[Hz]) | selected HMBC (H → C) | ROE[c] |
|---|---|---|---|---|
| 1 | 37.38 | 1.66[a] β;<br>1.043 ddd (13.0/13.0/3.7) α | | 20; |
| 2 | 19.37 | 1.565[b] α;<br>1.40[a] β | | |
| 3 | 42.01 | 1.39[a] β;<br>1.148 ddd (13.7/13.2/4.1) α | | 18[d] |
| 4 | 33.39 | --- | | |
| 5 | 52.49 | 1.210 dd (12.5/2.0) α | | 1α[d],7α[d],18[d] |
| 6 | 19.20 | 1.687[b] α;<br>1.503 m β | 10;<br>5,7,10 | 18; |
| 7 | 30.81 | 2.189 m α;<br>1.750 br dd (17.1/6.2) β | | 5[d],15[d],16[d],18[d];<br>14[d] |
| 8 | 126.41 | --- | | |
| 9 | 138.22 | --- | | |
| 10 | 37.86 | --- | | |
| 11 | 26.41 | 2.53[a] | | 20 |
| 12 | 121.84 | 5.611 m | | 11[d],17[d] |
| 13 | 133.66 | --- | | |
| 14 | 46.64 | 2.462 m | 8,9,12,13,15 | 7α[d],7β[d],15[d] |
| 15 | 21.41 | 1.61[a] | | |
| 16 | 22.89 | 0.763 t (7.4) | 14,15 | |
| 17 | 7.72 | 1.656 qd (1.7/0.5) | 12,13,14 | |
| 18 | 33.44 | 0.921 s | 3,4,5,19 | |
| 19 | 21.94 | 0.877 s | 3,4,5,18 | 20 |
| 20 | 19.42 | 1.014 s | 1,5,9,10 | |

[a] $^{1}$H chemical shift of HSQC correlation peaks; [b] $^{1}$H chemical shift from selective TOCSY spectrum; [c] each NOE is listed only once; [d] correlations (H → H) from selective ROESY spectra. 14*S* configuration is more likely because of NOE H-7β/H-14 und H-7α/H-15.

**Tab. S5: NMR data of compound 1 ($C_6D_6$)**

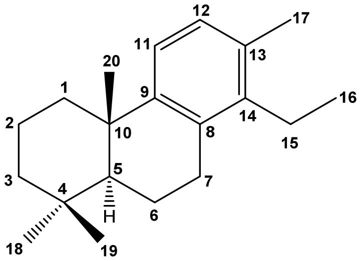

| Pos. | $^{13}$C: δ [ppm] | $^{1}$H: δ [ppm] m (J[Hz][a]) | HMBC (H→C) | ROE[d] |
|---|---|---|---|---|
| 1 | 39.6 | 2.206 m β;<br>1.344 td-like (13.2/3.8) α | | 11,20; |
| 2 | 19.8 | 1.670 qt-like (13.9/3.4) β;<br>1.5054 m α | 1<br>4,10 | 20; |
| 3 | 41.9 | 1.409 m β;<br>1.138 td-like (13.5/4.0) α | <br>4 | |
| 4 | 33.4 | --- | | |
| 5 | 50.0 | 1.243 dd (12.6/2.2) α | | 18 |
| 6 | 19.7 | 1.797 ddt-like (13.2/7,8/1.9) α;<br>1.613 m β | | |
| 7 | 28.0 | 2.849 ddd (17.1/6.6/1.3) β;<br>2.643 ddd (17.1/11.6/7.8) α | | |
| 8 | 132.5[b] | --- | | |
| 9 | 148.5 | --- | | |
| 10 | 38.1 | --- | | |
| 11 | 122.4 | 7.092 d (8.0) | | |
| 12 | 128.4[c] | 7.020 d (8.0) | | 17 |
| 13 | 132.6[b] | --- | | |
| 14 | 140.2 | --- | | |
| 15 | 22.4 | 2.52 m | 16 | |
| 16 | 13.4 | 1.035 t (7.5) | 14,15 | |
| 17 | 19.5 | 2.209 s | 12,13,14 | |
| 18 | 33.4 | 0.923 s | 3,4,5,19 | |
| 19 | 21.8 | 0.899 s | 3,4,5,18 | 20 |
| 20 | 25.2 | 1.197 br s | 1,5,9,10 | |

[a] "-like" multiplicities: the given value represents the distance of the multiplet lines, not the exact coupling constant; [b] might be interchanged; [c] $^{13}$C chemical shift of HSQC correlation peak; [d] each NOE is listed only once.

**Tab. S6: NMR data of compound 5 ($C_6D_6$).**

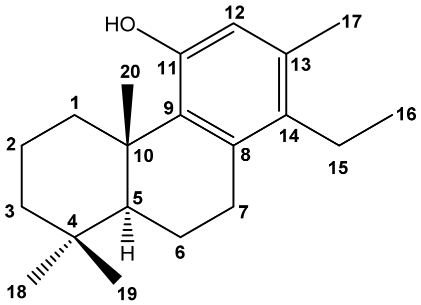

| Pos. | $^{13}$C: δ [ppm] | $^{1}$H: δ [ppm] m (J[Hz][a]) | HMBC (H→C) | ROE (H ↔ H)[b] |
|---|---|---|---|---|
| 1 | 37.26 | 3.848 br dt (13.4/3.7) β;<br>1.350 td (13.4/3.7) α | 2,3,5,10;<br>2,10 | 20; |
| 2 | 19.98 | 1.824 dtt (14.0/13.4/3.7) β;<br>1.582 dt-like (14.0/3.7) α | 1,3,4,10;<br>4,10 | 20; |
| 3 | 41.83 | 1.453 dtd-like (13.1/3.5/1.5) β;<br>1.234 td-like (13.4/4.2) α | 2,4;<br>2,4,18,19 | |
| 4 | 33.77 | --- | | |
| 5 | 52.88 | 1.276 dd (12.2/1.3) α | 4,6,10,18,20 | 18 |
| 6 | 19.64 | 1.769 br dd-like (12.9/6.8) α;<br>1.521 qt-like (12.5/5.4) β | 8,10;<br>5,10 | 18,19;<br>19 |
| 7 | 30.64 | 2.791 ddd (16.8/5.4/1.5) β;<br>2.614 ddd (16.8/12.6/6.8) α | 5,6,8,9,14;<br>6,8 | 15; |
| 8 | 136.02 | --- | | |
| 9 | 133.62 | --- | | |
| 10 | 39.72 | --- | | |
| 11 | 152.11 | --- | | |
| 12 | 116.67 | 5.668 br s | 9,11,14,17 | 11-OH,17 |
| 13 | 133.29 | --- | | |
| 14 | 132.84 | --- | | |
| 15 | 21.99 | 2.475 q (7.5) | 8,13,14,16 | 17 |
| 16 | 13.74 | 1.030 t (7.5) | 14,15 | |
| 17 | 19.12 | 2.097 br s | 12,13,14 | |
| 18 | 33.88 | 0.960 s | 3,4,5,19 | |
| 19 | 22.36 | 0.954 s | 3,4,5,18 | 20 |
| 20 | 20.05 | 1.549 s | 1,5,9,10 | |
| 11-OH | --- | 3.726 s | 9,11,12 | |

[a] "-like" multiplicities: the given value represents the distance of the multiplet lines, not the exact coupling constant; [b] each NOE is listed only once.

**Tab. S7: NMR data of compound 8 ($C_6D_6$).**

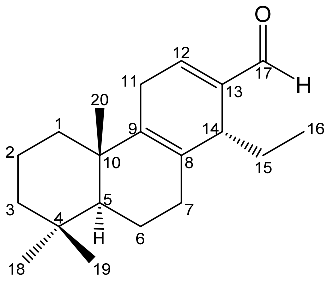

| Pos. | $^{13}C$: δ [ppm] | $^{1}H$: δ [ppm] m (J[Hz][a]) | HMBC (H→C) | COSY | ROE[d] |
|---|---|---|---|---|---|
| 1 | 37.33 | 0.927 m α<br>1.514 m β | 3, 10, 2<br>2, 10 | 2 | ;<br>20, 11 |
| 2 | 19.26 | 1.400 m α<br>1.520 m β | 4, 1, 10,3<br>10, 1 | | ;<br>20 |
| 3 | 41.84 | 1.102[c] dt-like (4.3, 13.5) α<br>1.380 m[c] β | 2,4,19<br>5, 19, 2 | | |
| 4 | 33.30 | --- | | | |
| 5 | 52.26 | 1.076 dd (2.1,12.6) α | 10, 19, 7, 20 | 6 | 18 |
| 6 | 18.88 | 1.329[c] qd-like (6.4, 12.4) β<br>1.552[c] br dd-like (7.3, 12.6) α | 7, 10, 5<br>7, 10, 5, 8 | 7,5 | <br>18 |
| 7 | 30.41 | 1.636[c] br dd like (6.4, 17.5) β<br>2.066 m α | 6, 8, 9, 5<br>6, 8, 9 | 6 | |
| 8 | 127.65 | --- | | | |
| 9 | 136.16 | --- | | | |
| 10 | 37.70 | --- | | | |
| 11 | 27.25 | 2.432 m | 8,9,13, 12 | 12 | 20 |
| 12 | 148.83 | 6.214 dd (4.4, 3.3) | 11, 14, 17, 9 | 11 | 17 |
| 13 | 142.69 | --- | | | |
| 14 | 39.17 | 3.272 m | 7, 8, 9, 12, 13, 15, 16, 17 | 15 | 7β |
| 15 | 24.47 | 1.872 m [c]<br>1.613 m [c] | 16, 14, 8, 13;<br>16, 14, 8, 13 | 14,16 | |
| 16 | 8.27 | 0.609 t (7.5) | 14, 15 | 15 | 7β, 5 |
| 17 | 191.91 | 9.350 s | 12, 13, 8, 14 | | |
| 18 | 33.34 | 0.869 s | 3,4,5,19 | | 6α |
| 19 | 21.82 | 0.836 s | 3,4,5,18 | | 6α, 20 |
| 20 | 19.14 | 0.891 s | 1, 9, 5, 10 | | |

[a] "-like" multiplicities: the given value represents the distance of the multiplet lines, not the exact coupling constant; [c] $^{1}H$ chemical shift of HSQC correlation peak; [d] each NOE is listed only once. 14R configuration is more likely because of NOE H-7β/H-14

**Tab. S8: NMR data of compound 6 ($C_6D_6$).**

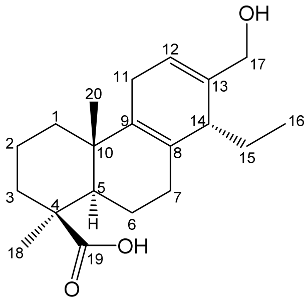

| Pos. | $^{13}$C: δ [ppm] | $^{1}$H: δ [ppm] m (J[Hz]$^{a}$) | HMBC (H→C) | COSY | ROE$^{d}$ |
|---|---|---|---|---|---|
| 1 | 37.61 | 1.656$^{c}$ m β<br>0.974$^{c}$ m α | 2, 3, 5,10, 20<br>2, 3, 5, 9, 10, 20 | 2 | 11; |
| 2 | 19.90 | 1.451$^{c}$ m α<br>2.079$^{c}$ m β | 1, 3, 4, 10, | 3 | ;<br>20 |
| 3 | 37.65 | 2.308 dt-like (13.4, 3.2) β<br>0.860 td-like(13.7, 4.4) α | 1, 2, 4, 5;<br>1, 2, 4, 5, 18, 19 | 2 | |
| 4 | 43.87 | --- | | | |
| 5 | 53.92 | 1.287$^{c}$ dd (3.3, 10.7) α | 3, 4, 6, 7, 9, 10, 18, 19, 20 | 6 | 1α, 3α |
| 6 | 20.92 | 2.030$^{c}$ m | 8, 5, 7, 10 | 5, 7 | 20 |
| 7 | 31.62 | 2.104$^{c}$ m α<br>1.760 brdd-like (16.0, 2.8) β | ;<br>6, 8, 9 | 6 | |
| 8 | 127.56 | --- | | | |
| 9 | 136.92 | --- | | | |
| 10 | 38.62 | --- | | | |
| 11 | 26.12 | 2.435 br-s | --- | 12 | 20 |
| 12 | 122.16 | 5.711 t-like (3.4) | 9, 11, 14, 17 | 11 | 17, 11 |
| 13 | 137.92 | --- | | | |
| 14 | 42.64 | 2.704 m | 7, 8, 9, 12, 13, 15, 16 | 15 | 7β |
| 15 | 23.42 | 1.564 m | 8, 13, 14, 16 | | 7α |
| 16 | 7.86 | 0.709 t (7.4) | 14, 15 | 15 | |
| 17 | 64.91 | 3.871$^{c}$ m | 12, 13, 14 | | |
| 18 | 28.49 | 1.150 s | 5, 4, 3, 4, 5, 19 | | 3, 6 |
| 19 | 182.42 | --- | | | |
| 20 | 17.38 | 1.043 s | 9, 10, 5, 1 | | 1β |

[a] "-like" multiplicities: the given value represents the distance of the multiplet lines, not the exact coupling constant; [c] $^{1}$H chemical shift of HSQC correlation peak; [d] each NOE is listed only once. 14R configuration is more likely because of NOE H-7β/H-14

**Tab. S9: NMR data of compound 19 ($C_6D_6$).**

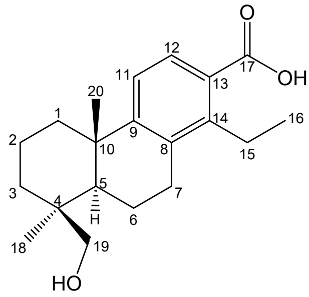

| Pos. | $^{13}$C: δ [ppm] | $^{1}$H: δ [ppm] m (J[Hz]) | selected HMBC (H → C) | ROE[c] | COSY |
|---|---|---|---|---|---|
| 1 | 39.20 | 1.990 dt-like (8.4, 3.5) β<br>1.153 m α | 6, 20, 3, 10 | 2, 11 | 2 |
| 2 | 19.29 | 1.40[e] m<br>1.48[e] m | | | 3 |
| 3 | 35.30 | 1.833[a] β<br>0.814[a] α | 2, 4, 19 | | |
| 4 | 38.84 | --- | | | |
| 5 | 50.08 | 1.163[a] α | 10, 19, 7, 20, 6 | 6 | |
| 6 | 19.28 | 1.761 α<br>1.427[a] β | 6, 5, 10, 7 | 7α;<br>7β | 5 |
| 7 | 28.18 | 2.715 dd (17.2, 6.2) β<br>2.429 ddd α | 9, 14, 8, 5, 6 | 16, 6, 15;<br>6α, 15 | 6 |
| 8 | 134.24 | --- | | | |
| 9 | 154.97 | --- | | | |
| 10 | 38.63 | --- | | | |
| 11 | 122.73 | 6.998 d(8.5) | 13, 8, 10 | 1β | 12 |
| 12 | 129.19 | 7.914 d(8.4) | 17, 9, 14 | | 11 |
| 13 | 126.32 | --- | | | |
| 14 | 145.28 | --- | | | |
| 15 | 23.05 | 3.147<br>2.946 | 16, 13, 8, 14 | 7β<br>7α | 16 |
| 16 | 14.72 | 1.285 t (7.4) | 14, 15 | | 15 |
| 17 | 170.34 | --- | | | |
| 19 | 64.83 | 3.506 d (10.6)<br>3.226 dd (10.6, 1.2) | 18, 3, 10, 5 | 20<br>6β | 20 |
| 18 | 26.96 | 0.950 s | 4, 3, 5, 19 | | 6α |
| 20 | 25.54 | 0.987 s | 9, 5, 10, 1 | 19 | |

[a] $^{1}$H chemical shift of HSQC correlation peaks; [b] $^{1}$H chemical shift from selective TOCSY spectrum; [c] each NOE is listed only once; [e] 1H chemical shift from TOCSY

**Tab. S10: NMR data of compound 21a ($C_6D_6$).**

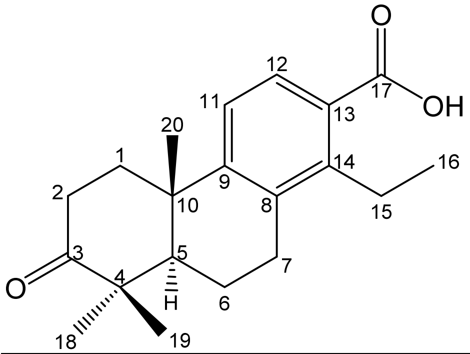

| Pos. | $^{13}$C: δ [ppm] | $^{1}$H: δ [ppm] m ($J$[Hz] | Imp. HMBC (H→C) | Imp. ROE[b] |
|---|---|---|---|---|
| 1 | 37.4 | 1.87[a] m β<br>1.40 m α | 2, 3, 5, 10, 20<br>2, 3, 5, 10, 20 | |
| 2 | 34.5 | 2.34 m β<br>2.28 m α | 1, 3, 4, 10<br>1, 3, 4, 10 | |
| 3 | 214.1 | --- | | |
| 4 | 46.9 | --- | | |
| 5 | 49.4 | 1.44 m | 3, 4, 6, 7, 10 | 1α |
| 6 | 20.2 | 1.39 m β<br>1.33 m α | 5, 7, 8, 10<br>5, 8, 10 | |
| 7 | 27.7 | 2.66 m α<br>2.12 m β | 6, 8, 9 | |
| 8 | 134.0 | --- | | |
| 9 | 152.5 | --- | | |
| 10 | 37.8 | --- | | |
| 11 | 123.4 | 6.80 d (10.0) | [c]--- | |
| 12 | 129.0 | 7.85 d (10.0) | [c]--- | |
| 13 | 126.8 | --- | | |
| 14 | 145.0 | --- | | |
| 15 | 23.0 | 3.15 m<br>2.92 m | 8, 13, 14 | |
| 16 | 14.7 | 1.29 t (7.4) | 14, 15 | |
| 17 | 170.1 | --- | | |
| 18 | 26.8 | 1.05 s | 3, 4, 5, 19 | 6 |
| 19 | 20.9 | 0.97 s | 3, 5, 18 | |
| 20 | 24.2 | 0.95 s | 1, 5, 9, 10 | 1β |

[a] $^{1}$H chemical shift of HSQC correlation peaks; [b] important correlations only and each NOE is listed only once; [c] very weak HMBC corrections and NMR data for C-11 and C12 were assigned via comparison of NMR data with compound **21a.**

**Tab. S11: NMR data of compound 35 ($C_6D_6$).**

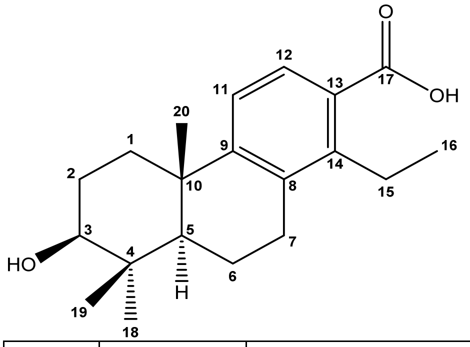

| Pos. | $^{13}C$: δ [ppm] | $^{1}H$: δ [ppm] m (J[Hz]) | selected HMBC (H → C) | NOE[b] |
|---|---|---|---|---|
| 1 | 37.10 | 1.901 dt-like (13.0, 3.6) β<br>1.174 α | --- | 2,11 |
| 2 | 28.107 | 1.492[a] m | --- | |
| 3 | 77.82 | 2.956[a] α | 19 | 18 |
| 4 | 39.24 | --- | | |
| 5 | 48.67 | 0.969[a] | --- | |
| 6 | 18.84 | 1.649m α<br>1.459[a] β | 8 | |
| 7 | 28.18 | 2.722 dd (16.9, 6.4)<br>2.451 m | 8 | |
| 8 | 134.40 | --- | | |
| 9 | 154.36 | --- | | |
| 10 | ---[c] | --- | | |
| 11 | 122.31 | 6.945 d(8.4) | 8, 13 | |
| 12 | 128.80 | 7.897 d(8.4) | --- | |
| 13 | 126.34 | --- | | |
| 14 | 144.87 | --- | | |
| 15 | 22.60 | 3.141[a]<br>2.948[a] | ---<br>8,14 | 7α,7β |
| 16 | 14.48 | 1.286 t (7.4) | 14,15 | 7α,7β |
| 17 | ---[c] | --- | | |
| 18 | 27.85 | 0.980 s | 3,4,5,19 | 6α |
| 19 | 15.27 | 0.81s | 5, 4, 18, 3 | |
| 20 | 24.4 | 0.99 s | 1,5,9 | 19 |

[a] $^{1}H$ chemical shift of HSQC correlation peaks; [b] each NOE is listed only once; COSY; [c] not detectable due to low concentration

**Tab. S12: NMR data of compound 21b ($C_6D_6$).**

| **Gene** | **Application** | **Direction** | **Sequence** |
|---|---|---|---|
| GenBank accession no. M60175 (*HvUBIQUITIN*) | qPCR | Forward | ACCCTCGCCGACTACAACAT |
| | | Reverse | CAGTAGTGGCGGTCGAAGTG |
| HORVU2Hr1G004540 (*HvKSL4*) | qPCR | Forward | GTTATCTCTGCGCTGCTGCC |
| | | Reverse | GAGGATCCTCCCATTTCTCAGC |
| | genotyping | Forward | GATCGGGAGGCCAGGATAAG |
| | | Reverse | CAGATCATAGTTGCGAATTAATGCAG |
| | truncate transit peptide | Forward | TTGGTCTCAACATAATGGCTTACGTTGAATCTAGACC |
| | | Reverse | TTGGTCTCAACAAACCAATTCGTTTTGAGACAAAATG |
| HORVU2Hr1G004620 (*HvCPS2*) | qPCR | Forward | GCGTCTGCAGCCCATGAGA |
| | | Reverse | GCGGTCTTCCTCCCTCTGC |
| | genotyping | Forward | GTTGCAGGTATGTTATTTATCAAAG |
| | | Reverse | CAATTTTTTTCTGTTCAAATTTGGTATG |
| | truncate transit peptide | Forward | TTGGTCTCAACATAATGGTTTTGTCCTCTAAATCTCCA |
| | | Reverse | TTGGTCTCAACAAGGGTTAACTTCGTAACCATGTTG |
| *BsTEF* (*2*) | qPCR | Forward | CGCCGTACCGGAAAGTCTG |
| | | Reverse | GGCGAAACGACCAAGAGGA |
| *hygromycin* | genotyping | Forward | GCGATTGCTGATCCCCATGT |
| | | Reverse | GGCGTCGGTTTCCACTATCG |

**Tab. S13: Primers used in this study.**

### References

1. S. Wawra *et al.*, The fungal-specific β-glucan-binding lectin FGB1 alters cell-wall composition and suppresses glucan-triggered immunity in plants. *Nature Communications* **7**, 13188 (2016).
2. D. Sarkar *et al.*, The inconspicuous gatekeeper: endophytic Serendipita vermifera acts as extended plant protection barrier in the rhizosphere. *New Phytologist* **224**, 886-901 (2019).
3. S. Kumar, G. Stecher, M. Li, C. Knyaz, K. Tamura, MEGA X: Molecular Evolutionary Genetics Analysis across Computing Platforms. *Mol Biol Evol* **35**, 1547-1549 (2018).
4. J. D. Thompson, D. G. Higgins, T. J. Gibson, CLUSTAL W: improving the sensitivity of progressive multiple sequence alignment through sequence weighting, position-specific gap penalties and weight matrix choice. *Nucleic Acids Res* **22**, 4673-4680 (1994).
5. K. J. Livak, T. D. Schmittgen, Analysis of relative gene expression data using real-time quantitative PCR and the 2(T)(-Delta Delta C) method. *Methods* **25**, 402-408 (2001).
6. S. Deshmukh *et al.*, The root endophytic fungus Piriformospora indica requires host cell death for proliferation during mutualistic symbiosis with barley. *Proc. Natl. Acad. Sci. U. S. A.* **103**, 18450-18457 (2006).
7. U. Scheler *et al.*, Elucidation of the biosynthesis of carnosic acid and its reconstitution in yeast. *Nat Commun* **7**, 12942 (2016).
8. H. Yadav *et al.*, Medicago TERPENE SYNTHASE 10 Is Involved in Defense Against an Oomycete Root Pathogen. *Plant Physiol* **180**, 1598-1613 (2019).
9. E. Weber, R. Gruetzner, S. Werner, C. Engler, S. Marillonnet, Assembly of Designer TAL Effectors by Golden Gate Cloning. *Plos One* **6**, (2011).
10. P. Urban, C. Mignotte, M. Kazmaier, F. Delorme, D. Pompon, Cloning, yeast expression, and characterization of the coupling of two distantly related Arabidopsis thaliana NADPH-cytochrome P450 reductases with P450 CYP73A5. *J Biol Chem* **272**, 19176-19186 (1997).
11. U. Lahrmann *et al.*, Mutualistic root endophytism is not associated with the reduction of saprotrophic traits and requires a noncompromised plant innate immunity. *New Phytol* **207**, 841-857 (2015).
12. M. Hilbert *et al.*, Indole derivative production by the root endophyte Piriformospora indica is not required for growth promotion but for biotrophic colonization of barley roots. *New Phytologist* **196**, 520-534 (2012).
13. D. G. Cooney, R. Emerson, Thermophilic fungi. An account of their biology, activities, and classification. *Thermophilic fungi. An account of their biology, activities, and classification.*, (1964).
14. C. A. Schneider, W. S. Rasband, K. W. Eliceiri, NIH Image to ImageJ: 25 years of image analysis. *Nature Methods* **9**, 671-675 (2012).
15. Y. Wang *et al.*, MCScanX: a toolkit for detection and evolutionary analysis of gene synteny and collinearity. *Nucleic Acids Res* **40**, e49 (2012).
16. H. Tang *et al.*, Unraveling ancient hexaploidy through multiply-aligned angiosperm gene maps. *Genome research* **18**, 1944-1954 (2008).
17. G. C. d. Silva, L. M. M. Valente, M. L. Patitucci, A. d. C. Pinto, N. L. d. Menezes, Diterpenóides com esqueleto cleistantano de Vellozia aff. carunculares Martius ex Seubert (Velloziaceae). *Química Nova* **24**, 619-625 (2001).

18. F. Chen, D. Tholl, J. Bohlmann, E. Pichersky, The family of terpene synthases in plants: a mid-size family of genes for specialized metabolism that is highly diversified throughout the kingdom. *The Plant Journal* **66**, 212-229 (2011).
19. T. U. Consortium, UniProt: the universal protein knowledgebase in 2021. *Nucleic Acids Research* **49**, D480-D489 (2020).
